# Epigenetic Memory of COVID-19 in Innate Immune Cells and Their Progenitors

**DOI:** 10.1101/2022.02.09.479588

**Authors:** Jin-Gyu Cheong, Arjun Ravishankar, Siddhartha Sharma, Christopher N. Parkhurst, Djamel Nehar-Belaid, Sai Ma, Lucinda Paddock, Benoit Fatou, Onur Karakaslar, Asa Thibodeau, Michael J. Bale, Vinay K. Kartha, Jim K Yee, Minh Yen Mays, Louise Leyre, Alexia Martinez de Paz, Andrew W. Daman, Sergio Alvarez Mullett, Lexi Robbins, Elyse LaFond, Karissa Weidman, Sabrina Racine-Brzostek, He S. Yang, David Price, Brad Jones, Edward J. Schenck, Robert J. Kaner, Amy Chadburn, Zhen Zhao, Hanno Steen, Virginia Pascual, Jason Buenrostro, Rachel E. Niec, Lindsay Lief, Duygu Ucar, Steven Z. Josefowicz

## Abstract

Severe coronavirus disease 2019 (COVID-19) is characterized by systemic inflammation and can result in protracted symptoms. Robust systemic inflammation may trigger persistent changes in hematopoietic cells and innate immune memory through epigenetic mechanisms. We reveal that rare circulating hematopoietic stem and progenitor cells (HSPC), enriched from human blood, match the diversity of HSPC in bone marrow, enabling investigation of hematopoiesis and HSPC epigenomics. Following COVID-19, HSPC retain epigenomic alterations that are conveyed, through differentiation, to progeny innate immune cells. Epigenomic changes vary with disease severity, persist for months to a year, and are associated with increased myeloid cell differentiation and inflammatory or antiviral programs. Epigenetic reprogramming of HSPC may underly altered immune function following infection and be broadly relevant, especially for millions of COVID-19 survivors.

**One Sentence Summary:** Transcriptomic and epigenomic analysis of blood reveal sustained changes in hematopoiesis and innate immunity after COVID-19.

**Graphical Abstract:** 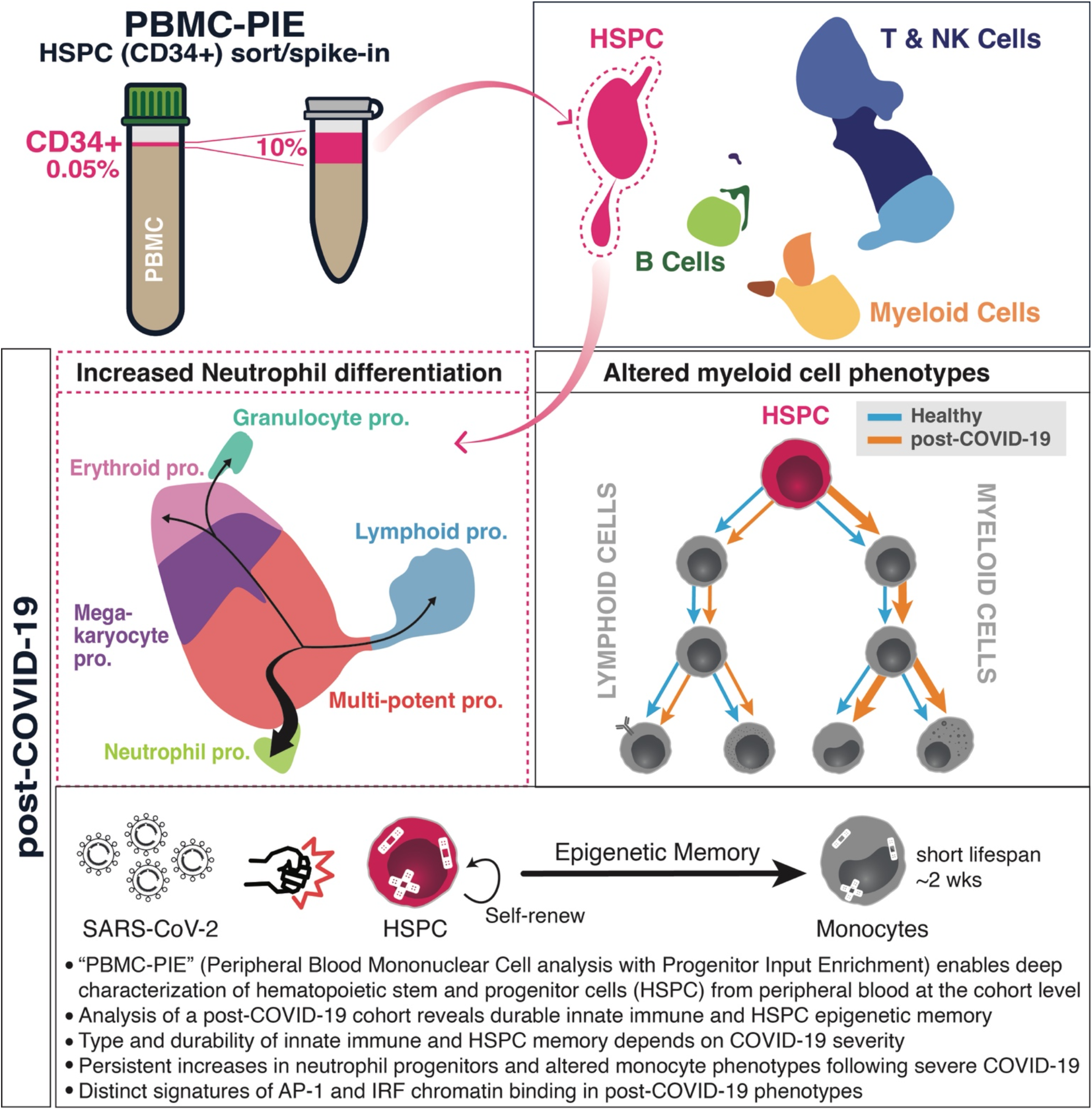

## Main Text

Coronavirus disease 2019 (COVID-19), the illness caused by infection with severe acute respiratory syndrome coronavirus 2 (SARS-CoV2), is characterized by a broad range of symptoms and severity and can result in a protracted course. Delayed adaptive immune and interferon (IFN) responses together with robust innate immune cell activity feature prominently in acute severe COVID-19 (*1*–*10*). However, the long-term effects of COVID-19 on the immune system are unclear. Durable changes in the immune system following COVID-19 could influence subsequent immune responses to pathogens, vaccines, or even contribute to long-term clinical symptoms, i.e., post-acute sequelae of SARS-CoV-2 infection (PASC) (*11*–*13*) and COVID-19-associated multisystem inflammatory syndrome in adults (MIS-A) and children (MIS-C) (*14*–*18*). Despite clinical observations of long-term sequelae, the nature of persistent molecular and cellular changes following COVID-19 are poorly understood.

Recent studies have established that innate immune cells and their progenitors can maintain durable epigenetic memory of previous infectious or inflammatory encounters, thereby altering innate immune equilibrium and responses to subsequent challenges (*19*). This innate immune memory, also termed trained immunity, has been attributed largely to persistent chromatin alterations that modify the type and scope of responsiveness of the cells that harbor them, including long-lived innate immune cells (*20*), epithelial stem cells (*21*), and self-renewing hematopoietic progenitors (*22*) and their mature progeny cells (*23*–*28*). While innate immune memory phenotypes have been well-studied with *in vivo* mouse models, the breadth, relevance, and molecular features of such phenotypes in humans have been more elusive.

A paradox of innate immune memory has been that many of the innate immune cells that retain durable alterations are themselves short-lived (*29*). Several recent studies in mice have provided a partial explanation of this paradox, revealing that hematopoietic stem and progenitor cells (HSPC) can be durably altered and epigenetically reprogrammed to persistently convey inflammatory memory to mature progeny cells with altered phenotypes, sometimes termed central trained immunity (*19*, *20*, *22*, *26*, *30*). Molecular features of such phenotypes in humans have focused on innate immune and hematopoietic progenitor cell alterations following administration of the tuberculosis vaccine Bacillus Calmette-Guérin (BCG) (*23*–*28*, *31*), an area of intrigue since protection to heterologous (non-tuberculous) infections post-BCG vaccination were historically observed (*32*–*34*). The complexities of studying human HSPC, especially in the context of large, cohort-level infectious disease studies, have limited our understanding of this type of hematopoietic progenitor cell-based memory.

HSPC are long-lived self-renewing precursors to diverse mature immune cells. Thus, HSPC are endowed with the unique potential to serve as reservoirs of post-inflammation epigenetic memory and retain programs that drive altered hematopoiesis and phenotypes in innate immune cell progeny. Based on these characteristics and on the inflammatory features of COVID-19, we hypothesized that exposure of HSPC to inflammatory signaling events during COVID-19 may result in epigenetic memory and persisting altered phenotypes following COVID-19. This may be especially true following severe COVID-19, which is characterized by sustained fever, elevated inflammatory cytokines and chemokines, immune complement activity, and tissue damage (*35*–*41*). Understanding the long-lasting effects of COVID-19-associated inflammation on hematopoiesis and innate immune memory is relevant for the hundreds of millions of people worldwide who have recovered from COVID-19 and especially the millions that still experience COVID-19 associated symptoms. The long-term clinical sequelae of COVID-19 (“long-haulers,” “long-COVID,” post-acute sequelae of SARS-CoV-2 infection or PASC) (*11*, *13*), particularly among those admitted to the ICU (*42*), suggests that persistent changes or alterations in immune activity may play a role. Furthermore, the etiology of incomplete recovery after critical illnesses is poorly understood, and it is possible that lasting alterations in immune cells and their inflammatory response programs contribute to incomplete recovery and post-ICU syndrome (PICS) (*43*).

In this study, we identify epigenetic innate immune memory that results from SARS-CoV-2 infection by characterizing the molecular features of the post-infection period (2 to 12 months after the start of mild and severe COVID-19). We focus on comprehensive analyses of alterations in chromatin and transcription at the single-cell level in monocytes and their HSPC progenitors. In order to study HSPC in-depth, at single-cell resolution, we developed a new workflow to enrich and profile rare hematopoietic stem and progenitor cells from peripheral blood, termed Peripheral Blood Mononuclear Cell analysis with Progenitor Input Enrichment (PBMC-PIE). PBMC-PIE addresses the challenge in access to HSPC (generally acquired through bone marrow aspirate), a limitation that has severely hindered the study of altered hematopoiesis and innate immune memory in the context of human infection and disease. We paired PBMC-PIE with single-nuclei combined RNA-sequencing (RNA-seq) and assay for transposase-accessible chromatin (snRNA/ATAC-seq). This approach revealed a high-resolution transcriptomic and chromatin accessibility map of diverse HSPC subsets and PBMC in a unique cohort of convalescent COVID-19 study participants, including mild (non-hospitalized) and severe (S, requiring intensive care unit (ICU) admission) convalescent COVID-19 patients, following acute SARS-CoV-2 infection from months (2-4mo, early convalescence) to a year (4-12mo, late convalescence). We compared these cohorts to healthy participants and also to participants recovering from non-COVID-19 critical illness (nonCoV; patients requiring ICU admission) in order to identify features common among patients recovering from critical illness and also unique to patients recovering from severe COVID-19. To complement this deep characterization of HSPC and PBMC from convalescent COVID-19 study participants, we also profiled (i) plasma proteins, by liquid chromatography-mass spectrometry (LC-MS); (ii) SARS-CoV-2 antibody quantity and quality; (iii) circulating cytokines/chemokines; and (iv) progenitor and PBMC populations, by flow cytometry. Thus, the study was designed to understand the molecular features of recovery for a range of SARS-CoV-2 infection severity over a period of months to one year, with the inclusion of uninfected individuals and also non-COVID-19 critical illness as an important control group.

We reveal the persistence of epigenetic and transcription programs in HSPC and monocytes following severe disease that are indicative of an altered innate immune responsiveness. These included durable epigenetic memory linked to inflammatory programs, neutrophil differentiation, and monocyte phenotypes. Further, following mild COVID-19, we uncover a months-long, active, interferon regulatory factor/signal transducer and activator of transcription (IRF/STAT)-driven, anti-viral signature in monocytes (with matching programs in HSPC progenitors), indicative of a state of antiviral vigilance (*44*). This highlights the potential for acute viral infection to drive a durable program of HSPC origin that is conveyed through to progeny monocytes to mediate a heightened anti-viral response program, with potential implications for heterologous protection and resilience to seasonal infections.

## Results

### Cohort Characterization, Plasma Analysis, and HSPC Enrichment

We studied molecular features of immune cells from convalescent COVID-19 patients with varying disease severities (mild and severe), compared with those of i) healthy participants (n=47) and ii) study participants recovering from non-COVID-19-related critical illness (due to either infectious or non-infectious non-COVID-19 disease, “nonCoV,” n=25). Our convalescent COVID-19 cohort included both mild (“M,” non-hospitalized, WHO score 1-2; n=25) and severe (“S,” WHO score 6-7, requiring ICU admission; n=87) study participants. To study the durability of molecular features of immune cells, we collected samples between 2 to 4 months following the onset of symptoms (S:2-4mo and M:2-4mo), termed “early convalescence,” or between 4 to 12 months post-acute (S:4-12mo and M:4-12mo), termed “late convalescence.” Healthy donors (symptom-free and seronegative) were enrolled with consideration to age, sex, and comorbidity to match COVID-19 groups (demographic and clinical information, table S1, data S1). Since study participants were enrolled at Weill Cornell New York Presbyterian Hospital during the initial wave of infections in New York City (spring through winter of 2020), their infections were likely wildtype/non-variant SARS-CoV-2, and none were administered COVID-19 vaccines. To comprehensively study immune cells of these cohorts, including HSPC, we established a multimodal assay and analysis workflow for deep characterization of cellular and molecular features of immune cells in the post-infection period and across a range of disease cohorts (Fig. 1A, fig. S2A).

**Fig. 1.**
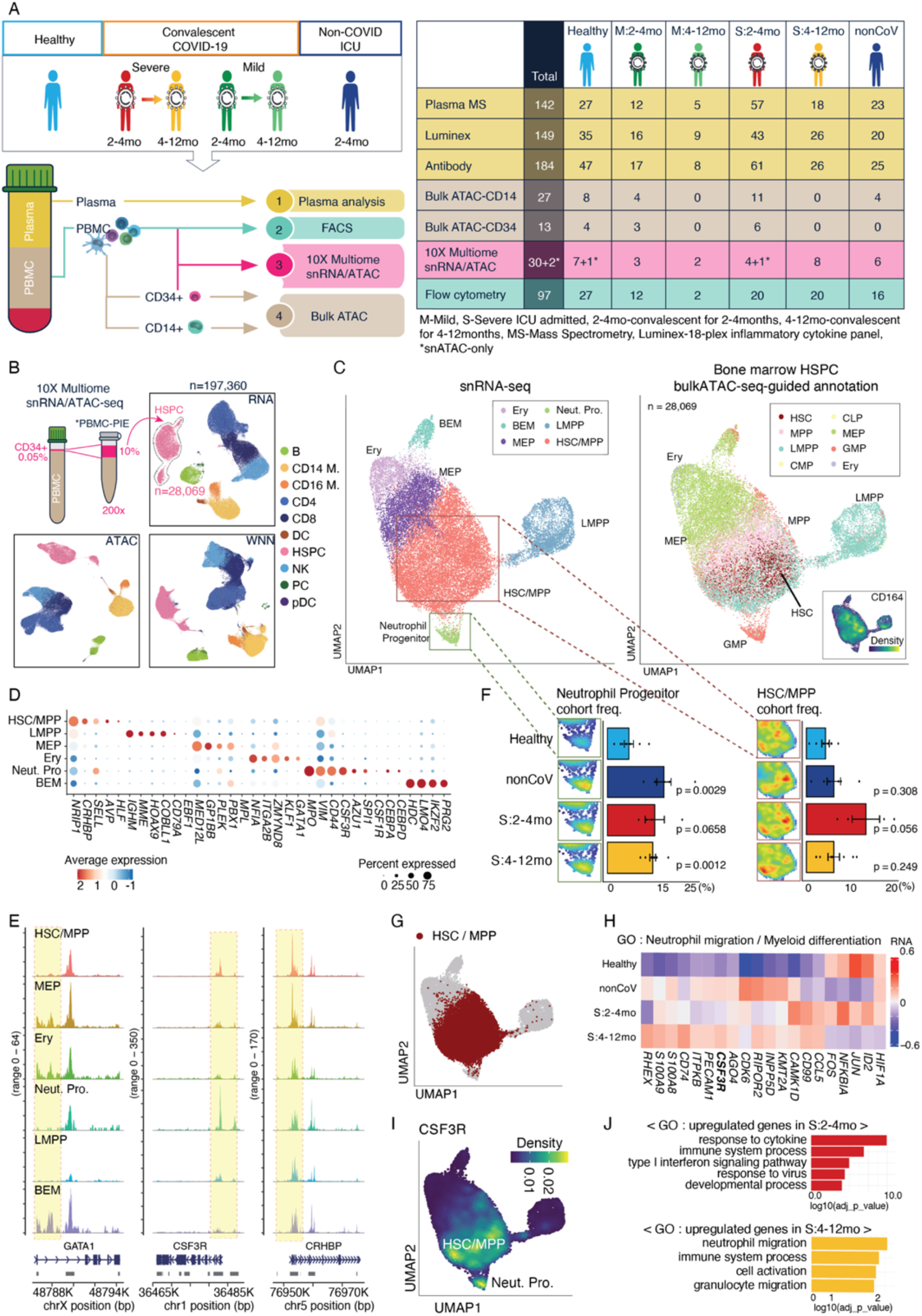
Deep characterization of hematopoietic stem and progenitor cells (HSPC) from peripheral blood reveals sustained alterations in hematopoiesis post-COVID-19. **(A)** Study overview depicting cohort design, workflow, and summary table of samples used for different analyses. Cohort definitions and abbreviations listed on bottom right. **(B)** Schema for analysis of peripheral blood mononuclear cells with progenitor input enrichment (PBMC-PIE) followed by single-nuclei combined RNA/ATAC (snRNA/ATAC) analysis and UMAP visualizations for snRNA-seq (RNA), snATAC-seq (ATAC), and weighted nearest neighbor (WNN). **(C)** UMAP visualizations of HSPC with annotations based on expression of manually curated marker genes for HSPC subsets (left) and with annotation guided by ATAC-seq data from sorted human bone marrow HSPC subsets (right) (*48*). Inset on right shows expression density for *CD164*, a marker gene of early HSC. **(D)** Dot plot of expression of select markers in each HSPC subcluster. Colors and sizes of dots indicate the average expression of and percent of cells expressing the indicated gene, respectively, in each HSPC subcluster. **(E)** Examples of chromatin accessibility measured by ATAC-seq at HSPC subcluster-marker gene loci: *GATA1* (Ery, MEP); CSF3R (neutrophil progenitors); *CRHBP* (HSC/MPP). **(F)** Bar plot showing percent of neutrophil progenitors (left) and HSCMPP (right) among total HSPC in individuals within the indicated cohorts, Adjacent are paired UMAP density plots of neutrophil progenitors (left) and HSC/MPP (right) for each cohort. (*p<0.05, t-test, Healthy group as a reference) **(G)** HSC/MPP cells highlighted in red among HSPC cells (grey) on UMAP space. **(H)** Expression of genes associated with neutrophil migration (GO:1990266) and myeloid differentiation (GO:0030099) in the HSC/MPP cluster. **(I)** *CSF1R* expression in HSPC projected on UMAP space. **(J)** GO analysis of differentially upregulated genes in HSCMPP of S:2-4mo and S:4-12mo compared to the healthy cohort. *B cells (B), CD14+ monocytes (CD14 M.), CD16+ monocytes (CD16 M.), CD4+ T cells (CD4), CD8+ T cells (CD8), dendritic cell (DC), hematopoietic stem and progenitor cells (HSPC), natural killer cells (NK), plasma B cells (PC), plasmacytoid dendritic cells (pDC), erythroid progenitor cells (Ery), neutrophil progenitor cells (Neut. Pro.), basophil-eosinophil-mast cell progenitor cells (BEM), lymphoid-primed multipotent progenitor cells (LMPP), megakaryocyte-erythroid progenitor cells (MEP), hematopoietic stem cells/multipotent progenitor cells (HSC/MPP), common lymphoid progenitor cells (CLP), granulocyte-macrophage progenitor cells (GMP), common myeloid progenitor cells (CMP)

Plasma analysis included SARS-CoV-2 receptor-binding domain (RBD) spike-specific antibody levels, surrogate neutralizing antibody activities, and antibody avidity; 18-plex cytokine and chemokine quantification (Luminex); and unbiased LC/MS-based mapping of plasma proteins. These data and their analyses uncovered plasma factors that persist for months post-acute SARS-CoV-2 infection (fig. S1B-C), and also expected antibody dynamics, with increased quality and durability of antibody responses in those with severe disease (fig. S1A). No sustained alterations in plasma factors (i.e., proteins or cytokines/chemokines from LC/MS and Luminex analysis) were observed in mild post-COVID-19 (M:2-4mo and M:4-12mo, fig. S1B-C) compared to healthy study participants. In contrast, we observed a persistent and distinctive active plasma response featuring inflammatory cytokines (IL-6, IL-8) (fig. S1C), immune complement (C1QA, C1R, C3, C4A), and vascular response factors (SAA1, ORM1, LBP, fibrinogen-FGG/FGA, LRG1) in early convalescent severe COVID-19 study participants (S:2-4mo) (fig. S1B). Levels of acute-phase proteins and platelet factors were elevated in both S:2-4mo and nonCoV, compared to healthy study participants (fig. S1B). These data establish the presence of a months-long activation of immune complement and vascular response factors that could contribute to post-COVID-19 sequelae (fig. S1B-C and Supplementary text). Importantly, in late convalescence following severe COVID-19 (S:4-12mo), there were no significant alterations in plasma factors by LC/MS compared to healthy donors, indicating a full resolution of the active plasma response and overt inflammation at the protein level (fig. S1B). This suggests that any persistent changes in cellular composition or phenotypes in late convalescence of study participants with severe COVID-19 are independent of an ongoing and active inflammatory program and hence point to epigenetic mechanisms behind prolonged effects of COVID-19.

To overcome the challenges of obtaining HSPC to study hematopoiesis and central trained immunity in human cohorts, we developed an experimental workflow to enrich for HSPC, enabling their in-depth study paired with the mature immune cell populations from the same donor, termed Peripheral Blood Mononuclear Cell analysis with Progenitor Input Enrichment (PBMC-PIE). Rare circulating CD34^+^ HSPC (~0.05% of PBMCs) were enriched by sequential antibody-conjugated bead-based enrichment and FACS sorting from PBMCs and “spiked” into sorted PBMCs from the same donor at an approximate ratio of 1:10, a 200-fold enrichment of their original frequency. This approach is particularly suitable for singlecell profiling; we profiled PBMC-PIE samples from a subset of our cohort (n=32) using single-cell transcriptomic and epigenomic assays. These genomic profiles were complemented by additional bulk ATAC-seq profiling of sorted CD14^+^ monocytes (total of 27, all groups) and peripheral CD34^+^ HSPC (total of 13, all groups) to increase our detection power for differential analyses (Fig. 1A, right; data S1-2).

### Representation of diverse blood progenitors among peripheral HSPC

PBMC-PIE combined with snRNA/ATAC-seq (10X Genomics Chromium Multiome) enabled the study of gene regulatory programs across clinical cohorts and between HSPC progenitors and their mature progeny cells among PBMC. Combined analyses of ATAC- and RNA-seq using weighted nearest neighbor (WNN) resulted in 10 primary clusters reflecting the major PBMC populations. Previous studies (*45*–*47*) have reported lymphopenia and increases in neutrophils, eosinophils, proliferating lymphocytes, and plasma B cells during acute severe COVID-19. These differences appear to mostly resolve in late convalescence. However, increased frequencies of circulating B cells persist to late convalescence; reduced frequencies of circulating plasmacytoid dendritic cells (pDC) were found in early convalescence; and frequencies of circulating CD4^+^ T cells were decreased in early convalescence (fig. S2D). While we focus analyses on the HSPC and myeloid populations for insight into innate immune memory phenotypes, the complete snRNA/ATAC dataset serves as a comprehensive single-cell atlas for understanding post-COVID and post-ICU epigenomic and transcriptomic alterations in diverse immune cell types and their progenitors.

To assess the suitability of peripheral blood CD34^+^ HSPC from PBMC-PIE as surrogates for bone marrow HSPC, we annotated the 28,069 HSPC from our cohort using (i) bulk ATAC-seq data from FACS-sorted bone marrow HSPC subsets from our previous studies (Fig. 1C, right panel; fig. S3A) (*48*); and (ii) scRNA-seq data from bone marrow mononuclear cells (BMMC) (fig. S3B) (*49*). HSPC subcluster annotations from both bone marrow datasets (bulk ATAC-seq and scRNA-seq) were projected onto single-cell peripheral HSPC data resulting in two alternative annotations of our HSPC subsets. The two projections have good agreement and reveal the diversity of peripheral HSPC and their similarity to BMMC HSPC. Importantly, early hematopoietic stem cell (HSC) and multipotent progenitor (MPP) subsets were prominently represented among peripheral HSPC; *CD164*, a gene suggested as an early hematopoietic stem cell marker (*50*), was highly expressed in the HSC subcluster as defined by bulk-ATACseq-guided annotation (Fig. 1C, right, inset; fig. S3C).

We then used an unbiased workflow to cluster our CD34^+^ HSPC and identify differentially expressed genes among the 12 resulting subclusters (fig. S3A). Manual curation of subclusters using marker genes from the literature (*48*–*50*) led us to merge select subclusters resulting in six HSPC subsets with both well-characterized and novel marker genes (Fig. 1C-D; fig. S4A, right), that manifests all major bone marrow HSPC subsets. These HSPC subclusters were defined as hematopoietic stem cells and multipotent progenitors (HSC/MPP), lymphoid-primed MPP (LMPP), megakaryocyte-erythroid progenitors (MEP), erythroid progenitors (Ery), neutrophil progenitors (Neut. Pro.), and basophil-eosinophil-mast cell progenitors (BEM) (Fig. 1C-D; fig. S4A, right). Importantly, projection of these annotations onto snATAC-based clustering and UMAP visualization also generated distinct HSPC subclusters (fig. S3A) and indicated that these HSPC subsets feature both distinct expression and epigenetic profiles. Recent single-cell analysis has revealed intriguing heterogeneity among granulocyte-monocyte progenitors with distinguishing features between BEM and neutrophil progenitors (*50*). In our dataset, these BEM and neutrophil progenitor populations were clearly distinguishable, forming distinct subclusters and annotated by characteristic marker genes. For neutrophil progenitors, these included *MPO*, *VIM*, *AZU1*, and prominent neutrophil differentiation receptor, *CSF3R*, and for BEM, markers included *HDC*, *LMO4*, *PRG2*, and *IKZF2* (Fig. 1D; fig. S4A, right). Analysis of snATAC-seq revealed chromatin accessibility profiles with selective accessibility at lineage-defining loci, for example, *GATA1* (Ery), *CSF3R* (neutrophil progenitors), and *CRHBP* (HSC/MPP) (Fig. 1E).

In summary, PBMC-PIE enabled us to perform single-cell profiling of HSPC from convalescent COVID-19, healthy donor, and non-COVID19 ICU participants, with an unprecedented level of resolution and detail (with ~30,000 annotated HSPC), and with combined chromatin and expression analysis. Further, our results establish that although circulating HSPC are rare, they recapitulate the diverse HSPC subclusters conventionally studied in the bone marrow niche.

### Altered hematopoiesis following COVID-19

We next explored if there were alterations in HSPC subset frequencies across COVID-19 and control study participants. We observed increased frequencies and population shifts in HSC/MPP following severe disease, in general, both in S:2-4mo and nonCoV (Fig. 1F, right; fig. S3G). These changes were more significant in the post-COVID-19 cohort and pointed to an emergency hematopoiesis phenotype (i.e., a broad activation of HSPC subsets induced by inflammation to replenish leukocytes and mobilize an effective response) that was consistent with the still-active pro-inflammatory plasma response (fig. S1B-C). Flow cytometry analysis revealed similar increases in LT-HSC subsets (Lin^-^CD34^+^CD38^-^CD45RA^-^CD90^+^CD49f^+^) (fig. S3E), providing complementary validation of this emergency hematopoiesis phenotype. We also observed increased frequencies of neutrophil progenitors following critical illness (S:2-4mo and nonCoV), consistent with ongoing emergency granulopoiesis that could be a consequence of maintained elevated plasma cytokines and acute-phase proteins, which promote neutrophil production. Notably, this increase in neutrophil progenitors (but not BEM) persisted into late convalescence following severe COVID-19 (S:4-12mo) (Fig. 1F, left; fig. S3F-G), despite resolution of the active plasma response (fig. S1B-C), suggesting a durable alteration in HSPC.

To investigate features that may drive a persistent bias towards neutrophil differentiation in precursor cells post-COVID-19, we identified genes within the HSC/MPP subcluster that were differentially expressed between healthy and afflicted cohorts (Fig. 1H, J). Gene ontology (GO) analysis of upregulated genes in severe COVID-19 cohorts (both early and late convalescence) included terms linked to granulopoiesis. In early convalescence (S:2-4mo), a still-active response to inflammation and an inflammatory response in HSPC was detected (GO: response to cytokine/virus, type I interferon signaling, and immune system process). In late convalescence (S:4-12mo) a granulocyte/neutrophil program was detected (GO: neutrophil/granulocyte migration) (Fig. 1J). Among differentially expressed neutrophil differentiation-related genes in this precursor population were *S100A8/A9* and *CSF3R* (Fig. 1H). *CSF3R*, while a marker for neutrophil progenitors, is also expressed early in HSC/MPP subclusters (Fig. 1H-I), and its signaling in response to CSF3 (G-CSF) is necessary and sufficient to drive neutrophil differentiation (*51*). These data suggest that persistent and increased *CSF3R* expression in HSC/MPP post-COVID-19 may contribute to augmented neutrophil differentiation. Beyond the identification of post-COVID-19 features, our large (32 subjects, 28,069 cells) single-cell dataset of diverse HSPC subsets and their annotations demonstrate that PBMC-PIE provides the opportunity to study alterations in hematopoiesis and HSPC transcription and epigenetic signatures in the context of health and disease states using peripheral blood specimens.

### Linked chromatin-expression analysis and altered myelopoiesis following COVID-19

PBMC-PIE also provides a means of studying changes in HSPC phenotypes that may be durably propagated by stem cell self-renewal and conveyed to progeny, including mature innate immune cells, through differentiation. To further define post-COVID-19 HSPC phenotypes, we cumulatively compared gene expression programs of HSPC across clinical groups (Fig. 2A-B, fig. S4B-D). This revealed that study participants with the mild disease had few, but notable, changes during early convalescence (M:2-4mo), including reduced expression of AP-1 complex members (*FOS* and *JUN*) and histone variant H3.3B (*H3F3B*) and increased expression of interferon-stimulated genes (*IFITM3*, and *IFI44L*) and antigen presentation related genes (*HLA-C*, *HLA-A*, *CD74*) (Fig. 2A, fig. S4B-D). In contrast, an ongoing inflammatory response was apparent in early convalescent severe (S:2-4mo) and nonCoV cohorts with the upregulation of pro-inflammatory alarmins *S100A8*, *S100A9*, and other important immune signaling and transcription-regulating genes, including *USP15*, *STAT1*, *IRF1*, and *TRAF3* (Fig. 2A, fig. S4B-D). Severe COVID-19 study participants maintained an altered HSPC program into late convalescence (S:4-12mo), including increased expression of *CSF3R*, antigen presentation-related factors, and S100 genes, and decreased expression of *H3F3B*, *FOS*, and *JUN* (Fig. 2A, fig. S4B-D). The loci of these transcriptionally upregulated genes were often characterized by increased chromatin accessibility. For example, paralleling the observed upregulation of the *S100A8* gene, chromatin accessibility around this locus also increased post-COVID-19 (Fig. 2D-E). *S100A8* is an endogenous ligand for Toll-like receptor 4 (TLR4), an important proinflammatory mediator, and is upregulated in acute COVID-19 (*52*). Cumulatively, these data suggest that COVID-19, and particularly severe disease, induce gene expression changes in HSPC that last up to a year.

**Fig. 2.**
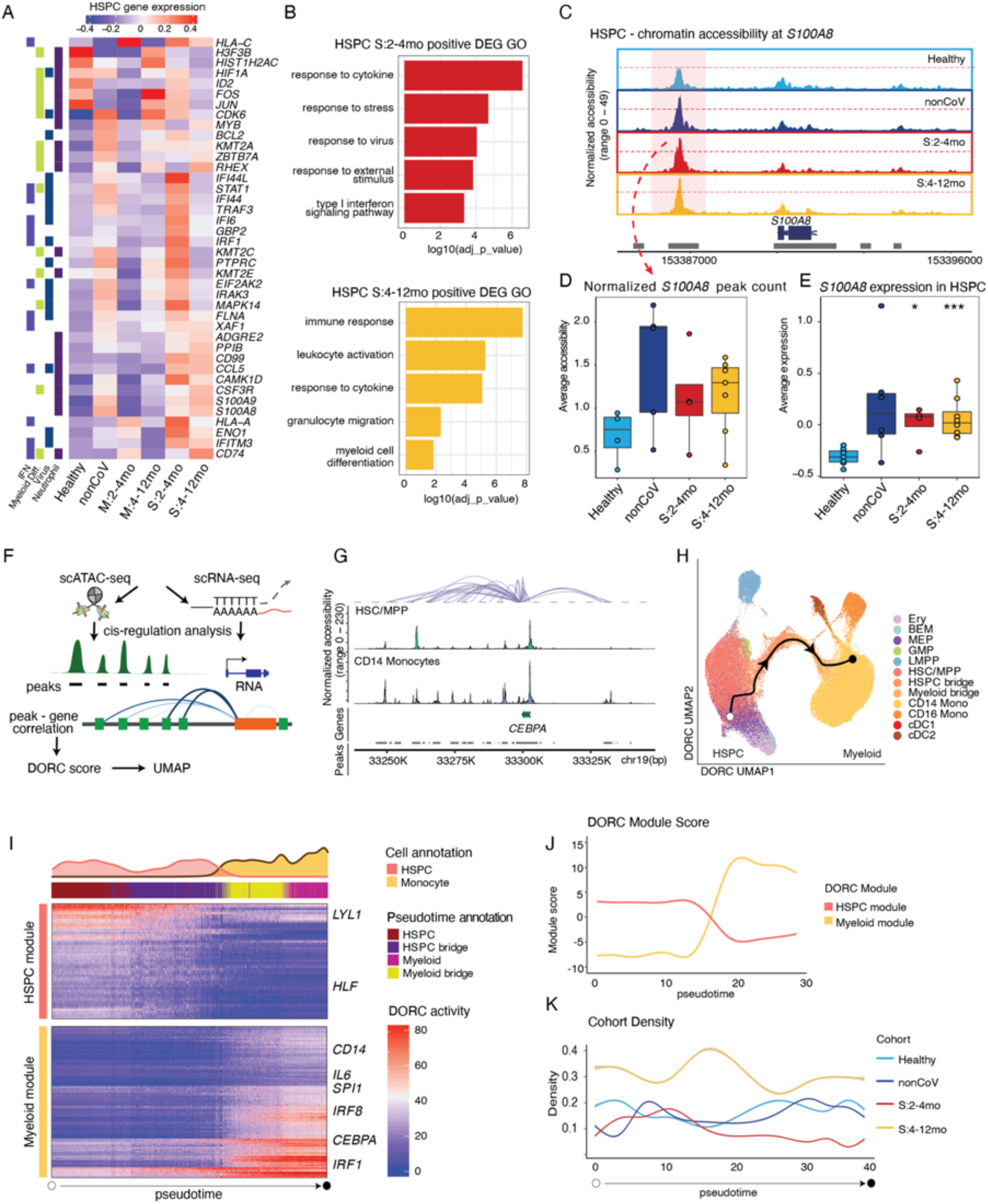
Durable transcriptomic and chromatin accessibility signatures and altered myeloid differention post-COVID-19. **(A)** Expression level of selected genes differentially expressed in HSPC following severe COVID-19 compared to healthy individuals (S:2-4mo and S:4-12mo compared to Healthy) across all cohorts. The genes are categorized as bar annotations (left). **(B)** GO analysis of differentially upregulated genes in HSPC of early and late convalescent severe (S:2-4mo and S:4-12mo) compared to Healthy. **(C)** Genome track demonstrating chromatin accessibility around *S100A8* loci of HSPC across clinical cohorts. **(D)** Box plot showing average gene expression of S100A8 in each sample across clinical groups. Each clinical group was compared to Healthy (t-test, ns). **(E)** Average normalized chromatin accessibility in an *S100A8*-proximal peak shown by reads per kilobase per million (RPKM) for each sample compared across clinical groups (t-test, *p<0.05, ***p<0.001). **(F)** Schematic describing workflow for analyzing distal regulatory elements and gene expression using singlenuclei multiome data including domains of regulatory chromatin (DORC). ATAC-seq peaks were linked to putative target genes in cis, based on the co-variation of chromatin accessibility and gene expression levels across individual cells (*53*). Developmentally related HSPC and monocytes were then co-clustered based on DORC scores. **(G)** Genome track showing chromatin accessibility of CEBPA-associated DORC in HSC/MPP and CD14^+^ monocytes. Curved lines (loops) above indicate peaks with significant correlation (P<0.05). The grey bar (below gene track) represents peaks. **(H)** UMAP visualization of HSPC and myeloid cell types. The line shows a subset of trajectories with starting node chosen based on high expression of *CD164* (early HSPC) and ending node selected based on first CD14^+^ monocyte cluster after the bridge (Monocle 3 (*99*)) from which we obtained a subset pseudotime for downstream analyses. **(I)** Heatmap showing DORC activity on pseudotime to identify HSPC and myeloid modules. Annotations on top are colored by representation of cell types based on clusters in H (top) and based on pseudotime analysis (just below top). **(J)** Line graph showing changes of DORC module scores (see methods) across pseudotime for the HSPC module and myeloid module. **(K)** Line plot showing the cohort density across pseudotime. The cohort density is defined as the frequence of cells from the indicated cohort among the 50 neighboring cells on the UMAP plot normalized for total cells in the cohort (see methods).

To aid in the discovery of genes that are regulated by disease-associated epigenetic changes, we identified Domains of Regulatory Chromatin (DORC) (*53*). ATAC-seq peaks were linked to their putative target genes in cis, based on the co-variation of chromatin accessibility and gene expression levels across individual cells (*53*) (Fig. 2F). DORC represent regulators of the non-coding genome, and DORC genes are highly enriched for genes associated with developmental programs and lineage specification and depleted for housekeeping, metabolic, and cell-cycle associated genes (*53*). We detected 968 DORC from our data, where at least 15 cis-regulated peaks were associated to each DORC/gene (fig. S5A-B). Among prominent DORCs was one associated with the key monocyte differentiation factor, *CEBPA*, which was more accessible and more highly expressed in CD14^+^ monocytes compared to HSC/MPP (Fig. 2G). We took advantage of the fact that DORC-associated genes are enriched for those that regulate immune development and clustered developmentally related HSPC, monocytes, and B cells based on the DORC scores (fig. S5G). Single cells reliably clustered by cell type (Fig. 2H). Further, this DORC-based clustering revealed cells that are intermediate in a state between HSPC to mature immune cell subsets, which appeared as bridges between clusters (Fig. 2H, fig. S5G). The DORC activity of developmental driver genes *CEBPA* (myeloid) and *EBF1* (B cell) increased from HSPC to monocytes and B cells, respectively, across the cellular bridges, suggesting that DORC activity of single cells can capture differentiation trajectories of distinct lineages and associated marker genes (fig. S5G-I).

We then sought to further validate DORC-based co-clustering and trajectories and to study putative monocyte precursors and potential alterations in myelopoiesis post-COVID-19. We co-clustered HSPC and myeloid cells (CD14^+^ and CD16^+^ monocytes, DC, and pDC) using DORC scores and revealed putative trajectories. Among these, we focused on a continuous trajectory from HSPC to monocyte for further characterization to reveal putative monocyte precursors (Fig. 2H, fig. S5J-L). Two groups of genes defined this HSPC-monocyte trajectory based on their patterns of DORC activity: i) those with declining DORC activity across this pseudotime trajectory from HSPC to monocytes including marker genes for HSC such as *HLF* (*54*) and *LYL1* (*55*) (Fig. 2I, pink HPSC module; fig. S5D); and ii) those with increasing activity including important molecules involved in monocyte cell function such as *CD14*, *CEBPA*, *SPI1*, *IRF1*, *IRF8*, and *IL6* (Fig. 2I, yellow myeloid module; fig. S5D). An additional module of DORCs described lymphoid development genes that were repressed along the myeloid bridge as expected (fig. S5D). We considered that these annotations might reflect either developmental trajectories and/or inflammation-altered phenotypes of progenitor cells and monocytes. We, therefore, investigated normalized frequencies of cells along this pseudotime trajectory for post-COVID-19 and nonCoV cohorts. Consistent with HSPC gene expression analysis (Fig. 1H, J; Fig. 2A-B), we found that the late convalescent severe (S:4-12mo) cohort had more cells constituting the HSPC-myeloid trajectory compared to other clinical groups and healthy controls, suggesting increased myelopoiesis potential of these study participants’ HSPC (Fig. 2K; fig. S5M, N).

Thus, PBMC-PIE paired with combined snRNA/ATAC analysis enabled detailed study of HSPC and immune progeny cells from our cohort, including chromatin landscapes, gene expression programs, and developmental trajectories. Further, this approach uncovered durably altered myelopoiesis following severe COVID-19, with increased monocyte and neutrophil precursors among HSPC that persist into late convalescence (S:4-12mo).

### Post-COVID-19 memory of interferon or inflammatory signatures in monocytes and progenitors

We next investigated whether mature circulating monocytes bear phenotypic differences as a consequence of alterations in HSPC and myelopoiesis in COVID-19 convalescence. Altered monocyte phenotypes have been reported during acute COVID-19, including reduced interferon responses and increased inflammatory signatures in severe disease (*9*, *47*, *56*). We sought to examine if such characteristics persist in convalescence. Clustering of CD14^+^ monocytes (n=19,400 cells) based on gene expression identified four distinct subclusters (SC0 to SC3) and their associated marker genes (Fig. 3A-B, fig. S6A-C). SC2 (n=7,867) was defined by high expression of *NRIP1*, co-activator of NF-kB (*57*), *NEAT* or Virus Inducible NonCoding RNA (*VINC*), *NAMPT*, and other genes annotated as enriched in GO category “response to cytokine” (p = 4.79E-6), and was expanded specifically in both early convalescent severe (S:2-4mo) and nonCoV cohorts (Fig. 3B-C, left). Notably, this distinct cluster of inflammatory monocytes resolved in late convalescent severe (S:4-12mo) study participants (though partial inflammatory and migratory macrophage programs persisted (see below, Fig. 4). In contrast to SC2, SC3 exhibited a Type-1 IFN signature (IFN^high^), including genes such as *MX1*, *MX2*, *IFI44*, or *IFI44L* (annotated as “type I interferon signaling pathway,” p=2.77E-9 and “defense response to virus,” p=4.73E-22) and was specifically expanded within early convalescent mild study participants (M:2-4mo) (Fig. 3B-D, right). Mild disease-associated anti-viral SC3 monocytes also contracted in late convalescence (Fig. 3B, right). These observations are in line with previous studies associating acute mild cases of COVID-19 with strong activation of anti-viral programs, whereas severe cases were more linked to the induction of pro-inflammatory programs (*4*, *7*). Notably, our study reveals the longevity of these phenotypes during the convalescent period.

**Fig. 3.**
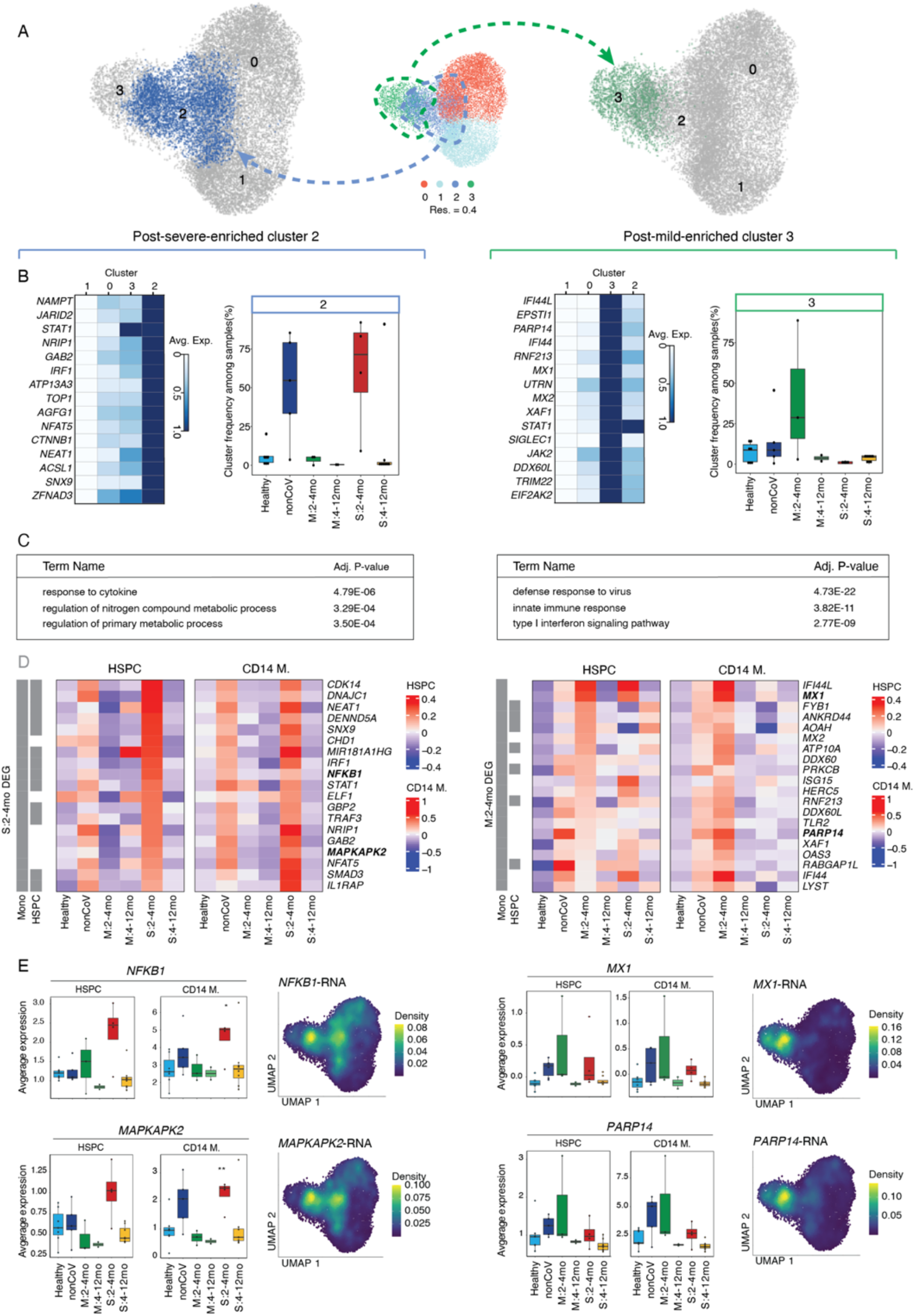
Anti-viral and inflammatory programs define months-long monocyte memory following severe disease. **(A)** UMAP visualization of CD14^+^ monocytes with sub-clusters. ub-cluster2 (SC2, post-severe-enriched) and sub-cluster3 (SC3, post-mild-enriched) are highlighted in blue and green, on the left and right, respectively. **(B)** Heatmap representing the scaled expression values of the top 15 genes defining SC2 (in blue) and SC3 (in green). **(C)** Top three enriched GO categories from the top SC2 and SC3 marker genes. **(D)** Heatmap showing expression of SC2 and SC3 marker genes across the clinical cohorts in both HSPC (left panels) and CD14^+^ monocytes (right panels). Bar annotations indicate which genes are significantly differentially expressed (P<0.05) compared to HD. **(E)** Box plots presenting gene expression of selected SC2- (*NKFB1* and *MAPKAPK2*) or SC3- (*MX1* and *PARP14*) specific genes in individual samples grouped by clinical cohorts for HSPC (left panel) and CD14^+^ monocytes (right panel). The expression level was projected on the UMAP. (*p<0.05, **P<0.01, t-test, Healthy group as a reference)

**Fig. 4.**
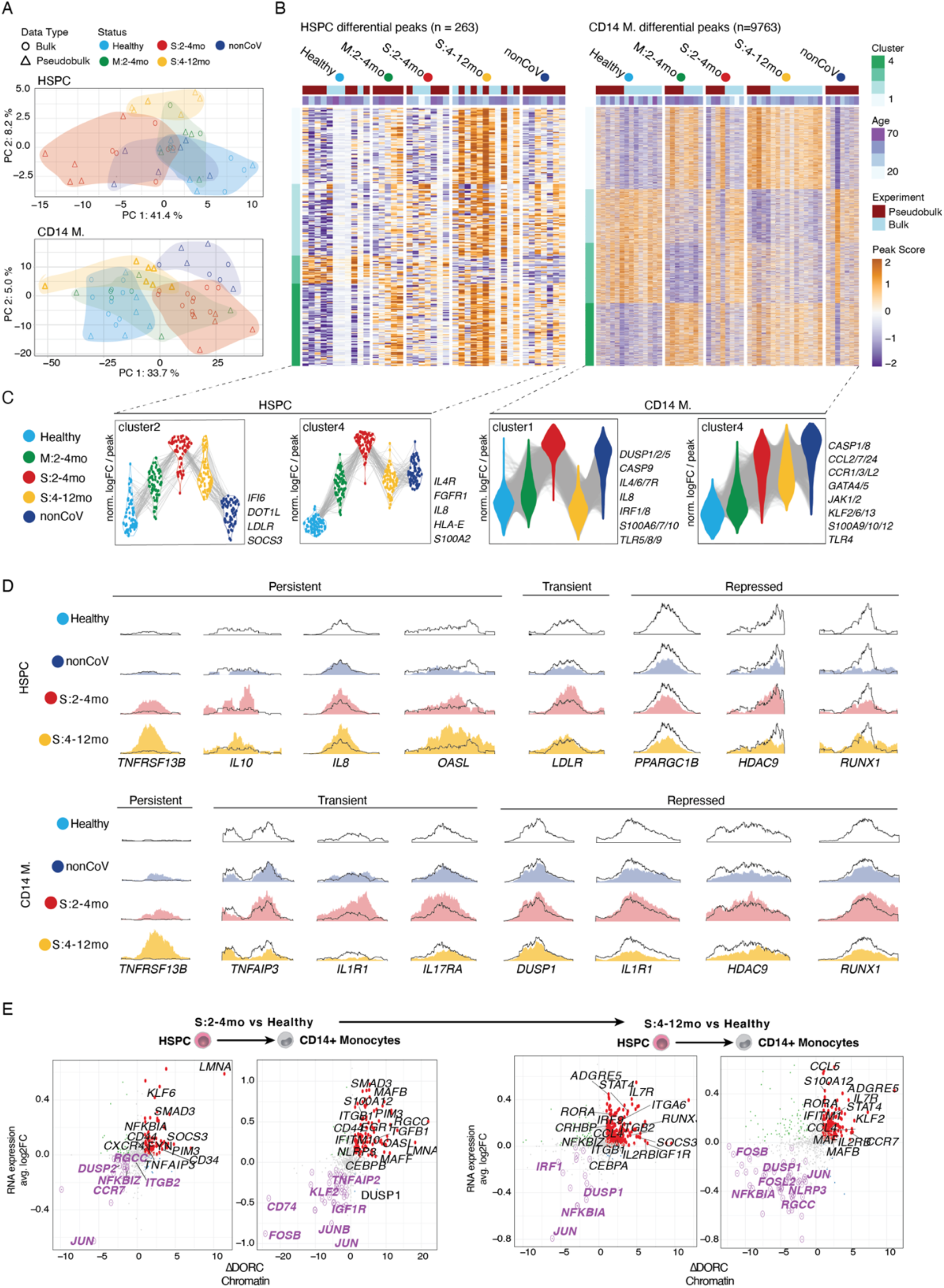
Durable epigenetic memory post-COVID-19 in HSPC and monocytes. **(A)** PCA of bulk ATAC-seq data of CD34^+^ HSPC and CD14^+^ monocytes (circles) plotted together with pseudobulk ATAC-seq data for both cell types from single-nuclei ATAC-seq data are included (triangles). **(B)** Heatmap of differentially accessible regions in HSPC (left) and CD14^+^ monocytes (right) clustered by pattern across groups. **(C)** Violin plots for peaks in select clusters of the heatmaps in (B), labeled with inflammatory genes as examples. **(D)** Genome tracks of individual chromatin accessible regions with persistent, transient or repressed changes across groups (Healthy, outline; S2-4mo, red; S4-12mo, yellow; nonCoV, dark blue) in HSPC and CD14^+^ monocytes. Genome track was generated using SparK (*110*) with aggregated pseudobulk ATAC-seq data. **(E)** Correlation of fold changes of gene expression and difference of DORC accessibility between late convalescent severe and healthy cohorts in HSPC and CD14^+^ monocytes.

Given the short half-life of circulating monocytes (*29*) we looked for a continued and active influence that underlies these persisting monocyte phenotypes months after acute infection. The persistence of an active plasma response in the S:2-4mo cohort (fig. S1B-C) could contribute to the transcriptional changes observed in these participants’ monocytes. However, no active plasma response was detected in the M:2-4mo cohort. (fig. S1B-C) Therefore, we hypothesized that inflammatory signaling present early in the disease process and resulting epigenetic memory in HSPC is conveyed to monocyte progeny for the 2-4 month period post-acute COVID-19 (or ICU). To assess if HSPC exhibited disease-associated phenotypes (similar to those for CD14^+^ monocytes), we investigated the expression of SC2 and SC3 defining genes within HSPC (n=28,069). Indeed, we found that S:2-4mo HSPC had a matching differential expression of severe/SC2 genes from our monocyte analysis (Fig. 3D, left). *NFKB1* and *MAPKAPK2* are two examples of inflammatory monocyte genes that also have matched upregulation in HSPC, particularly in S:2-4mo (Fig 3E, left panel). These genes feature expression patterns in monocytes that project directly onto, and are generally restricted to, inflammatory SC2 (Fig. 3E left; fig. S6D).

Similarly, M:2-4mo HSPC have matching differential expression of mild/SC3 genes (Fig. 3D, right; fig. S6E). *MX1* and *PARP14*, both interferon-stimulated genes (ISG) and anti-viral factors, had elevated expression in both HSPC and monocytes from M:2-4mo participants and were expressed specifically by the IFN^high^ SC3 monocytes as well as in nonCov (Fig. 3E, right; fig. S6E).

These data establish that COVID-19 durably reprograms HSPC transcriptomes, with discrete signatures for mild and severe disease, and with concordant transcriptional changes also detected in progeny monocytes. These changes include the activation of either anti-viral programs in mild COVID-19 or pro-inflammatory programs in severe disease, in both HSPC and monocytes. Notably, the activation of pro-inflammatory programs in HSPC was most prominent in severe COVID-19 (S:2-4mo) and was not observed to the same extent in non-COVID-19 ICU controls (nonCoV). Finally, these active months-long post-COVID-19 programs that drive discrete monocyte subsets generally resolve over time in both mild and severe diseases.

### Altered chromatin accessibility and durable epigenetic memory following COVID-19

To increase our power to detect epigenetic reprogramming of HSPC and progeny monocytes post-COVID-19, we analyzed additional samples in each cohort by bulk ATAC-seq on FACS sorted CD34^+^ HSPC (n=13) and CD14^+^ monocytes (n=27) (Fig. 1A, data S1-2). These additional data were integrated with the pseudobulk ATAC profiles from single-cell data for the corresponding cell types (n=32) for differential peak analysis among clinical groups that benefit from increased sample numbers (and complements DORC analysis). After peak calling and quality control, we identified differentially accessible regions (DAR) for both HSPC (n=263) and monocytes (n=9763) between disease cohorts (S:2-4mo, S:4-12mo, M:2-4mo, and nonCoV) and healthy controls as well as between S:2-4mo and nonCoV cohorts (data S1-2). Principal component analysis (PCA) of HSPC and monocyte ATAC-seq samples using DAR (Fig. 4A) highlighted consistent and significant epigenetic reprogramming associated with disease status in our cohort. PCA representation also confirmed that bulk and pseudobulk ATAC-seq samples were integrated well and clustered by cohort (Fig. 4A).

Disease-associated ATAC-seq peaks (9763 peaks in monocytes and 263 peaks in HPSCs) clustered into four major groups based on their accessibility profiles across all clinical cohorts (Fig. 4B-C). Overall, both in HSPC and monocytes, we observed that epigenetic changes are the most significant for the S:2-4mo cohort when compared to the healthy cohort, with mild COVID-19 (M:2-4mo) and nonCoV cohorts typically exhibiting similar epigenetic changes but with reduced magnitudes compared to severe COVID-19. Among the four HSPC peak clusters (Fig. 4B-C), we focused our attention on clusters where epigenetic activation is observed in both mild and severe COVID-19 participants (HSPC cluster 2 and HSPC cluster 4). HSPC cluster 2 contained peaks with COVID-19 specific accessibility (present in both early-convalescent mild *and* severe study participants) that persisted into late convalescence but was largely absent in the nonCoV cohort. These peaks were associated with genes that could influence immune cell phenotypes, including *IFI6*, *SOCS3*, *LDLR*, and *DOT1L*. Cluster 4 was composed of peaks that are more accessible in both S:2-4mo and nonCoV cohorts and were associated with pro-inflammatory molecules, including *IL8*, *S100A12*, *IL4R*, and *HLA-E* (Fig. 4C). While over time, many transcriptional changes resolved, notable epigenetic changes associated with disease persisted, suggesting that the epigenetic reprogramming at some loci in HPSC is longer lasting than the transcriptional signatures of the disease (Fig 4B-C).

Similarly, we detected four major clusters among disease-associated monocyte peaks (Fig. 4B-C). CD14^+^ monocyte Cluster 2 was observed in participants with severe disease (nonCOV and S:2-4mo) and featured an epigenetic activation signature, including pro-inflammatory molecules, dual-specificity phosphatase (DUSP) family phosphatases, several S100A factors, *CASP9*, *IRF1/8*, *IL8*, and cytokine receptors *IL4R*, and *IL6R* (associated with activated macrophage phenotypes) (*58*). Interestingly, this pro-inflammatory epigenetic signature was generally absent in mild COVID-19 and was resolved in late convalescence in participants with severe disease (S:4-12mo cohort). In contrast, another epigenetic activation signature in CD14^+^ monocytes was increased in both S:2-4mo and nonCoV cohorts and *persisted* into late convalescence (S:4-12mo) (Cluster 4) (Fig. 4B). This cluster included chemokines and chemokine receptors, inflammatory *S100A* factors, *IL1R1*, *TLR4*, *CASP1/8*, as well as *GATA4/5/6*, *JAK1/2*, antiinflammatory *IL10*, and several *KLF* transcription factors. Given the resolution of an active inflammatory response at the protein level in plasma in late convalescence (fig. S1B-C) and the short life-span of circulating monocytes (*29*), these persistent epigenetic changes could derive from HSPC intrinsic memory, durable changes in the HSPC environment (niche), or undetected changes in circulating factors. Examples of differentially accessible regions reveal distinct classes of epigenetic changes that occur post-COVID-19, including (i) transient peaks associated with early convalescence that return to baseline in late convalescence (e.g., *LDLR*, *TNFAIP3*, *IL17RA*, *IL1R1*); persistent peaks that remain altered in late convalescence (e.g., *IL8*, *IL10*, *OASL*); repressed peaks (often persistent; e.g., *DUSP1*, *IL1R1*); and other peaks with shared characteristics across both HSPC and monocytes such as the positively regulated *TNFRSF13B* (TACI) peak and the repressed *RUNX1* peak (Fig. 4D). Importantly, a number of interferon response genes, including *IFI6*, *IFITM3*, and *OASL*, were epigenetically active in late convalescence (S:4-12mo), yet corresponding gene expression changes were not detected (Fig. 3D, fig. S6E) suggestive of a poised epigenetic interferon program associated with severe COVID-19 (Fig. 4; see Fig. 5, below).

**Fig. 5.**
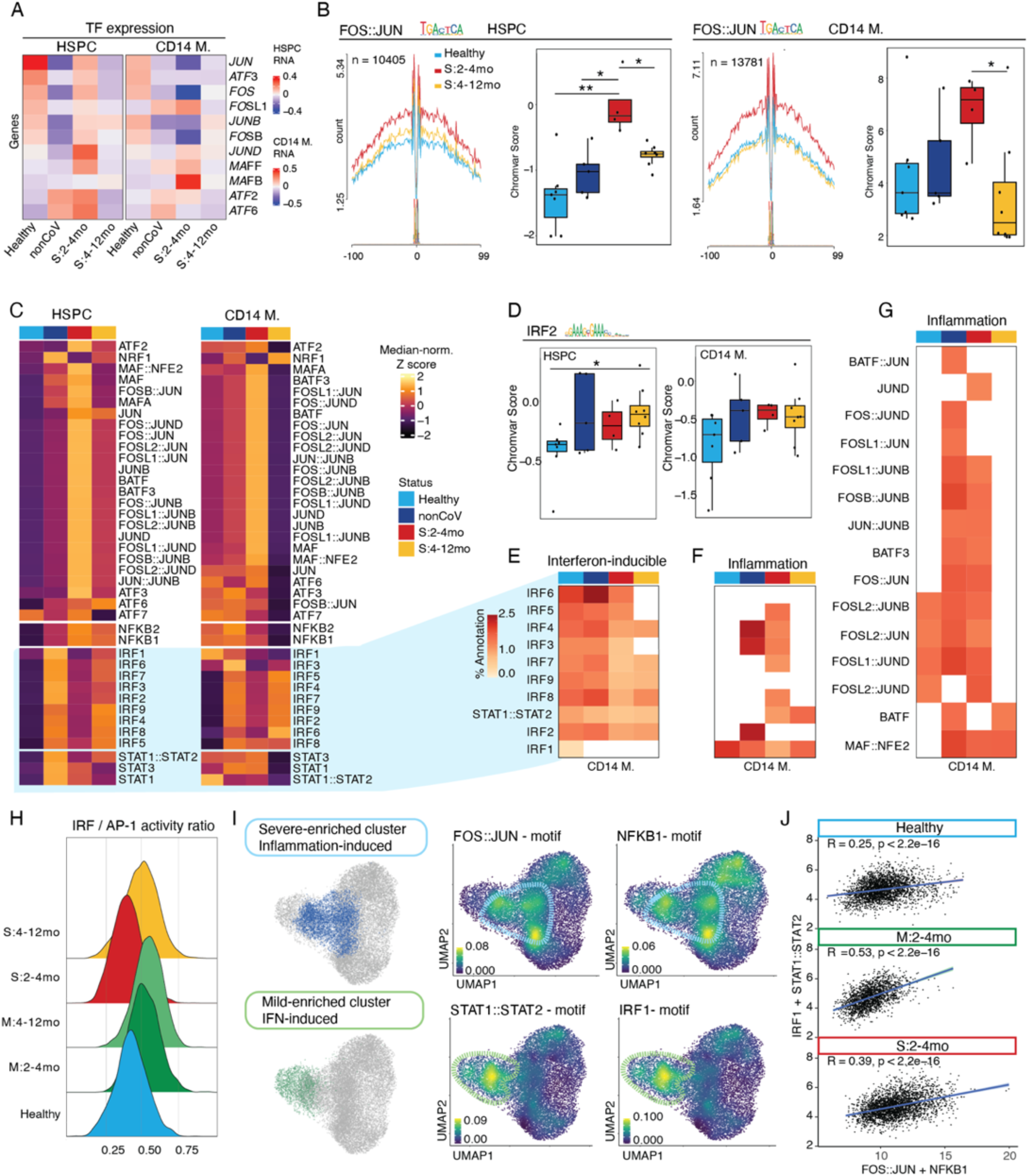
Post-COVID-19 memory in HSPC and monocytes features negative feedback of AP-1 and persistence of an IRF/STAT anti-viral program. **(A)** Heatmap for gene expression of selected AP-1 family genes in HSPC and CD14^+^ monocytes. **(B)** Transcription factor JUN::FOS footprints (HINT (*109*)) and boxplot for TF motif-associated chromatin accessibility (chromVAR (*112*)) activity in HSPC and CD 14^+^ monocytes across cohorts. **(C)** Heatmap presenting the Z-score-normalized median chromVAR (*112*) score for TFs in HSPC and CD14^+^ monocytes in each clinical cohort. **(D)** Boxplots for mean chromVAR score of IRF1 in HSPC and CD14^+^ monocytes for individual samples grouped by cohorts (t-test, * p < 0.05, Healthy as a reference). **(E-G)** Heatmap of gene module enrichment across cohorts for AP-1 and IRF TF families. Red shades indicate percent enrichment of the module. White shading indicates no statistical significance in the enrichment. **(H)** Histograms showing the distribution of aggregated AP1 and IRF TF families within each cell across cohorts. **(I)** CD14^+^ monocyte UMAP plots highlighting IFN-induced clusters (bottom) and inflammation-induced clusters (top). Enrichment of cluster-defining TF chromatin binding (chromVAR score) projected onto the CD14+ monocyte UMAP (right): FOS::JUN and NFKB1 motif enrichment for inflammation-induced cluster; STAT1::STAT2 and IRF1 for IFN-induced cluster. **(J)** Scatter plots showing the ratio of aggregate chromVAR scores for combined NFKB1+ FOS::JUN (x-axis), and combined IRF1+STAT1::STAT2 (y-axis).

To study the relationship between epigenetic and transcriptional changes in disease, we plotted COVID-19 associated gene expression changes versus changes in chromatin accessibility at cis-regulatory elements annotated to those genes based on our DORC analysis (Fig. 2F, fig. S5). This analysis highlighted a general correlation between chromatin accessibility and gene expression, with both measurements increasing or decreasing together for most genes. However, a subset of genes was characterized by altered chromatin accessibility in the absence of gene expression changes representing putatively poised or repressed genes (fig. S7A-B). Among these, *SOCS1* (suppressor of cytokine signaling 1), a negative feedback regulator of JAK1/2 and TLR signaling, was poised (3~5-fold increased DORC score in the absence of differential gene expression) in both HSPC and monocytes in late convalescence (S:4-12mo) (fig. S7A, right). These putatively poised genes identified from DORC analysis together with extensive differential chromatin accessibility in post-COVID-19 cohorts (i.e., monocyte Cluster 4, Fig. 4B-C) highlights that COVID-19 may epigenetically tune future inflammatory responses in addition to driving chromatin changes linked to active transcription programs.

We next focused on genes that were positively regulated (increased gene expression and DORC activity) in association with recovery from severe COVID-19 either transiently (i.e., only observed in S:2-4mo, Fig. 4E, left) or persistently (i.e., observed in S:4-12mo, Fig. 4E, right). Several genes in this positively regulated category relate to immune cell migration and activation, including *CD97*, *CCR7*, *S100A12*, *CCL4*, *CCL5*, and *IL7R*. One gene with persistent epigenetic and transcriptional changes in both HSPC and monocytes in late convalescence (S:4-12mo) is *ADGRE5* (*CD97*) (Fig. 4E, right two), an adhesion GPCR factor that guides myeloid cell migration and activity in inflamed tissues, is critical for protection from pneumococcal pneumonia, and required for optimal neutrophil recruitment and production of TNFa and IL-1β in infected lungs (*59*). Further, *CD97* expression on macrophages directs infiltration and activity to the synovial tissue in rheumatoid arthritis (RA) (*60*) and drives pathogenesis in mouse models of RA (*61*, *62*). A major ligand for *CD97* is *CD55*, which protects from complement-mediated damage, of interest given extensive and months-long complement activity in COVID-19 (fig. S1B). *CD97/CD55* interactions are elevated in multiple sclerosis and suggested to contribute to the inflammatory process (*63*). We note that the complement activated regulator of cell cycle (RGCC) had elevated DORC activity and RNA in monocytes of early convalescent study participants (S:2-4mo). Thus, complement-driven induction of *RGCC* may contribute to proliferative and altered hematopoiesis phenotypes in early convalescence (*64*).

Durable changes post-COVID-19 could also augment the ability of cells to recruit other cells to sites of inflammation via the chemokines *CCL4* and *CCL5*. DORC activity and RNA were elevated for *CCL4* (in both HSPC and monocytes) and *CCL5* (in monocytes) in late convalescence (S:4-12mo) (Fig. 4B-C). The inflammatory, antimicrobial *S100A12* gene had increased DORC activity and RNA expression in monocytes both in early and late convalescence (S:2-4mo and S:4-12mo). Further, in late convalescent monocytes (S:4-12mo), several genes characteristic of a persisting inflammatory monocyte program had both increased DORC activity and gene expression, including *CCR7*, *CCL4*, *CCL5*, *IL2RB*, and *IL7R. IL7R* and *IL2RB* were also epigenetically and transcriptionally activated in HSPC in late convalescence (S:4-12mo), suggestive of progenitor-progeny inheritance (Fig. 4E). *IL7R* is induced upon activation of CD14^+^ monocytes, regulates lung and tissue macrophage differentiation, and is a general feature of acute severe COVID-19 (*65*–*67*). Further, monocyte expression of *IL7R* is strongly influenced by an autoimmune *IL7R* polymorphism (*68*), indicating myeloid-specific epigenetic regulation of the locus. Thus, progenitor and monocyte upregulation of these inflammatory and migratory behavior promoting factors post-COVID-19 are suggestive of important and distinct post-infection phenotypes and warrant further study.

We then focused on genes that were repressed both in terms of gene expression and chromatin activity. Downregulated genes in late convalescence (S:4-12mo) featured critical negative feedback regulators of inflammation, *NFKBIA*, and *DUSP1* (*69*, *70*). In late convalescence, both in HSPC and monocytes, *NFKBIA* was among the most repressed genes (RNA and DORC). While repressed in late convalescence, inflammation-induced genes *NFKBIA* and negative regulator of NFκB, TNFAIP3 (A20) (*69*, *70*), were induced in early convalescence, likely by ongoing inflammatory signals (Fig. 4E). Highlighting the importance of NFKBIA-mediated negative feedback, a previous study using a model of the autoimmune disease Sjogren’s syndrome (induced by introduction of a human mutation of the NFκB motif within the *NFKBIA* gene promoter) established that reduced transcription of *NFKBIA* in response to NFκB pathway activation is causal in chronic inflammation (*71*). Repression (reduced RNA and DORC activity) of *DUSP1* (*MKP1*) is notable given its central role as a JNK/p38 phosphatase also induced by inflammation and critical for negative feedback of the inflammatory response, especially of TNFa (Fig. 4E) (*72*, *73*). Thus, two potent negative regulators of inflammation are durably downregulated post-severe-COVID-19 in both HSPC and their progeny monocytes, suggesting a potential mechanism for defective control of inflammation.

Among the most significant and consistent changes from the combined DORC/RNA analysis was the epigenetic and transcriptional repression of AP-1 transcription factor (TF) family members post-COVID-19. Among these, *JUN* was uniformly repressed, most prominently in HSPC, and *JUNB, FOSB*, and *FOSL2* were repressed in monocytes (Fig. 4E). AP-1 is critical in the regulation of the transcription response to MAPK activity, downstream of diverse cytokine receptors, TLR, and activating signals. Thus, with AP-1 TF activity a hallmark of inflammatory signaling and programming, repression of AP-1 factors themselves post-COVID-19 could represent a potent negative feedback program of inflammation.

### AP-1 and IRF activity define post-COVID-19 memory in HSPC and monocytes

Negative regulation of AP-1 TF expression and activity was recently described to characterize a program of “innate refractoriness,” or dampened inflammatory gene transcription, in response to influenza vaccination (*73*). Remarkably, the epigenetic features associated with innate refractoriness persisted for up to 6 months. It was speculated that HSPC propagated the program to short-lived progeny monocytes. Our observations of negative epigenetic and transcriptional regulation of AP-1 family members *JUN*, *JUNB*, *FOSB*, and *FOSL2* post-COVID-19 in both HSPC and monocytes (Fig. 4E) reveal the capacity for establishment and persistence of an AP-1 negative feedback program in HSPC following COVID-19 disease. Further, the epigenetic memory in HSPC that we describe (Fig. 4) provides an example of a program passed from progenitors to progeny monocytes that could explain persistent phenotypes beyond the lifespan of circulating monocytes following COVID-19 (as shown in this study) and following influenza vaccination (*44*).

We profiled the expression of several AP-1 factors across clinical cohorts and observed that there was decreased expression in both HSPC and monocytes following COVID-19 (Fig. 5A, fig. S8), including downregulation of *JUN* and *FOS* in monocytes. An exception was a switch in AP-1 family member expression in HSPC in early convalescence (S:2-4mo) (decreased *JUN*, *ATF3*, but increased *JUND*, *ATF2*, *ATF6*), which might result from ongoing signaling events driven by inflammation and associated emergency hematopoiesis. However, by late convalescence (S:4-12mo), the expression of several AP-1 family members was diminished both in HSPC and monocytes (Fig. 5A, and fig. S8). Next, we asked if this reduced AP-1 expression post-COVID-19 translated into a significant decrease in their binding to chromatin. First, we identified footprints— TF chromatin binding events inferred by TF-mediated steric occlusion of chromatin accessibility-defining Tn5 enzyme activity and protection of the DNA motif site—for different TFs using HINT (83) and compared the accessibility at these footprints across different clinical conditions. This analysis revealed higher activity of FOS:JUN TFs in the S:2-4mo cohort that is reduced over time (i.e., in late convalescent severe, S:4-12mo). Intriguingly, this increased accessibility was more prominent following COVID-19 and not observed to the same extent in nonCoV (Fig. 5B). Second, by identifying global chromatin accessibility at TF binding sites using ChromVAR (84), we observed that chromatin accessibility around binding sites of AP-1 complex members— particularly TFs from the JUN and FOS families— are increased in early convalescence (S:2-4mo) followed by decreased activity in late convalescence (S:4-12) (Fig. 5C, Fig. S9A). Together, these data point to a transient increase in AP-1 binding activity in early convalescence following COVID-19, followed by a decrease in late convalescence (S:4-12mo), both in HSPC and monocytes. Ongoing inflammatory signaling early following severe disease (S:2-4mo) is likely responsible for transient increases in AP-1 activity, while negative regulation of AP-1 family gene expression likely underlies the reduced activity observed in late convalescence (S:4-12mo). We suggest that these features of AP-1 factor repression and reduced steady-state chromatin binding are analogous to a state of innate immune refractoriness upon influenza vaccination (*44*). Beyond characterization of this state in post-COVID-19 monocytes, we also show that HSPC share this signature following COVID-19 and are the likely origin of its persistence in progeny monocytes.

We initially focused on AP-1 chromatin binding and its significance due to our identification of the prominent repression of AP-1 TFs. We next sought to identify, through footprint and motif accessibility analyses, other TFs with important roles in epigenetic programs following COVID-19. We surveyed TF activity associated with other inflammatory and anti-viral programs, including additional AP-1 members, interferon response factors (IRFs), STAT family, and NFkB members, that include critical mediators of signaling from the cytokine, TLR, and interferon receptors (IFNR). In contrast to AP-1 activity, chromatin accessibility at IRF motifs not only increased in early convalescence (S:2-4mo) but persisted into late convalescence (S:4-12mo) in both HSPC and monocytes. An exception to this pattern is decreased IRF1 activity in both early and late convalescence (S:2-4mo, S:4-12mo) (Fig. 5C-D).

To link these alterations in TF activity to meaningful gene targets, we associated TF footprints using HINT (83) to proximal genes and performed gene module enrichment analysis (Methods). As expected, we found that IRF target genes in both HSPC and monocytes were enriched in interferon-inducible genes across all clinical cohorts (Fig. 5E). Surprisingly, we also observed enrichment of inflammatory genes at sites of IRF activity, selectively in early post-COVID and post-ICU cohorts (S:2-3mo, nonCoV), suggesting that both COVID-19 and critical illness, in general, might be altering IRF binding activity and target genes (Fig. 5F). In addition, AP-1 family members displayed matching enrichment for inflammatory gene sets in the same cohorts (S:2-4mo and nonCoV) (Fig. 5G). Thus, a baseline IRF program targeting interferon-regulated genes appears to be redirected to inflammatory genes by inflammation-associated AP-1 activity. Consistent with a resolution of active inflammation in late convalescence (S:4-12mo) and AP-1 negative feedback (Fig. 5A-C), we observed a decrease in AP-1 chromatin binding activity and loss of inflammatory gene targeting by both AP-1 and IRF family members (Fig. 5F-G, yellow column), while ISG targeting by IRF factors was maintained (Fig. 5E).

To further explore the activity of AP-1 and IRF transcription factors in disease-associated monocyte subsets, we visualized the distribution of average AP-1 and IRF family TF activity in individual cells across the clinical cohorts. Consistent with the still-active inflammatory milieu and prevalent inflammatory monocyte phenotypes early following severe COVID-19 (S:2-4mo) (Fig. 3, fig. S1B-C), monocytes from this cohort featured higher distributions of AP-1 activity inferred from single cells (Fig. 5H, red distribution). AP-1 chromatin binding declined following mild disease (M:2-4mo and M:4-12mo, green distributions) and in late convalescence (S:4-12mo, yellow) (fig. S9B), fitting with post-infection (and post-vaccine (*44*)) AP-1 negative feedback. In contrast, IRF factor activity increased in mild early convalescence (M:2-4mo), consistent with prominent interferon (rather than inflammatory) program (fig. S9B). IRF activity also increased modestly in early and late convalescence following severe disease (S:2-4mo and S:4-12mo), suggesting that interferon signaling is driving IRF chromatin binding but that a competing inflammatory program results in the predominance of an inflammatory expression signature rather than an interferon signature.

Thus, we suggest that the ratio of IRF/AP-1 chromatin binding (Fig. 5H) is critical in driving either persisting inflammatory (low ratio, e.g., S:2-4mo) or persisting anti-viral (high ratio, e.g., M:2-4mo) programs in monocytes following infection. Despite evidence of IRF factor chromatin binding, S:2-4mo cohort monocytes fail to maintain the months-long interferon signature program apparent in mild (M:2-4mo) cohort driven by high-level STAT1:STAT2 and IRF1 activity (Fig. 3, right; Fig. 5I). Instead, prominent AP-1 and NFkB chromatin binding activity appear to direct a dominant inflammatory program that distinguishes the S:2-4mo monocyte phenotype (Fig. 5I-J). Visualization of independent FOS/JUN+NFkB and IRF1+STAT1:STAT2 programs across individual cells further highlights the influence of the AP-1/NFkB program in directing transcriptional programs despite the chromatin binding activity of IRF and STAT factors (Fig. 5J). Intriguingly, in late convalescence (S:4-12mo), following resolution of the inflammatory monocyte program and negative feedback of AP-1, IRF factor activity persists (Fig. 5C, H; fig. S9B). Together, these data reveal distinct, disease severity-dependent TF programs in HSPC and monocytes, with negative regulation of AP-1 and persisting IRF activity as standout features of the epigenetic memory of COVID-19.

## Discussion

Innate immune memory in response to natural infection in humans has yet to be well characterized. Moreover, whether and how hematopoietic stem and progenitor cells are altered in humans following infection is poorly understood. This study explores if COVID-19 and its associated inflammatory responses result in innate immune memory and whether these phenotypes propagate through HSPC even after the resolution of acute infection.

Enrichment of circulating CD34^+^ HSPC among PBMC via PBMC-PIE (PBMC analysis with progenitor input enrichment), paired with single-cell combined ATAC/RNA analysis, enabled deep characterization of these rare circulating CD34^+^ HSPC cells in diverse clinical cohorts. We reveal that while rare (~0.05% of PBMC), these cells accurately capture the diversity of bone marrow HSPC subsets. Therefore, PBMC-PIE serves as a powerful tool to study hematopoiesis, epigenetic programming of HSPC, and the relationship between progenitor and progeny cell types in blood without directly accessing the bone marrow. We applied this approach to study the lasting effects of COVID-19 and critical illness on hematopoiesis and central trained immunity concepts (*20*), with a unique cohort including study participants (i) following mild COVID-19 in early (M:2-4mo) and late (M:4-12mo) convalescence; (ii) following severe COVID-19 in early (S:2-4mo) and late (S:4-12mo) convalescence; and (iii) recovering from non-COVID-19 related critical illness (nonCoV). With these participants, we assessed plasma factors, SARS-CoV-2 antibodies, PBMC immunophenotypes by flow, and epigenomic and transcriptional changes in circulating HSPC and immune cells. Notably, we show that severe disease induces long-lasting epigenetic alterations in HSPC and their short-lived progeny, circulating monocytes.

Transcription factor activity, chromatin accessibility, and gene expression programs linked phenotypes of post-infection HSPC and monocytes. This indicates that inflammatory signaling in progenitor cells establishes epigenetic memory that not only persists in self-renewing stem cells but is also conveyed through differentiation to influence progeny cell phenotypes. Our results establish a precedent for central trained immunity following viral infection and suggest that recent observations of persistent monocyte epigenetic memory following influenza vaccination (*44*) may also derive from HSPC alterations. Beyond this epigenetic innate immune memory, we demonstrate that hematopoiesis is altered following COVID-19, with increases in myeloid and neutrophil progenitor populations.

### Potential Mechanisms Contributing to Long-term Sequelae following COVID-19

Our study design and cohort size were suitable for the detection of post-COVID-19 programs that are commonly altered after COVID-19 and severe disease. Further studies that follow long-term outcomes could parse the molecular features that associate with the vast range of unexplained long-term sequelae after severe (and sometimes mild) illness, including post-acute sequelae of SARS-CoV-2 infection (PASC) (*12*, *13*, *74*) and post-ICU syndrome (PICS) (*75*).

Indeed, our findings suggest potential molecular mechanisms that may contribute to PASC and PICS, including (i) a months-long active plasma response including complement activity, acute phase proteins, and vascular risk factors, which, if deposited in tissues, could contribute to ongoing inflammation; (ii) persistently activated (*CCL4*, *CCL5*, *IL7R*) and tissue-migratory (*CCR7*) monocyte phenotypes, with underlying HSPC epigenetic programs, that may continue to fuel inflammation and fibrosis, notably in lung and upper respiratory mucosa; and (iii) altered hematopoiesis including increased myeloid and especially neutrophil progenitors whose progeny may contribute to ongoing inflammation. Based upon neutrophil stimulating CSF-3 (G-CSF) production by lung epithelium in response to SARS-CoV-2 infection (*76*) and persistent CSF-3R upregulation in post-COVID-19 HSPC (Fig. 1–2), we suggest that a feedforward loop of CSF-3/CSF-3R could contribute to neutrophil driven pathology in both acute and, due to the durability of this response, also in chronic COVID-19 disease.

### Innate Immune Memory Following Mild and Severe SARS-CoV-2 Infection

One standout finding of our study is the identification of months-long altered monocyte programs following severe, but also mild, SARS-CoV-2 infection, suggesting monocytes may also contribute to chronic inflammation, either in affected tissues, *via* migratory and chemoattractant programs, or systemically. These epigenetic programs underlie distinct CD14^+^ monocyte phenotypes with variable persistence depending on disease severity. Unlike mild post-COVID-19 interferon programming, the HSPC and monocyte programs following severe COVID-19 are complex, with individual cells bearing mixed inflammatory and interferon signatures and reduced expression of key negative feedback factors *DUSP1* and *NFKBIA*. We show that mild (non-hospitalized) COVID-19 can result in a months-long epigenetic and transcriptional program, characterized by prominent IRF transcription factor activity and interferon-stimulated gene (ISG) expression (e.g., *ISG15*, *MX1*, *MX2*, *IFI44L*, *IFI44*, *OAS3*, *AOAH*, and *PARP14*). In contrast, CD14^+^ monocytes of patients recovering from severe disease (2-4 months post-acute, following discharge from the ICU) feature epigenetic and transcriptional signatures of inflammation likely mediated by NFkB and AP-1 TFs. This active inflammatory CD14^+^ monocyte program resolves in late convalescence (4-12 months), though a distinct epigenetic monocyte phenotype persists, including increased chromatin accessibility at certain chemokines (e.g., *CCL2*, *CCL7*, *CCL24*), chemokine receptors (e.g., *CCR1*, *CCR3*, *CCRL2*) ISG (e.g., *IFI6*, *SOC3*, *OASL*), and inflammatory genes (e.g., *IL8*, caspases, S100A genes). This highlights the importance of further research to better understand functional changes in the post-COVID-19 immune system and the clinical implications of these prolonged epigenetic signatures of severe COVID-19 in HPSCs and their progeny.

Further, persisting alterations in HSPC and monocytes following mild and severe SARS-CoV-2 infection suggests the possibility of months-long alterations in innate immune status as a general feature of diverse infections. We propose that this dynamic aspect of blood development and innate immune memory could have major implications for vaccine responses and design, understanding post-infectious inflammatory disease, non-genetic variance in responses to infection, and the epidemiology of seasonal infections. Based on these results, it is intriguing to speculate that acute viral infections may induce months-long anti-viral resilience programs similar to what we describe following mild SARS-CoV-2 infection. For example, as a corollary, the aberrantly low frequency of non-SARS-CoV-2 respiratory infections in the winter of 2020-2021 may have led to subsequent increased susceptibility to pathogenic viral infections and the unusual epidemic of respiratory syncytial virus and rhinovirus in the summer of 2020 (*77*).

### AP-1 and IRF Transcription Factor Programs Underlie Post-COVID-19 Phenotypes

How the transcriptional and epigenetic changes we observed in HSPC and monocytes following COVID-19 might alter cell differentiation, function, and response to stimulation is a critical open question. The molecular phenotypes of mild post-COVID-19 monocytes were very similar to those described in monocytes following adjuvanted influenza vaccine (H5N1+AS03), which provided a degree of heterologous anti-viral protection (Zika and Dengue) (*44*). Shared features of post-influenza vaccine and post-COVID-19 monocytes include a reduced AP-1 signature and increased IRF activity (which, in the case of COVID-19, persisted in both HSPC and monocytes). Notably, for up to one year into late convalescence following severe COVID-19, AP-1 activity returns to, or below, baseline, as a result of negative feedback regulation of AP-1 family members, while IRF factor activity remains elevated. Our single-cell analyses also reveal that both reduced AP-1 and increased IRF programs can co-exist within the same cells (Fig. 5J), raising the possibility that the persistent IRF activity following severe disease may represent a primed rather than active anti-viral program. Indeed, IRF factors interact with BAF complex (SWI/SNF) chromatin remodelers to maintain open or poised chromatin states and to drive active transcription (*78*). IRF1 activity, which drives active inflammation-responsive interferon-stimulated gene transcription, was reduced post-COVID-19, but several other IRF factors were increased, including IRF2 and IRF3, which have been shown to interact with the BAF complex to retain ISG in a poised state (*79*). We suggest that persisting IRF chromatin binding activity post-COVID-19 could result in increased poising and responsiveness of IRF target genes, in part through the maintenance of accessibility via IRF-BAF complex interactions.

While our study focuses on blood cells, it is important to point out that diverse other cell types have been demonstrated to harbor epigenetic memory (*21*, *80*, *81*). Particularly when they reside in affected tissues, these cells may change in their frequencies, differentiation programs, and phenotypes, and also retain epigenetic memory of anti-viral inflammation with important and enduring influence on tissue defense or sequelae (*80*, *82*).

Here, we present evidence of central trained immunity, in the form of epigenetic reprogramming in HSPC, in humans following viral infection and severe illness. Importantly, enrichment of rare circulating progenitor cells using PBMC-PIE was a critical advance enabling evaluation of hematopoietic stem and progenitor cells together with their progeny immune cells from peripheral blood samples. Extending this approach to diverse tissues (particularly those with resident stem and progenitor cells, e.g., intestinal epithelium) and disorders (hematologic disease, malignancy, inflammation, and infection) can unveil epigenetic and progenitor-based mechanisms of pathogenesis and inform therapeutic strategies and targets.

## Acknowledgments

We thank all the volunteers who donated their blood samples for this study. We thank the following members of the Josefowicz lab for critical feedback and helpful discussions: D. Ahimovic and C. Jiang. We thank B. Sleckman, J. Tyler, A. Rudensky, J. Sun, L. Ivashkiv, T. Holh, and R. Schwartz for helpful discussions. We thank Jane Cha and JAX creative department for help with figures and Stephen Sampson for help with editing.

## Funding

COVID-19 – Weill Cornell Medicine WCG-COVID-19 (WCM, SZJ)

NIH-R01AI148416-01 (WCM, SZJ)

NIH-U19AI144301-01 (WCM, SZJ)

NIH-R01AI148416-01S1 (WCM, SZJ)

NIH-RM1GM139738-01 (WCM, SZJ)

Asan Foundation, ROK (JGC)

NIH-NCI R25 CA233208 (MSKCC Computational Biomedicine Summer Program Training Grant, LP)

## Author contributions

Conceptualization: SZJ, DU, JGC, AR, RN

Resources: ZZ, AC, LL, DU, RK, SZJ

Methodology: SZJ, JGC, AR, DU

Investigation: JGC, AR, SS, DN, SM, BF, OK, AT, LP, MJB, JB, HS, DU, SZJ

Visualization: JGC, SS, AR, MJB, DN, LP, OK

Funding acquisition: SZJ

Project administration: SZJ

Supervision: SZJ, DU

Writing – original draft: SZJ, DU, JGC, AR, SS

Writing – review & editing: All authors

## Competing interests

J.D.B. holds patents related to ATAC-seq and scATAC-seq and serves on the Scientific Advisory Board of CAMP4 Therapeutics, seqWell, and CelSee.

## Data and materials availability

All raw and processed data are available on GEO database under accession number GSE190004. All other data used for analysis in this research are present in the manuscript or the Supplementary Materials. The code used for the downstream analysis is available upon request to correspondence authors and will be made anonymously available to reviewers upon submission.

## Supplementary Materials

### Summary

Materials and Methods

Supplementary Text

Figs. S1 to S9

Table. S1

Data S1 to S3

References and note (81–117)

## References

1. Y. Xiong, Y. Liu, L. Cao, D. Wang, M. Guo, A. Jiang, D. Guo, W. Hu, J. Yang, Z. Tang, H. Wu, Y. Lin, M. Zhang, Q. Zhang, M. Shi, Y. Liu, Y. Zhou, K. Lan, Y. Chen, Transcriptomic characteristics of bronchoalveolar lavage fluid and peripheral blood mononuclear cells in COVID-19 patients. Emerg. Microbes Infect. 9, 761–770 (2020).

2. D. Yang, H. Chu, Y. Hou, Y. Chai, H. Shuai, A. C.-Y. Lee, X. Zhang, Y. Wang, B. Hu, X. Huang, T. T.-T. Yuen, J.-P. Cai, J. Zhou, S. Yuan, A. J. Zhang, J. F.-W. Chan, K.-Y. Yuen, Attenuated Interferon and Proinflammatory Response in SARS-CoV-2-Infected Human Dendritic Cells Is Associated With Viral Antagonism of STAT1 Phosphorylation. J. Infect. Dis. 222, 734–745 (2020).

3. J. Wang, M. Jiang, X. Chen, L. J. Montaner, Cytokine storm and leukocyte changes in mild versus severe SARS-CoV-2 infection: Review of 3939 COVID-19 patients in China and emerging pathogenesis and therapy concepts. J. Leukoc. Biol. 108, 17–41 (2020).

4. C. Lucas, P. Wong, J. Klein, T. B. R. Castro, J. Silva, M. Sundaram, M. K. Ellingson, T. Mao, J. E. Oh, B. Israelow, T. Takahashi, M. Tokuyama, P. Lu, A. Venkataraman, A. Park, S. Mohanty, H. Wang, A. L. Wyllie, C. B. F. Vogels, R. Earnest, A. Iwasaki, Longitudinal analyses reveal immunological misfiring in severe COVID-19. Nature. 584, 463–469 (2020).

5. Q. Zhang, P. Bastard, Z. Liu, J. Le Pen, M. Moncada-Velez, J. Chen, M. Ogishi, I. K. D. Sabli, S. Hodeib, C. Korol, J. Rosain, K. Bilguvar, J. Ye, A. Bolze, B. Bigio, R. Yang, A. A. Arias, Q. Zhou, Y. Zhang, F. Onodi, J.-L. Casanova, Inborn errors of type I IFN immunity in patients with life-threatening COVID-19. Science. 370 (2020), doi:10.1126/science.abd4570.

6. R. Zhou, K. K.-W. To, Y.-C. Wong, L. Liu, B. Zhou, X. Li, H. Huang, Y. Mo, T.-Y. Luk, T. T.-K. Lau, P. Yeung, W.-M. Chan, A. K.-L. Wu, K.-C. Lung, O. T.-Y. Tsang, W.-S. Leung, I. F.-N. Hung, K.-Y. Yuen, Z. Chen, Acute SARS-CoV-2 Infection Impairs Dendritic Cell and T Cell Responses. Immunity. 53, 864–877.e5 (2020).

7. P. S. Arunachalam, F. Wimmers, C. K. P. Mok, R. A. P. M. Perera, M. Scott, T. Hagan, N. Sigal, Y. Feng, L. Bristow, O. Tak-Yin Tsang, D. Wagh, J. Coller, K. L. Pellegrini, D. Kazmin, G. Alaaeddine, W. S. Leung, J. M. C. Chan, T. S. H. Chik, C. Y. C. Choi, C. Huerta, B. Pulendran, Systems biological assessment of immunity to mild versus severe COVID-19 infection in humans. Science. 369, 1210–1220 (2020).

8. B. Israelow, E. Song, T. Mao, P. Lu, A. Meir, F. Liu, M. M. Alfajaro, J. Wei, H. Dong, R. J. Homer, A. Ring, C. B. Wilen, A. Iwasaki, Mouse model of SARS-CoV-2 reveals inflammatory role of type I interferon signaling. J. Exp. Med. 217 (2020), doi:10.1084/jem.20201241.

9. J. Hadjadj, N. Yatim, L. Barnabei, A. Corneau, J. Boussier, N. Smith, H. Péré, B. Charbit, V. Bondet, C. Chenevier-Gobeaux, P. Breillat, N. Carlier, R. Gauzit, C. Morbieu, F. Pène, N. Marin, N. Roche, T.-A. Szwebel, S. H. Merkling, J.-M. Treluyer, B. Terrier, Impaired type I interferon activity and inflammatory responses in severe COVID-19 patients. Science. 369, 718–724 (2020).

10. D. Acharya, G. Liu, M. U. Gack, Dysregulation of type I interferon responses in COVID-19. Nat. Rev. Immunol. 20, 397–398 (2020).

11. NIH launches new initiative to study “Long COVID” | National Institutes of Health (NIH), (available at https://www.nih.gov/about-nih/who-we-are/nih-director/statements/nih-launches-new-initiative-study-long-covid).

12. R. K. Ramakrishnan, T. Kashour, Q. Hamid, R. Halwani, I. M. Tleyjeh, Unraveling the Mystery Surrounding Post-Acute Sequelae of COVID-19. Front. Immunol. 12, 686029 (2021).

13. C. Del Rio, L. F. Collins, P. Malani, Long-term Health Consequences of COVID-19. JAMA. 324, 1723–1724 (2020).

14. M. J. Carter, M. Fish, A. Jennings, K. J. Doores, P. Wellman, J. Seow, S. Acors, C. Graham, E. Timms, J. Kenny, S. Neil, M. H. Malim, S. M. Tibby, M. Shankar-Hari, Peripheral immunophenotypes in children with multisystem inflammatory syndrome associated with SARS-CoV-2 infection. Nat. Med. 26, 1701–1707 (2020).

15. C. R. Consiglio, N. Cotugno, F. Sardh, C. Pou, D. Amodio, L. Rodriguez, Z. Tan, S. Zicari, A. Ruggiero, G. R. Pascucci, V. Santilli, T. Campbell, Y. Bryceson, D. Eriksson, J. Wang, A. Marchesi, T. Lakshmikanth, A. Campana, A. Villani, P. Rossi, P. Brodin, The Immunology of Multisystem Inflammatory Syndrome in Children with COVID-19. Cell. 183, 968–981.e7 (2020).

16. C. N. Gruber, R. S. Patel, R. Trachtman, L. Lepow, F. Amanat, F. Krammer, K. M. Wilson, K. Onel, D. Geanon, K. Tuballes, M. Patel, K. Mouskas, T. O’Donnell, E. Merritt, N. W. Simons, V. Barcessat, D. M. Del Valle, S. Udondem, G. Kang, S. Gangadharan, D. Bogunovic, Mapping Systemic Inflammation and Antibody Responses in Multisystem Inflammatory Syndrome in Children (MIS-C). Cell. 183, 982–995.e14 (2020).

17. M. W. Tenforde, S. B. Morris, Multisystem inflammatory syndrome in adults: coming into focus. Chest. 159, 471–472 (2021).

18. S. B. Morris, N. G. Schwartz, P. Patel, L. Abbo, L. Beauchamps, S. Balan, E. H. Lee, R. Paneth-Pollak, A. Geevarughese, M. K. Lash, M. S. Dorsinville, V. Ballen, D. P. Eiras, C. Newton-Cheh, E. Smith, S. Robinson, P. Stogsdill, S. Lim, S. E. Fox, G. Richardson, S. Godfred-Cato, Case Series of Multisystem Inflammatory Syndrome in Adults Associated with SARS-CoV-2 Infection - United Kingdom and United States, March-August 2020. MMWR Morb Mortal Wkly Rep. 69, 1450–1456 (2020).

19. M. G. Netea, J. Domínguez-Andrés, L. B. Barreiro, T. Chavakis, M. Divangahi, E. Fuchs, L. A. B. Joosten, J. W. M. van der Meer, M. M. Mhlanga, W. J. M. Mulder, N. P. Riksen, A. Schlitzer, J. L. Schultze, C. Stabell Benn, J. C. Sun, R. J. Xavier, E. Latz, Defining trained immunity and its role in health and disease. Nat. Rev. Immunol. 20, 375–388 (2020).

20. S. Bekkering, J. Domínguez-Andrés, L. A. B. Joosten, N. P. Riksen, M. G. Netea, Trained immunity: reprogramming innate immunity in health and disease. Annu. Rev. Immunol. 39, 667–693 (2021).

21. S. Naik, S. B. Larsen, N. C. Gomez, K. Alaverdyan, A. Sendoel, S. Yuan, L. Polak, A. Kulukian, S. Chai, E. Fuchs, Inflammatory memory sensitizes skin epithelial stem cells to tissue damage. Nature. 550, 475–480 (2017).

22. I. Mitroulis, K. Ruppova, B. Wang, L.-S. Chen, M. Grzybek, T. Grinenko, A. Eugster, M. Troullinaki, A. Palladini, I. Kourtzelis, A. Chatzigeorgiou, A. Schlitzer, M. Beyer, L. A. B. Joosten, B. Isermann, M. Lesche, A. Petzold, K. Simons, I. Henry, A. Dahl, T. Chavakis, Modulation of myelopoiesis progenitors is an integral component of trained immunity. Cell. 172, 147–161.e12 (2018).

23. J. Kleinnijenhuis, J. Quintin, F. Preijers, L. A. B. Joosten, D. C. Ifrim, S. Saeed, C. Jacobs, J. van Loenhout, D. de Jong, H. G. Stunnenberg, R. J. Xavier, J. W. M. van der Meer, R. van Crevel, M. G. Netea, Bacille Calmette-Guerin induces NOD2-dependent nonspecific protection from reinfection via epigenetic reprogramming of monocytes. Proc Natl Acad Sci USA. 109, 17537–17542 (2012).

24. J. Quintin, S. Saeed, J. H. A. Martens, E. J. Giamarellos-Bourboulis, D. C. Ifrim, C. Logie, L. Jacobs, T. Jansen, B.-J. Kullberg, C. Wijmenga, L. A. B. Joosten, R. J. Xavier, J. W. M. van der Meer, H. G. Stunnenberg, M. G. Netea, Candida albicans infection affords protection against reinfection via functional reprogramming of monocytes. Cell Host Microbe. 12, 223–232 (2012).

25. J. Kleinnijenhuis, J. Quintin, F. Preijers, L. A. B. Joosten, C. Jacobs, R. J. Xavier, J. W. M. van der Meer, R. van Crevel, M. G. Netea, BCG-induced trained immunity in NK cells: Role for nonspecific protection to infection. Clin. Immunol. 155, 213–219 (2014).

26. E. Kaufmann, J. Sanz, J. L. Dunn, N. Khan, L. E. Mendonça, A. Pacis, F. Tzelepis, E. Pernet, A. Dumaine, J.-C. Grenier, F. Mailhot-Léonard, E. Ahmed, J. Belle, R. Besla, B. Mazer, I. L. King, A. Nijnik, C. S. Robbins, L. B. Barreiro, M. Divangahi, BCG Educates Hematopoietic Stem Cells to Generate Protective Innate Immunity against Tuberculosis. Cell. 172, 176–190.e19 (2018).

27. J. C. Dos Santos, A. M. Barroso de Figueiredo, M. V. Teodoro Silva, B. Cirovic, L. C. J. de Bree, M. S. M. A. Damen, S. J. C. F. M. Moorlag, R. S. Gomes, M. M. Helsen, M. Oosting, S. T. Keating, A. Schlitzer, M. G. Netea, F. Ribeiro-Dias, L. A. B. Joosten, β-Glucan-Induced Trained Immunity Protects against Leishmania braziliensis Infection: a Crucial Role for IL-32. Cell Rep. 28, 2659–2672.e6 (2019).

28. B. Cirovic, L. C. J. de Bree, L. Groh, B. A. Blok, J. Chan, W. J. F. M. van der Velden, M. E. J. Bremmers, R. van Crevel, K. Händler, S. Picelli, J. Schulte-Schrepping, K. Klee, M. Oosting, V. A. C. M. Koeken, J. van Ingen, Y. Li, C. S. Benn, J. L. Schultze, L. A. B. Joosten, N. Curtis, A. Schlitzer, BCG vaccination in humans elicits trained immunity via the hematopoietic progenitor compartment. Cell Host Microbe. 28, 322–334.e5 (2020).

29. A. A. Patel, Y. Zhang, J. N. Fullerton, L. Boelen, A. Rongvaux, A. A. Maini, V. Bigley, R. A. Flavell, D. W. Gilroy, B. Asquith, D. Macallan, S. Yona, The fate and lifespan of human monocyte subsets in steady state and systemic inflammation. J. Exp. Med. 214, 1913–1923 (2017).

30. A. Christ, P. Günther, M. A. R. Lauterbach, P. Duewell, D. Biswas, K. Pelka, C. J. Scholz, M. Oosting, K. Haendler, K. Baβler, K. Klee, J. Schulte-Schrepping, T. Ulas, S. J. C. F. M. Moorlag, V. Kumar, M. H. Park, L. A. B. Joosten, L. A. Groh, N. P. Riksen, T. Espevik, E. Latz, Western Diet Triggers NLRP3-Dependent Innate Immune Reprogramming. Cell. 172, 162–175.e14 (2018).

31. L. Kong, S. J. C. F. M. Moorlag, A. Lefkovith, B. Li, V. Matzaraki, L. van Emst, H. A. Kang, I. Latorre, M. Jaeger, L. A. B. Joosten, M. G. Netea, R. J. Xavier, Single-cell transcriptomic profiles reveal changes associated with BCG-induced trained immunity and protective effects in circulating monocytes. Cell Rep. 37, 110028 (2021).

32. K. J. Jensen, N. Larsen, S. Biering-Sørensen, A. Andersen, H. B. Eriksen, I. Monteiro, D. Hougaard, P. Aaby, M. G. Netea, K. L. Flanagan, C. S. Benn, Heterologous immunological effects of early BCG vaccination in low-birth-weight infants in Guinea-Bissau: a randomized-controlled trial. J. Infect. Dis. 211, 956–967 (2015).

33. M. L. T. Berendsen, C. B. Øland, P. Bles, A. K. G. Jensen, P.-E. Kofoed, H. Whittle, L. C. J. de Bree, M. G. Netea, C. Martins, C. S. Benn, P. Aaby, Maternal Priming: Bacillus Calmette-Guérin (BCG) Vaccine Scarring in Mothers Enhances the Survival of Their Child With a BCG Vaccine Scar. J. Pediatric Infect. Dis. Soc. 9, 166–172 (2020).

34. C. Näslund, Resultats des experiences de vaccination par le BCG poursuivies dans le Norrbotten (Suède)(Septembre 1927–Décembre 1931). Vaccination Preventative de Tuberculose, Rapports et Documents. Paris: Institut Pasteur (1932).

35. N. Zhu, D. Zhang, W. Wang, X. Li, B. Yang, J. Song, X. Zhao, B. Huang, W. Shi, R. Lu, P. Niu, F. Zhan, X. Ma, D. Wang, W. Xu, G. Wu, G. F. Gao, W. Tan, China Novel Coronavirus Investigating and Research Team, A Novel Coronavirus from Patients with Pneumonia in China, 2019. N. Engl. J. Med. 382, 727–733 (2020).

36. C. Huang, Y. Wang, X. Li, L. Ren, J. Zhao, Y. Hu, L. Zhang, G. Fan, J. Xu, X. Gu, Z. Cheng, T. Yu, J. Xia, Y. Wei, W. Wu, X. Xie, W. Yin, H. Li, M. Liu, Y. Xiao, B. Cao, Clinical features of patients infected with 2019 novel coronavirus in Wuhan, China. Lancet. 395, 497–506 (2020).

37. W.-J. Guan, Z.-Y. Ni, Y. Hu, W.-H. Liang, C.-Q. Ou, J.-X. He, L. Liu, H. Shan, C.-L. Lei, D. S. C. Hui, B. Du, L.-J. Li, G. Zeng, K.-Y. Yuen, R.-C. Chen, C.-L. Tang, T. Wang, P.-Y. Chen, J. Xiang, S.-Y. Li, China Medical Treatment Expert Group for Covid-19, Clinical characteristics of coronavirus disease 2019 in china. N. Engl. J. Med. 382, 1708–1720 (2020).

38. D. Wang, B. Hu, C. Hu, F. Zhu, X. Liu, J. Zhang, B. Wang, H. Xiang, Z. Cheng, Y. Xiong, Y. Zhao, Y. Li, X. Wang, Z. Peng, Clinical Characteristics of 138 Hospitalized Patients With 2019 Novel Coronavirus-Infected Pneumonia in Wuhan, China. JAMA. 323, 1061–1069 (2020).

39. N. Chen, M. Zhou, X. Dong, J. Qu, F. Gong, Y. Han, Y. Qiu, J. Wang, Y. Liu, Y. Wei, J. Xia, T. Yu, X. Zhang, L. Zhang, Epidemiological and clinical characteristics of 99 cases of 2019 novel coronavirus pneumonia in Wuhan, China: a descriptive study. Lancet. 395, 507–513 (2020).

40. J. C. Holter, S. E. Pischke, E. de Boer, A. Lind, S. Jenum, A. R. Holten, K. Tonby, A. Barratt-Due, M. Sokolova, C. Schjalm, V. Chaban, A. Kolderup, T. Tran, T. Tollefsrud Gjølberg, L. G. Skeie, L. Hesstvedt, V. Ormåsen, B. Fevang, C. Austad, K. E. Müller, T. E. Mollnes, Systemic complement activation is associated with respiratory failure in COVID-19 hospitalized patients. Proc Natl Acad Sci USA. 117, 25018–25025 (2020).

41. D. A. Berlin, R. M. Gulick, F. J. Martinez, Severe Covid-19. N. Engl. J. Med. 383, 2451–2460 (2020).

42. Y. Xie, B. Bowe, Z. Al-Aly, Burdens of post-acute sequelae of COVID-19 by severity of acute infection, demographics and health status. Nat. Commun. 12, 6571 (2021).

43. A. Jaffri, U. A. Jaffri, Post-Intensive care syndrome and COVID-19: crisis after a crisis? Heart Lung. 49, 883–884 (2020).

44. F. Wimmers, M. Donato, A. Kuo, T. Ashuach, S. Gupta, C. Li, M. Dvorak, M. H. Foecke, S. E. Chang, T. Hagan, S. E. De Jong, H. T. Maecker, R. van der Most, P. Cheung, M. Cortese, S. E. Bosinger, M. Davis, N. Rouphael, S. Subramaniam, N. Yosef, B. Pulendran, The single-cell epigenomic and transcriptional landscape of immunity to influenza vaccination. Cell. 184, 3915–3935.e21 (2021).

45. A. J. Wilk, A. Rustagi, N. Q. Zhao, J. Roque, G. J. Martínez-Colón, J. L. McKechnie, G. T. Ivison, T. Ranganath, R. Vergara, T. Hollis, L. J. Simpson, P. Grant, A. Subramanian, A. J. Rogers, C. A. Blish, A single-cell atlas of the peripheral immune response in patients with severe COVID-19. Nat. Med. 26, 1070–1076 (2020).

46. E. Stephenson, G. Reynolds, R. A. Botting, F. J. Calero-Nieto, M. D. Morgan, Z. K. Tuong, K. Bach, W. Sungnak, K. B. Worlock, M. Yoshida, N. Kumasaka, K. Kania, J. Engelbert, B. Olabi, J. S. Spegarova, N. K. Wilson, N. Mende, L. Jardine, L. C. S. Gardner, I. Goh, Cambridge Institute of Therapeutic Immunology and Infectious Disease-National Institute of Health Research (CITIID-NIHR) COVID-19 BioResource Collaboration, Single-cell multi-omics analysis of the immune response in COVID-19. Nat. Med. (2021), doi:10.1038/s41591-021-01329-2.

47. J. Schulte-Schrepping, N. Reusch, D. Paclik, K. Baβler, S. Schlickeiser, B. Zhang, B. Krämer, T. Krammer, S. Brumhard, L. Bonaguro, E. De Domenico, D. Wendisch, M. Grasshoff, T. S. Kapellos, M. Beckstette, T. Pecht, A. Saglam, O. Dietrich, H. E. Mei, A. R. Schulz, Deutsche COVID-19 OMICS Initiative (DeCOI), Severe COVID-19 Is Marked by a Dysregulated Myeloid Cell Compartment. Cell. 182, 1419–1440.e23 (2020).

48. J. D. Buenrostro, M. R. Corces, C. A. Lareau, B. Wu, A. N. Schep, M. J. Aryee, R. Majeti, H. Y. Chang, W. J. Greenleaf, Integrated Single-Cell Analysis Maps the Continuous Regulatory Landscape of Human Hematopoietic Differentiation. Cell. 173, 1535–1548.e16 (2018).

49. J. M. Granja, S. Klemm, L. M. McGinnis, A. S. Kathiria, A. Mezger, M. R. Corces, B. Parks, E. Gars, M. Liedtke, G. X. Y. Zheng, H. Y. Chang, R. Majeti, W. J. Greenleaf, Single-cell multiomic analysis identifies regulatory programs in mixed-phenotype acute leukemia. Nat. Biotechnol. 37, 1458–1465 (2019).

50. D. Pellin, M. Loperfido, C. Baricordi, S. L. Wolock, A. Montepeloso, O. K. Weinberg, A. Biffi, A. M. Klein, L. Biasco, A comprehensive single cell transcriptional landscape of human hematopoietic progenitors. Nat. Commun. 10, 2395 (2019).

51. H. M. Mehta, S. J. Corey, G-CSF, the guardian of granulopoiesis. Semin. Immunol., 101515 (2021).

52. L. Mellett, S. A. Khader, S100A8/A9 in COVID-19 pathogenesis: Impact on clinical outcomes. Cytokine Growth Factor Rev. (2021), doi:10.1016/j.cytogfr.2021.10.004.

53. S. Ma, B. Zhang, L. M. LaFave, A. S. Earl, Z. Chiang, Y. Hu, J. Ding, A. Brack, V. K. Kartha, T. Tay, T. Law, C. Lareau, Y.-C. Hsu, A. Regev, J. D. Buenrostro, Chromatin Potential Identified by Shared Single-Cell Profiling of RNA and Chromatin. Cell. 183, 1103–1116.e20 (2020).

54. B. Lehnertz, J. Chagraoui, T. MacRae, E. Tomellini, S. Corneau, N. Mayotte, I. Boivin, A. Durand, D. Gracias, G. Sauvageau, HLF expression defines the human hematopoietic stem cell state. Blood (2021), doi:10.1182/blood.2021010745.

55. C. Capron, Y. Lécluse, A. L. Kaushik, A. Foudi, C. Lacout, D. Sekkai, I. Godin, O. Albagli, I. Poullion, F. Svinartchouk, E. Schanze, W. Vainchenker, F. Sablitzky, A. Bennaceur-Griscelli, D. Duménil, The SCL relative LYL-1 is required for fetal and adult hematopoietic stem cell function and B-cell differentiation. Blood. 107, 4678–4686 (2006).

56. C. Yao, S. A. Bora, T. Parimon, T. Zaman, O. A. Friedman, J. A. Palatinus, N. S. Surapaneni, Y. P. Matusov, G. Cerro Chiang, A. G. Kassar, N. Patel, C. E. R. Green, A. W. Aziz, H. Suri, J. Suda, A. A. Lopez, G. A. Martins, B. R. Stripp, S. A. Gharib, H. S. Goodridge, P. Chen, Cell-Type-Specific Immune Dysregulation in Severely Ill COVID-19 Patients. Cell Rep. 34, 108590 (2021).

57. I. Zschiedrich, U. Hardeland, A. Krones-Herzig, M. Berriel Diaz, A. Vegiopoulos, J. Müggenburg, D. Sombroek, T. G. Hofmann, R. Zawatzky, X. Yu, N. Gretz, M. Christian, R. White, M. G. Parker, S. Herzig, Coactivator function of RIP140 for NFkappaB/RelA-dependent cytokine gene expression. Blood. 112, 264–276 (2008).

58. N. Lawlor, D. Nehar-Belaid, J. D. S. Grassmann, M. Stoeckius, P. Smibert, M. L. Stitzel, V. Pascual, J. Banchereau, A. Williams, D. Ucar, Single Cell Analysis of Blood Mononuclear Cells Stimulated Through Either LPS or Anti-CD3 and Anti-CD28. Front. Immunol. 12, 636720 (2021).

59. J. C. Leemans, A. A. te Velde, S. Florquin, R. J. Bennink, K. de Bruin, R. A. W. van Lier, T. van der Poll, J. Hamann, The epidermal growth factor-seven transmembrane (EGF-TM7) receptor CD97 is required for neutrophil migration and host defense. J. Immunol. 172, 1125–1131 (2004).

60. J. Hamann, J. O. Wishaupt, R. A. van Lier, T. J. Smeets, F. C. Breedveld, P. P. Tak, Expression of the activation antigen CD97 and its ligand CD55 in rheumatoid synovial tissue. Arthritis Rheum. 42, 650–658 (1999).

61. E. N. Kop, J. Adriaansen, T. J. M. Smeets, M. J. Vervoordeldonk, R. A. W. van Lier, J. Hamann, P. P. Tak, CD97 neutralisation increases resistance to collagen-induced arthritis in mice. Arthritis Res. Ther. 8, R155 (2006).

62. R. M. Hoek, D. de Launay, E. N. Kop, A. S. Yilmaz-Elis, F. Lin, K. A. Reedquist, J. S. Verbeek, M. E. Medof, P. P. Tak, J. Hamann, Deletion of either CD55 or CD97 ameliorates arthritis in mouse models. Arthritis Rheum. 62, 1036–1042 (2010).

63. L. Visser, A. F. de Vos, J. Hamann, M.-J. Melief, M. van Meurs, R. A. W. van Lier, J. D. Laman, R. Q. Hintzen, Expression of the EGF-TM7 receptor CD97 and its ligand CD55 (DAF) in multiple sclerosis. J. Neuroimmunol. 132, 156–163 (2002).

64. M. Fosbrink, F. Niculescu, H. Rus, The Role of C5b-9 Terminal Complement Complex in Activation of the Cell Cycle and Transcription. IR. 31, 37–46 (2005).

65. E. Evren, E. Ringqvist, K. P. Tripathi, N. Sleiers, I. C. Rives, A. Alisjahbana, Y. Gao, D. Sarhan, T. Halle, C. Sorini, R. Lepzien, N. Marquardt, J. Michaëlsson, A. Smed-Sörensen, J. Botling, M. C. I. Karlsson, E. J. Villablanca, T. Willinger, Distinct developmental pathways from blood monocytes generate human lung macrophage diversity. Immunity (2020), doi:10.1016/j.immuni.2020.12.003.

66. G. A. Leung, T. Cool, C. H. Valencia, A. Worthington, A. E. Beaudin, E. C. Forsberg, The lymphoid-associated interleukin 7 receptor (IL7R) regulates tissue-resident macrophage development. Development. 146 (2019), doi:10.1242/dev.176180.

67. B. Zhang, Y. Zhang, L. Xiong, Y. Li, Y. Zhang, J. Zhao, H. Jiang, C. Li, Y. Liu, X. Liu, H. Liu, Y.-F. Ping, Q. C. Zhang, Z. Zhang, X.-W. Bian, Y. Zhao, X. Hu, CD127 imprints functional heterogeneity to diversify monocyte responses in human inflammatory diseases. BioRxiv (2020), doi:10.1101/2020.11.10.376277.

68. H. Al-Mossawi, N. Yager, C. A. Taylor, E. Lau, S. Danielli, J. de Wit, J. Gilchrist, I. Nassiri, E. A. Mahe, W. Lee, L. Rizvi, S. Makino, J. Cheeseman, M. Neville, J. C. Knight, P. Bowness, B. P. Fairfax, Context-specific regulation of surface and soluble IL7R expression by an autoimmune risk allele. Nat. Commun. 10, 4575 (2019).

69. B. Gil-Araujo, M.-V. Toledo Lobo, M. Gutiérrez-Salmerón, J. Gutiérrez-Pitalúa, S. Ropero, J. C. Angulo, A. Chiloeches, M. Lasa, Dual specificity phosphatase 1 expression inversely correlates with NF-κB activity and expression in prostate cancer and promotes apoptosis through a p38 MAPK dependent mechanism. Mol. Oncol. 8, 27–38 (2014).

70. F. Arenzana-Seisdedos, P. Turpin, M. Rodriguez, D. Thomas, R. T. Hay, J. L. Virelizier, C. Dargemont, Nuclear localization of I kappa B alpha promotes active transport of NF-kappa B from the nucleus to the cytoplasm. J. Cell Sci. 110(Pt 3), 369–378 (1997).

71. J. T. Cooper, D. M. Stroka, C. Brostjan, A. Palmetshofer, F. H. Bach, C. Ferran, A20 blocks endothelial cell activation through a NF-kappaB-dependent mechanism. J. Biol. Chem. 271, 18068–18073 (1996).

72. B. Peng, J. Ling, A. J. Lee, Z. Wang, Z. Chang, W. Jin, Y. Kang, R. Zhang, D. Shim, H. Wang, J. B. Fleming, H. Zheng, S.-C. Sun, P. J. Chiao, Defective feedback regulation of NF-kappaB underlies Sjogren’s syndrome in mice with mutated kappaB enhancers of the IkappaBalpha promoter. Proc Natl Acad Sci USA. 107, 15193–15198 (2010).

73. I. S. Afonina, Z. Zhong, M. Karin, R. Beyaert, Limiting inflammation-the negative regulation of NF-κB and the NLRP3 inflammasome. Nat. Immunol. 18, 861–869 (2017).

74. L. Huang, Q. Yao, X. Gu, Q. Wang, L. Ren, Y. Wang, P. Hu, L. Guo, M. Liu, J. Xu, X. Zhang, Y. Qu, Y. Fan, X. Li, C. Li, T. Yu, J. Xia, M. Wei, L. Chen, Y. Li, B. Cao, 1-year outcomes in hospital survivors with COVID-19: a longitudinal cohort study. Lancet. 398, 747–758 (2021).

75. K. Kotfis, S. Williams Roberson, J. E. Wilson, W. Dabrowski, B. T. Pun, E. W. Ely, COVID-19: ICU delirium management during SARS-CoV-2 pandemic. Crit. Care. 24, 176 (2020).

76. C. Fang, J. Mei, H. Tian, Y.-L. Liou, D. Rong, W. Zhang, Q. Liao, N. Wu, CSF3 Is a Potential Drug Target for the Treatment of COVID-19. Front. Physiol. 11, 605792 (2020).

77. R. Agha, J. R. Avner, Delayed Seasonal RSV Surge Observed During the COVID-19 Pandemic. Pediatrics. 148 (2021), doi:10.1542/peds.2021-052089.

78. R. Song, Y. Gao, I. Dozmorov, V. Malladi, I. Saha, M. M. McDaniel, S. Parameswaran, C. Liang, C. Arana, B. Zhang, B. Wakeland, J. Zhou, M. T. Weirauch, L. C. Kottyan, E. K. Wakeland, C. Pasare, IRF1 governs the differential interferon-stimulated gene responses in human monocytes and macrophages by regulating chromatin accessibility. Cell Rep. 34, 108891 (2021).

79. G. Ren, K. Cui, Z. Zhang, K. Zhao, Division of labor between IRF1 and IRF2 in regulating different stages of transcriptional activation in cellular antiviral activities. Cell Biosci. 5, 17 (2015).

80. R. E. Niec, A. Y. Rudensky, E. Fuchs, Inflammatory adaptation in barrier tissues. Cell. 184, 3361–3375 (2021).

81. J. Friščić, M. Böttcher, C. Reinwald, H. Bruns, B. Wirth, S.-J. Popp, K. I. Walker, J. A. Ackermann, X. Chen, J. Turner, H. Zhu, L. Seyler, M. Euler, P. Kirchner, R. Krüger, A. B. Ekici, T. Major, O. Aust, D. Weidner, A. Fischer, M. H. Hoffmann, The complement system drives local inflammatory tissue priming by metabolic reprogramming of synovial fibroblasts. Immunity. 54, 1002–1021.e10 (2021).

82. J. Ordovas-Montanes, S. Beyaz, S. Rakoff-Nahoum, A. K. Shalek, Distribution and storage of inflammatory memory in barrier tissues. Nat. Rev. Immunol. 20, 308–320 (2020).

83. M. R. Corces, A. E. Trevino, E. G. Hamilton, P. G. Greenside, N. A. Sinnott-Armstrong, S. Vesuna, A. T. Satpathy, A. J. Rubin, K. S. Montine, B. Wu, A. Kathiria, S. W. Cho, M. R. Mumbach, A. C. Carter, M. Kasowski, L. A. Orloff, V. I. Risca, A. Kundaje, P. A. Khavari, T. J. Montine, H. Y. Chang, An improved ATAC-seq protocol reduces background and enables interrogation of frozen tissues. Nat. Methods. 14, 959–962 (2017).

84. T. B. Bennike, B. Fatou, A. Angelidou, J. Diray-Arce, R. Falsafi, R. Ford, E. E. Gill, S. D. van Haren, O. T. Idoko, A. H. Lee, R. Ben-Othman, W. S. Pomat, C. P. Shannon, K. K. Smolen, S. J. Tebbutt, A. Ozonoff, P. C. Richmond, A. H. J. van den Biggelaar, R. E. W. Hancock, B. Kampmann, H. Steen, Preparing for life: plasma proteome changes and immune system development during the first week of human life. Front. Immunol. 11, 578505 (2020).

85. S. E. Racine-Brzostek, J. K. Yee, A. Sukhu, Y. Qiu, S. Rand, P. D. Barone, Y. Hao, H. S. Yang, Q. H. Meng, F. S. Apple, Y. Shi, A. Chadburn, E. Golden, S. C. Formenti, M. M. Cushing, Z. Zhao, Rapid, robust, and sustainable antibody responses to mRNA COVID-19 vaccine in convalescent COVID-19 individuals. JCI Insight (2021).

86. A. M. Bolger, M. Lohse, B. Usadel, Trimmomatic: a flexible trimmer for Illumina sequence data. Bioinformatics. 30, 2114–2120 (2014).

87. H. Li, Aligning sequence reads, clone sequences and assembly contigs with BWA-MEM (2013).

88. Picard Tools - By Broad Institute, (available at https://broadinstitute.github.io/picard/).

89. H. Li, B. Handsaker, A. Wysoker, T. Fennell, J. Ruan, N. Homer, G. Marth, G. Abecasis, R. Durbin, 1000 Genome Project Data Processing Subgroup, The Sequence Alignment/Map format and SAMtools. Bioinformatics. 25, 2078–2079 (2009).

90. Y. Zhang, T. Liu, C. A. Meyer, J. Eeckhoute, D. S. Johnson, B. E. Bernstein, C. Nusbaum, R. M. Myers, M. Brown, W. Li, X. S. Liu, Model-based analysis of ChIP-Seq (MACS). Genome Biol. 9, R137 (2008).

91. Y. Liao, G. K. Smyth, W. Shi, featureCounts: an efficient general purpose program for assigning sequence reads to genomic features. Bioinformatics. 30, 923–930 (2014).

92. J. T. Robinson, H. Thorvaldsdóttir, W. Winckler, M. Guttman, E. S. Lander, G. Getz, J. P. Mesirov, Integrative genomics viewer. Nat. Biotechnol. 29, 24–26 (2011).

93. T. Stuart, A. Srivastava, S. Madad, C. A. Lareau, R. Satija, Single-cell chromatin state analysis with Signac. Nat. Methods. 18, 1333–1341 (2021).

94. A. Thibodeau, A. Eroglu, C. S. McGinnis, N. Lawlor, D. Nehar-Belaid, R. Kursawe, R. Marches, D. N. Conrad, G. A. Kuchel, Z. J. Gartner, J. Banchereau, M. L. Stitzel, A. E. Cicek, D. Ucar, AMULET: a novel read count-based method for effective multiplet detection from single nucleus ATAC-seq data. Genome Biol. 22, 252 (2021).

95. Y. Hao, S. Hao, E. Andersen-Nissen, W. M. Mauck, S. Zheng, A. Butler, M. J. Lee, A. J. Wilk, C. Darby, M. Zager, P. Hoffman, M. Stoeckius, E. Papalexi, E. P. Mimitou, J. Jain, A. Srivastava, T. Stuart, L. M. Fleming, B. Yeung, A. J. Rogers, R. Satija, Integrated analysis of multimodal single-cell data. Cell. 184, 3573–3587.e29 (2021).

96. S. L. Wolock, R. Lopez, A. M. Klein, Scrublet: Computational Identification of Cell Doublets in Single-Cell Transcriptomic Data. Cell Syst. 8, 281–291.e9 (2019).

97. I. Korsunsky, N. Millard, J. Fan, K. Slowikowski, F. Zhang, K. Wei, Y. Baglaenko, M. Brenner, P.-R. Loh, S. Raychaudhuri, Fast, sensitive and accurate integration of single-cell data with Harmony. Nat. Methods. 16, 1289–1296 (2019).

98. H. Zhao, Z. Sun, J. Wang, H. Huang, J.-P. Kocher, L. Wang, CrossMap: a versatile tool for coordinate conversion between genome assemblies. Bioinformatics. 30, 1006–1007 (2014).

99. C. Trapnell, D. Cacchiarelli, J. Grimsby, P. Pokharel, S. Li, M. Morse, N. J. Lennon, K. J. Livak, T. S. Mikkelsen, J. L. Rinn, The dynamics and regulators of cell fate decisions are revealed by pseudotemporal ordering of single cells. Nat. Biotechnol. 32, 381–386 (2014).

100. Z. Gu, R. Eils, M. Schlesner, Complex heatmaps reveal patterns and correlations in multidimensional genomic data. Bioinformatics. 32, 2847–2849 (2016).

101. J. Alquicira-Hernandez, J. E. Powell, Nebulosa recovers single cell gene expression signals by kernel density estimation. Bioinformatics (2021), doi:10.1093/bioinformatics/btab003.

102. Bioconductor - DiffBind, (available at https://bioconductor.org/packages/release/bioc/html/DiffBind.html).

103. G. Yu, L.-G. Wang, Q.-Y. He, ChIPseeker: an R/Bioconductor package for ChIP peak annotation, comparison and visualization. Bioinformatics. 31, 2382–2383 (2015).

104. D. Ucar, E. O. Karakaslar, cinaR: A comprehensive R package for the dif-ferential analyses and functional interpretation of ATAC-seq data. BioRxiv (2021), doi:10.1101/2021.03.05.434143.

105. M. D. Robinson, D. J. McCarthy, G. K. Smyth, edgeR: a Bioconductor package for differential expression analysis of digital gene expression data. Bioinformatics. 26, 139–140 (2010).

106. Ready-to-Use Curated Gene Sets for “cinaR” [R package cinaRgenesets version 0.1.1] (2021), (available at https://cran.r-project.org/web/packages/cinaRgenesets/index.html).

107. S. Heinz, C. Benner, N. Spann, E. Bertolino, Y. C. Lin, P. Laslo, J. X. Cheng, C. Murre, H. Singh, C. K. Glass, Simple combinations of lineage-determining transcription factors prime cis-regulatory elements required for macrophage and B cell identities. Mol. Cell. 38, 576–589 (2010).

108. U. Raudvere, L. Kolberg, I. Kuzmin, T. Arak, P. Adler, H. Peterson, J. Vilo, g:Profiler: a web server for functional enrichment analysis and conversions of gene lists (2019 update). Nucleic Acids Res. 47, W191–W198 (2019).

109. Z. Li, M. H. Schulz, T. Look, M. Begemann, M. Zenke, I. G. Costa, Identification of transcription factor binding sites using ATAC-seq. Genome Biol. 20, 45 (2019).

110. S. Kurtenbach, J. W. Harbour, SparK: A Publication-quality NGS Visualization Tool. BioRxiv (2019), doi:10.1101/845529.

111. F. Ramírez, F. Dündar, S. Diehl, B. A. Grüning, T. Manke, deepTools: a flexible platform for exploring deep-sequencing data. Nucleic Acids Res. 42, W187–91 (2014).

112. A. N. Schep, B. Wu, J. D. Buenrostro, W. J. Greenleaf, chromVAR: inferring transcription-factor-associated accessibility from single-cell epigenomic data. Nat. Methods. 14, 975–978 (2017).

113. F. J. Ibarrondo, J. A. Fulcher, D. Goodman-Meza, J. Elliott, C. Hofmann, M. A. Hausner, K. G. Ferbas, N. H. Tobin, G. M. Aldrovandi, O. O. Yang, Rapid Decay of Anti-SARS-CoV-2 Antibodies in Persons with Mild Covid-19. N. Engl. J. Med. 383, 1085–1087 (2020).

114. E. H. Y. Lau, O. T. Y. Tsang, D. S. C. Hui, M. Y. W. Kwan, W.-H. Chan, S. S. Chiu, R. L. W. Ko, K. H. Chan, S. M. S. Cheng, R. A. P. M. Perera, B. J. Cowling, L. L. M. Poon, M. Peiris, Neutralizing antibody titres in SARS-CoV-2 infections. Nat. Commun. 12, 63 (2021).

115. W. H. Self, M. W. Tenforde, W. B. Stubblefield, L. R. Feldstein, J. S. Steingrub, N. I. Shapiro, A. A. Ginde, M. E. Prekker, S. M. Brown, I. D. Peltan, M. N. Gong, M. S. Aboodi, A. Khan, M. C. Exline, D. C. Files, K. W. Gibbs, C. J. Lindsell, T. W. Rice, I. D. Jones, N. Halasa, IVY Network, Decline in SARS-CoV-2 Antibodies After Mild Infection Among Frontline Health Care Personnel in a Multistate Hospital Network - 12 States, April-August 2020. MMWR Morb Mortal Wkly Rep. 69, 1762–1766 (2020).

116. E. Marklund, S. Leach, H. Axelsson, K. Nyström, H. Norder, M. Bemark, D. Angeletti, A. Lundgren, S. Nilsson, L.-M. Andersson, A. Yilmaz, M. Lindh, J.-Å. Liljeqvist, M. Gisslén, Serum-IgG responses to SARS-CoV-2 after mild and severe COVID-19 infection and analysis of IgG non-responders. PLoS ONE. 15, e0241104 (2020).

117. K. K.-W. To, O. T.-Y. Tsang, W.-S. Leung, A. R. Tam, T.-C. Wu, D. C. Lung, C. C.-Y. Yip, J.-P. Cai, J. M.-C. Chan, T. S.-H. Chik, D. P.-L. Lau, C. Y.-C. Choi, L.-L. Chen, W.-M. Chan, K.-H. Chan, J. D. Ip, A. C.-K. Ng, R. W.-S. Poon, C.-T. Luo, V. C.-C. Cheng, K.-Y. Yuen, Temporal profiles of viral load in posterior oropharyngeal saliva samples and serum antibody responses during infection by SARS-CoV-2: an observational cohort study. Lancet Infect. Dis. 20, 565–574 (2020).

