## Supplemental Figures for "Epigenetic Memory of COVID-19 in Innate Immune Cells and Their Progenitors"

**This PDF file includes:**

Materials and Methods
Supplementary Text
Figs. S1 to S9
Table. S1
Captions for Data S1 to S3

**Other Supplementary Materials for this manuscript include the following:**

Data S1 to S3 (Excel)

### Materials and Methods

Study Cohorts

A total of 168 study participants were enrolled at Weill-Cornell Medicine/New York-Presbyterian Hospital between March 2020 and March 2021. Participants were recruited from the inpatient division of New York-Presbyterian Hospital and the Weill-Cornell Medicine pulmonary and post-ICU clinics. No statistical methods were used to predetermine sample size. COVID-19 severity scoring was based on the COVID-19 World Health Organization (WHO) Severity Classification [https://www.who.int/blueprint/priority-diseases/key-action/COVID-19_Treatment_Trial_Design_Master_Protocol_synopsis_Final_18022020.pdf]. Subjects were binned into the following groups: 1) healthy volunteer donors, 2) recovered mild COVID-19 patients (WHO score 1-2), 3) recovered severe COVID-19 patients (WHO score 6-7), 4) recovered non-COVID-19 critically ill patients. Inclusion criteria for each group were as follows; 1) healthy volunteer donors: absence of clinical COVID-19 symptoms at any time prior to blood collection (prior negative SARS-CoV2 PCR and/or seronegative status also considered when available), 2) recovered mild COVID-19 patients: PCR-proven SARS-CoV2 infection with the presence of clinical COVID-19 symptoms not requiring hospitalization, 3) severe COVID-19 patients: PCR-proven SARS-CoV2 infection with the presence of clinical COVID-19 symptoms requiring admission to ICU-level care and the use of mechanical ventilation, 4) recovered non-COVID19 critically ill patients: absence of SARS-CoV2 infection as measured by PCR and negative serology on admission and/or throughout hospital admission and non-COVID-19 related critical illness requiring admission to the medical, neurological, or cardiology intensive care unit. Prior infection status in healthy volunteer donors and recovered mild COVID-19 groups was confirmed by SARS-CoV2 serological testing after donation. There were no specific exclusion criteria other than an inability to provide informed consent or SARS-CoV2 positive serology in asymptomatic healthy volunteers (asymptomatic infection) and non-COVID19 critically ill participants. Blood was collected in EDTA or sodium heparin-coated vacutainers and kept on gentle agitation until processing, and all blood was processed on the day of collection. Age, sex, and comorbidity data were obtained through EPIC EHR records or when unavailable through a standardized form at the time of donation.

PBMC and Plasma isolation

Whole blood from EDTA or Heparin tubes (BD 366643 and BD 368480, respectively) was spun at 500g for 10 minutes at room temperature with no brake. The undiluted plasma was aliquoted to 1.5 ml microcentrifuge tubes and stored at –80° C for subsequent analysis.

After removal of plasma, the blood was mixed at a 1:1 ratio with room temperature RPMI medium (Corning 10-040-CM), layered over Ficoll-Paque PLUS (GE 17144002), and spun at 700g for 30 minutes at room temperature with minimum acceleration and no brake. The PBMC layer was isolated and washed with RPMI. Cells were then treated with ACK lysis buffer for 3 minutes and counted on a Countess 2 automated cell counter (Thermo Fisher AMQAX1000). Cells were centrifuged again and resuspended in freezing medium (90% FBS + 10% DMSO) and stored in cryogenic vials in a freezing container (Thermo Fisher 5100-0001) at –80° C.

CD34$^{+}$ and CD14$^{+}$ cell isolation

Frozen PBMCs were thawed in a 37° C water bath, washed with RPMI, and centrifuged. An aliquot of PBMCs was stained with 7-AAD (Biolegend 420404, 1:20) alone. The rest of the cells were incubated with CD34 microbeads (Miltenyi 130-046-702) and isolated by placing them on a magnetic column (Miltenyi 130-042-201) as per the manufacturer's specifications. The positive fraction obtained from the magnetic column was stained with the following antibodies – CD34-FITC (Miltenyi 130-113-178, 1:100), CD49f-Pacific Blue (Biolegend 313620, 1:200), CD90-PE (Biolegend 328110, 1:100), CD38-PE/cy7 (Biolegend 303516, 1:100), CD45RA-APC/cy7 (Biolegend 304128, 1:400), Lineage markers (CD20-Biotin {Biolegend 302350, 1:100}, CD16-Biotin {Biolegend 302004, 1:100}, CD3-Biotin {Biolegend 344820, 1:100}, CD56-Biotin {Biolegend 362536, 1:100}, and CD14-Biotin {Biolegend 301826, 1:100}), and 7-AAD (Biolegend 420404, 1:20). After incubating in the dark for 30 minutes, cells were washed with PBS and incubated with Streptavidin-BV605 (BD 563260, 1:500) for an additional 30 minutes. CD34+ cells from the positive fraction and bulk PBMCs from the PBMC aliquot were then sorted on a BD FACSAria cell sorter and mixed at 1:5-1:20 ratios.

The negative fraction from the magnetic column was stained with the following antibodies – CD14-APC (BD 340436, 1:1000), CD8-FITC (Biolegend 300906, 1:400), and 7-AAD (Biolegend 420404, 1:20). After incubating in the dark for 30 minutes, cells were washed with PBS, and CD14+ cells were sorted on a BD FACSAria cell sorter.

ATAC-seq $CD34^+$ HSPC and $CD14^+$ Monocytes

To perform ATAC-seq, we followed the Omni-ATAC-seq protocol (*81*). We used 50,000 cells for $CD14^+$ monocytes, and 3000~5000 for $CD34^+$ HSPC. HSPC were sorted directly into the PCR tubes prior to following Omni-ATAC-seq protocol.

Single-cell Assays

Nuclei were isolated from a mix of CD34+ cells and PBMCs according to 'Low Cell Input Nuclei Isolation' protocol (10x Genomics CG000365-Rev B) and were processed using Chromium Controller & Next GEM Accessory Kit (10x Genomics 1000202) and Chromium Next GEM Single Cell Multiome ATAC + Gene Expression Reagent Bundle (10x Genomics 1000285) following the manufacturer's User Guide (10x Genomics CG000338-Rev D). Targeted nuclei recovery ranged from 5,000 to 10,000. The single-cell RNA and ATAC sequencing libraries were prepared using Dual Index Kit TT Set A (10x Genomics 1000215) and Single Index Kit N Set A (10x Genomics 1000212) respectively and sequenced on Illumina NovaSeq6000 or NextSeq platform.

Plasma protein analyses

The frozen plasma was thawed, aliquoted, and either analyzed immediately or re-frozen and shipped for analysis. All analysis was performed on samples after their first freeze-thaw cycle.

Mass Spectrometry

MS/MS was performed as described previously (*82*). Briefly, 5 μL plasma was diluted in 100 μL sample buffer (8 M urea in TRIS-HCl, pH 8.5). Protein disulfide bonds were reduced with dithiothreitol (10 mM final concentration) and alkylated with iodoacetamide (50 mM final concentration). An aliquot with 10 μg protein was transferred to a 96-well plate with polyvinylidene fluoride (PVDF) membrane bottom (MSIPS4510, Millipore, MA, USA). Protein digestion was performed with sequencing-grade modified trypsin (V5111, Promega, Madison, WI, USA) at a nominal protease to protein ratio of 1:25 w/w. After incubation for 2 h at 37°C, the peptides were eluted and concentrated to dryness in a vacuum centrifuge. To monitor retention time stability and system performance, iRT peptides (Biognosys, Schlieren, Switzerland) were spiked into all samples. The samples were analyzed using a nanoLC system (Eksigent, Dublin, CA) equipped with an LCchip system (cHiPLC nanoflex, Eksigent, CA, USA) coupled online to a Q Exactive Mass Spectrometer (Thermo Scientific, Bremen, Germany).

Cytokine analysis

Plasma was shipped to Eve Technologies for their 15-plex human pro-inflammatory cytokine assay (Eve Technologies, Calgary, AB, Canada). All samples were analyzed in duplicate.

Antibody assay

The SARS-CoV-2 total RBD antibody (TAb), surrogate neutralizing antibody (SNAb), and avidity were used to measure plasma antibody levels on the TOP-Plus (Pylon 3D analyzer; ET Healthcare) as previously described (*83*).

Bulk ATAC-seq data processing

ATAC-seq paired-end sequences (n= 54 for sorted CD14+, n= 47 for sorted CD34+) were trimmed using trimmomatic (*84*) and the trimmed reads were aligned to the GRCh37 (hg19) genome using bwa (*85*). Further processing steps were carried out on the aligned data using Picard(*86*) and Samtools (*87*). MACS2 (*88*) was used for peak calling on the processed data using the BAMPE option with default parameters. For quality control, FRiP scores were calculated for all samples using the featurecounts program available in the Subread package (*89*).

Samples with low FRiP scores (< 0.15) were removed from downstream analysis. To ensure that the samples being removed were indeed poor quality, we conducted visual inspections using IGV (*90*). After FRiP score filtering, we were left with 27 CD14+ samples and 13 CD34+ samples.

Single-cell ATAC-seq data processing

Two single-cell ATAC-seq samples were preprocessed using the Cell Ranger ATAC 1.2.0 pipeline and aligned to the GRCh38 (hg38) genome. The cellranger output was processed using Signac (*91*). Individual samples were filtered out (Supplementary Table 1 for the details of QC cutoffs). Amulet (*92*) was used for filtering out doublets. Post QC and doublet removal, the remaining steps of the Signac pipeline (TF-IDF normalization, SVD, UMAP embedding, and clustering) were completed. UMAP embedding and clustering were done using 30 PCs. The cells were annotated by using a reference PBMC scRNAseq dataset with Seurat's anchor transfer functions.

Multiome data processing

The Multiome data (ATAC + RNA) (n=33) were preprocessed using the Cell Ranger ARC 1.0.0 pipeline and aligned to the hg38 genome. The cellranger output was then processed using the Seurat Weighted Nearest Neighbor Pipeline (*93*). Low QC cells were filtered out (data S2 for details). Doublets were removed using Amulet (*92*) for ATACseq and Scrublet (*94*) for RNAseq. Initial annotations of cells were carried out using the reference PBMC CITE-seq data in the Seurat package.

The Multiome data from different individuals were pooled using Seurat and Signac for RNA-seq and ATAC-seq, respectively. Three samples were excluded from the pooling process due to poor quality based on the initial clustering. Low QC cells from individual samples were filtered out from the pooled data. For the RNA-seq object, sctransform normalization was applied, followed by PCA, and 30 PCs were used for UMAP embedding and clustering. The merged dataset was also batch corrected using Harmony (95) with all samples being used as a batch, and the UMAP embedding and clustering were repeated using 20 PCs. The RNA object was then annotated using the reference PBMC CITE-seq object in Seurat.

We then pooled scATAC profiles from all samples and did the first round of cell-type annotation based on scRNA-seq annotations. Then we called peaks on each cell type using MACS2 (version 2.1.2) with the following parameters: `callpeak --nomodel --nolambda --keep-dup all --call-summits.` Peak summits from all cell types were combined, extended on both sides by 150 bp. Redundant peaks were removed based on the q-value from MACS2. Using the generated peak region list, the number of reads overlapping a given peak window was determined for each unique cell barcode tag. This generated a peak by cell counts matrix corresponding to ATAC reads in peaks for each cell profiled. High-quality cells are retained with a fraction of reads in peaks (FRiP)>0.4 and sequencing depth > 1000. The cells filtered out in this step were also removed from the scRNA object to ensure the same cells were retained across both modalities.

After QC, the scRNA-seq object was reprocessed using sctranform, PCA, clustering, UMAP, Harmony, and PBMC CITE-seq reference annotation. In the end, 23 clusters were obtained from the scRNA-seq data using default parameters of Seurat (30 PCs for PCA). Annotations of these clusters were finalized based on the expression of marker genes for distinct immune cell types. 2 of these clusters were labeled as potential doublets and removed from downstream analyses since they expressed marker genes of more than one immune cell type. In the end, 21 clusters were retained that included 197,260 cells out of 260K cells in total in the RNA object. The same cells were then retained in the scATAC-seq component of the Multiome data, and the annotations were transferred. Once the cell type annotations for major cell types are finalized, we repeated the peak calling using MACS2 and the aforementioned parameter settings, which are used in the downstream DORC analyses using the ArchR pipeline. The scATAC merged object was also processed using the Signac pipeline with TF-IDF normalization, SVD, UMAP embedding, and clustering. We used Harmony (as part of the Signac pipeline) for batch correction in scATAC data using individual samples as a batch. For both non-corrected and Harmony-batch-corrected UMAP embeddings and clustering, 30 PCs were used.

Pseudobulk profiles

The cells that were annotated as $CD14^+$ monocytes (n=34) and HSPC (n=34) from the Multiome ATAC-seq data and single-cell ATAC-seq data were used to create 'pseudobulk' profiles for these cell types using snATACClusteringTools. (https://github.com/UcarLab/snATACClusteringTools). For each sample, we generated bam files for these two cell types. 2 samples were filtered out based on low cell numbers (HDu and jcov124) (only for $CD14^+$), three samples were from the same individual and pooled together (lgtd17, lgtd18, and lgtd19), two samples were filtered out due to clinical reasons (jcov114 from $CD14^+$ only, and jcov49_2 from $CD14^+$ and $CD34^+$), resulting in 28 samples for monocytes and 31 for HSPC. In HSPC, The pseudobulk bam files were converted into hg19 using Crossmap (*96*). Peak calling was done using MACS2 (*88*) with the BAMPE option.

HSPC annotations

While we annotated subclusters of RNA-seq HSPC data based on the expression of manually curated marker genes in each cluster, the ATAC-seq data was separately annotated for the HSPC subtypes using a previously reported bulk-guided approach (46). Briefly, we first lift-over the reference bulk ATAC peak set from hg19 to hg38. Next, we generated a peak by cell counts matrix using the lifted peak set. We identified 27 principal components (PCs) of variation in reference bulk ATAC-seq samples, then scored every single cell by the contribution of each PC. Cells were subsequently clustered using the Euclidean distance between these normalized single-cell PCs scores and PCs of bulk samples. The annotations based on the bulk ATAC dataset were then transferred to the scRNAseq object based on identical cellular identities.

Integration of HSPC data from BM and PBMC-PIE.

To transfer HSPC subtype annotations from public data to our HSPC data, we used public scRNA-seq data of BMMC from healthy participants and the Seurat package (92). The data available from GEO with access code GSE139369. Seurat object was created using BMMC data, normalized, then anchors for transferring annotation was defined by FindTransferAnchors() function of Seurat package using BMMD data as a reference. BMMC annotation information was transferred via anchors to our HSPC data using TransferData(). After transferring annotation was done, we merged those two Seurat objects using merge() function. We used Harmony (95) for batch correction by the source of the data.

Peak-gene cis-association and DORC identification

To calculate peak-gene associations, we used our previously published approach. (*51*) We considered all the peaks that are located in the +/-50 kb window around annotated TSSs. We used peak counts and imputed gene expression to calculate the observed Spearman correlation (obs) of each peak-gene pair. To estimate the background, we generated 100 background peaks for each peak by matching accessibility and GC content (chromVAR) (104) and calculated the Spearman correlation coefficient between those background peaks and the gene, resulting in a null peak-gene Spearman correlation distribution. We then calculated the expected population mean (pop.mean) and expected population standard deviation (pop.sd) from expected Spearman correlations. The Z score is calculated by z=(obs-pop.mean)/pop.sd. For peaks associated with multiple genes, we only kept peak-gene associations with the smallest p-value.

To define DORCs, we selected genes with at least 8 peaks per gene. The DORC score was calculated at each DORC gene for each cell. We defined the DORC score by summing up all the significantly correlated peak counts per gene per cell. We then normalize the DORC score by dividing the DORC score by the total unique fragments in peaks and obtain a cell x DORC score matrix.

Pseudotime Analysis

We generated Seurat object for HSPC and myeloid cell types(CD14+ monocytes, CD16+ monocytes, DC1, and DC2) with DORC score matrix then reprocessed it as we did for RNA-seq data by using PCA, clustering, UMAP, Harmony (*95*), Cell type annotations from RNA-seq data were used for the DORC data. The object was then converted to CellDataSet(CDS) format, using as.cell_data_set() in SeuratWrappers package. Pseudotime trajectories were constructed with

Monocle3 (*97*). In order to align cells on the pseudotime, we chose one node, which is close to the center of CD164 expressing cells, as a root node using order_cells() function. Pseudotime values given to each cell was used for various downstream analysis and visualization. Subset pseudotime was obtained by assigning two nodes for the start point and endpoint.

Using the subset trajectory, we measured the DORC activity of each cell sorted by pseudotime. The DORC-associated genes were grouped by hierarchical clustering using the ComplexHeatmap package (*98*). Clusters of DORC-associated genes with increasing or decreasing trends throughout the pseudotime were defined as Myeloid or HSPC modules, respectively. DORC activity for genes within the monocyte and HSPC modules were row-normalized across all single cells, and a "cluster score" was computed based on the average of these row-normalized scores. HSPC and Monocyte cluster scores were plotted and visualized using a locally estimated scatterplot smoothing approach.

Visualizing cohort density on UMAP plots

With the processed Seurat RNA-seq object, we extracted shared nearest neighbor graph information (*93*). Using this data, we determined the 50 nearest cells of each cell, then calculated the frequency of each cohort. We then included cohort frequency information of each cell in the metadata to visualize it on a UMAP plot using Nebulosa (*99*).

UMAP visualization

UMAP plots to show clusters, cohorts, samples, and annotations were generated using DimPlot() function in the Seurat package (*93*). For visualizing features such as gene expression, motif activity, cohort density, we used Nebulosa (*99*).

Differential Analysis

For differential accessibility (DA) analyses, good quality bulk and pseudobulk ATAC-seq samples for $CD14^{+}$ monocytes (n=55) and HSPC (n=44) were used to generate consensus peak sets using DiffBind (*100*) by retaining peaks that are detected at least in 2 samples, resulting in 106,504 consensus peaks for Monocytes and 110,610 peaks for HSPC. jcov114 was removed from the Consensus Peakset before Differential Analysis for HSPC, leaving us with 43 samples in total. Consensus peaks were annotated using ChipSeeker (*101*) and the peaks were filtered out based on 3 criteria - CPM threshold (=1), library threshold (=2), and distance to TSS threshold (<50 KB). Post filtering, we had 79,713 consensus peaks for monocytes and 82,148 peaks for HSPC. These peaks were then used to identify differentially accessible regions among clinical groups using the cinaR R package. For differential accessibility analyses, we used GLM models as implemented in EdgeR (*102*, *103*) by conducting pairwise comparisons among the clinical groups. Due to the variability of ages across clinical groups, we used age as a covariate. Finally, to account for known and unknown batches, we used Surrogate Variable Analysis using all significant SVs. Differential peaks at FDR 10% were kept for downstream analyses. Consensus peaks that are not called in the single-cell object were discarded from downstream analyses leaving us with 75596 peaks for Monocytes and 79205 peaks for CD34.

Differential expression (DE) analyses were carried out using the pooled single-cell RNA object (n=30) using Seurat's FindMarkers function by conducting pairwise comparisons among clinical groups for each detect cell type. The normalized RNA counts matrix was used for DE analyses using the non-parametric Wilcoxon rank sum test with a minimum fold change value of 0.25 and minimum percentage of cells for feature detection at 10%. Differential genes with an adjusted p-value < 0.1 were kept.

DORC-RNA correlation

To correlate DORC and RNA-seq data, we performed differential expression analysis between two groups using the DORC score matrix and RNA-seq normalized data for 968 DORC-associated genes. log2FC of expression was obtained by using Seurat's FindMarkers() function, as well as adjusted p-value. (*93*) We performed a t-test between two groups of cells using the DORC score matrix using the t.test() function of R to obtain the difference between the mean of each group and p-values for each gene. Estimates of difference (log2FC and mean difference) between two groups were correlated for individual DORC associated genes.

Functional enrichment analyses

The peaks were annotated, including the closest genes using cinaR (*104*) and ChIPSeeker (*101*). The TSS regions were defined as -3KB to 3KB. In cases of overlap, the following order of genomic annotations is used: Promoter > 5' UTR > 3' UTR > Exon > Intron > Downstream (defined as downstream of gene end) > Intergenic. Functional enrichment for the differential peak sets was conducted using hypergeometric gene set enrichment tests followed by Benjamini-Hochberg FDR adjustment for P-values (cutoff adjusted p=0.1). For functional enrichment, we used different curated gene sets from the CinaRgenesets package (*104*). For HSPC, Gene Set Enrichment Analysis (GSEA) was carried out in addition to the hypergeometric testing since DA peak counts were smaller.

HOMER (*105*) was used for further functional enrichments for KEGG, GO, REACTOME, etc. (p-value cutoff=0.1). HOMER was also used to identify TF motifs enriched in differential peak sets using default settings both for known and de novo motifs. For motif enrichment analyses, we used the default (whole genome) as a background.

Gene Ontology analysis

All Gene ontology terms stated in the study were identified with web-based g:Profiler (*106*).

Time Series/Trend Analysis -

Differential Peaks from bulk and pseudobulk ATAC-seq data were pooled together for the following 4 comparisons: S:2-4mo vs. Healthy, S:4-12mo vs. Healthy, M:2-4mo vs. Healthy and nonCoV v.s Healthy for both $CD14^+$ Monocytes and HSPC. These peaks were then used to detect trends across different clinical groups using tcseq. The read counts and log fold change values associated with the differential peaks for each clinical group were used as input, and the clinical groups served as time points for the purpose of our analysis. A various number of clusters were tested before we chose 4 clusters as the ideal number to visualize the different patterns in the clinical groups. The differential peaks were then split into 4 clusters, and the Functional Enrichment and Motif Analysis using HOMER (*105*) were conducted for each of the individual clusters.

HINT

Footprints are more reliable with deeply sequenced data. To increase the read depth of our data, we pooled Pseudobulk ATAC profiles from each clinical group into a single profile for both

CD14 Monocytes and HSPC. Peaks were then called for these pooled profiles using macs2. The HINT (Hmm-based IdeNtification of Transcription factor footprints) framework (*107*) was then used to identify active transcription factor binding sites for each clinical group. We also used HINT to find motifs overlapping with the footprints using the JASPAR database for TF motifs analyzed in Fig. 5. We then used HINT to generate average ATAC-seq profiles around binding sites of transcription factors of interest. HINT was also used to calculate the differential changes in TF activity between different clinical groups.

The motif regions overlapping with footprints were annotated using ChipSeeker (*101*), and based on those annotations, hypergeometric geneset enrichment was carried out for the footprinting regions using various genetic datasets. The enrichment results were adjusted using the Benjamini-Hochberg FDR adjustment method (FDR = 10%).

Spark Tracks

We also visuzelid Pseudobulk profiles using a package called SparK (*108*). Firstly, bedgraph files were generated using deepTools (*109*) with the following command 'bamCoverage -b bamfile.bam -o outputfilename.bdg -bs 1 -of bedgraph'. Spark was then used to generate genome browser track figures using the bedgraph files for selected loci.

scATAC Motif analysis

Motif enrichment in the pooled snATAC-seq dataset was conducted using two methods. Before conducting the enrichment analysis, motif information was added to the pooled object using the AddMotifs function in Signac (*91*). Motif information for hg38 was added to the object from the JASPAR2020 database.

For the first approach, we calculated overrepresented motifs in a set of differentially accessible peaks. Differential accessibility (DA) analyses were carried out using the pooled single-cell ATAC object (n=30) using Seurat's FindMarkers function (*93*) by conducting pairwise comparisons among clinical groups for each cell type. The normalized ATAC counts matrix was used for DA analyses using a logistic regression framework with a minimum fold change value of 0.25 and minimum percentage of cells for feature detection at 10%. Differential peaks with an adjusted p-value < 0.05 were kept. We then used these top differential peaks to find overrepresented motifs using the FindMotifs function, which uses a hypergeometric test to find overrepresented motifs in a set of genomic features.

For the second approach, we performed differential motif activity analysis between different sets of cells. Firstly, we computed a per cell motif activity score using chromVAR (*110*) and added this information to the pooled ATAC object as a separate assay. We then conducted Differential Motif Analysis using the FindMarkers function as mentioned previously. For this analysis, the Motif activity scores were used for the Differential Analysis using the non-parametric Wilcoxon rank sum test with a minimum fold change value of 0.25 and minimum percentage of cells for feature detection at 10%.

To visualize the chromVAR score as a heatmap, we first took the mean of each TF chromVAR score for each cell. Then median was taken for each cohort, followed by Z-score normalization. We used select TF families to visualize the heatmap.

We also performed footprinting for a smaller select set of transcription factors using the Footprint function in Signac. This calculated the footprinting information of the motifs for every instance in the genome using the whole genome (Hg38) as a background.

### Supplementary Text

Characterization of Plasma

Antibody titers and antibody quality remained elevated from early to late convalescence following severe COVID-19 (S:2-4mo, S:4-12mo). In contrast, as reported (*111–115*), mild COVID-19 resulted in lower initial titers (M:2-4mo) that declined over time (M:4-12mo) (fig. S1A). Consistent with a distinguishing milieu and duration of inflammation, we observed the persistence of elevated plasma IL-6 in critical illness in general and severe COVID-19 (S:2-4mo and nonCoV) (fig. S1C). MCP-1, IL-10, IL-1ra were also elevated in S:2-4mo subjects, and IL-10 was unique as the only persistently elevated factor in S:4-12mo compared to healthy subjects (fig. S1C). For unbiased identification and quantification of plasma factors, we performed LC-MS/MS analysis of plasma proteins across cohorts. Early in convalescence, both S:2-4mo and nonCOV cohorts (characterized by severe disease) featured enrichment of active plasma response,s including an abundance of acute-phase response factors and platelet degranulation, clotting, and neovascularization factors (SAA1, ORM1, LBP, fibrinogen-FGG/FGA, LRG1). Plasma factors uniquely elevated in the S:2-4mo cohort included immune complement factors (C1R, C1QA, C3, C4A) and risk factors for atherosclerosis (APOC3 and LPA) (fig. S1B). With immune complement signaling recently implicated in local tissue inflammatory priming (*79*) it will be important to understand the clinical implications of months-long and aberrantly high immune complement activity following COVID-19 (compared to general critical illness) and its potential contribution to persistent tissue inflammation, fibrosis, and symptoms.

Referencesa


83. M. R. Corces, A. E. Trevino, E. G. Hamilton, P. G. Greenside, N. A. Sinnott-Armstrong, S. Vesuna, A. T. Satpathy, A. J. Rubin, K. S. Montine, B. Wu, A. Kathiria, S. W. Cho, M. R. Mumbach, A. C. Carter, M. Kasowski, L. A. Orloff, V. I. Risca, A. Kundaje, P. A. Khavari, T. J. Montine, H. Y. Chang, An improved ATAC-seq protocol reduces background and enables interrogation of frozen tissues. *Nat. Methods*. **14**, 959–962 (2017).

84. T. B. Bennike, B. Fatou, A. Angelidou, J. Diray-Arce, R. Falsafi, R. Ford, E. E. Gill, S. D. van Haren, O. T. Idoko, A. H. Lee, R. Ben-Othman, W. S. Pomat, C. P. Shannon, K. K. Smolen, S. J. Tebbutt, A. Ozonoff, P. C. Richmond, A. H. J. van den Biggelaar, R. E. W. Hancock, B. Kampmann, H. Steen, Preparing for life: plasma proteome changes and immune system development during the first week of human life. *Front. Immunol.* **11**, 578505 (2020).

85. S. E. Racine-Brzostek, J. K. Yee, A. Sukhu, Y. Qiu, S. Rand, P. D. Barone, Y. Hao, H. S. Yang, Q. H. Meng, F. S. Apple, Y. Shi, A. Chadburn, E. Golden, S. C. Formenti, M. M. Cushing, Z. Zhao, Rapid, robust, and sustainable antibody responses to mRNA COVID-19 vaccine in convalescent COVID-19 individuals. *JCI Insight* (2021).

86. A. M. Bolger, M. Lohse, B. Usadel, Trimmomatic: a flexible trimmer for Illumina sequence data. *Bioinformatics*. **30**, 2114–2120 (2014).

87. H. Li, Aligning sequence reads, clone sequences and assembly contigs with BWA-MEM (2013).

88. Picard Tools - By Broad Institute, (available at https://broadinstitute.github.io/picard/).

89. H. Li, B. Handsaker, A. Wysoker, T. Fennell, J. Ruan, N. Homer, G. Marth, G. Abecasis, R. Durbin, 1000 Genome Project Data Processing Subgroup, The Sequence Alignment/Map format and SAMtools. *Bioinformatics*. **25**, 2078–2079 (2009).

90. Y. Zhang, T. Liu, C. A. Meyer, J. Eeckhoute, D. S. Johnson, B. E. Bernstein, C. Nusbaum, R. M. Myers, M. Brown, W. Li, X. S. Liu, Model-based analysis of ChIP-Seq (MACS). *Genome Biol.* **9**, R137 (2008).

91. Y. Liao, G. K. Smyth, W. Shi, featureCounts: an efficient general purpose program for assigning sequence reads to genomic features. *Bioinformatics*. **30**, 923–930 (2014).

92. J. T. Robinson, H. Thorvaldsdóttir, W. Winckler, M. Guttman, E. S. Lander, G. Getz, J. P. Mesirov, Integrative genomics viewer. *Nat. Biotechnol.* **29**, 24–26 (2011).

93. T. Stuart, A. Srivastava, S. Madad, C. A. Lareau, R. Satija, Single-cell chromatin state analysis with Signac. *Nat. Methods*. **18**, 1333–1341 (2021).

94. A. Thibodeau, A. Eroglu, C. S. McGinnis, N. Lawlor, D. Nehar-Belaid, R. Kursawe, R. Marches, D. N. Conrad, G. A. Kuchel, Z. J. Gartner, J. Banchereau, M. L. Stitzel, A. E. Cicek, D. Ucar, AMULET: a novel read count-based method for effective multiplet detection from single nucleus ATAC-seq data. *Genome Biol.* **22**, 252 (2021).

95. Y. Hao, S. Hao, E. Andersen-Nissen, W. M. Mauck, S. Zheng, A. Butler, M. J. Lee, A. J. Wilk, C. Darby, M. Zager, P. Hoffman, M. Stoeckius, E. Papalexi, E. P. Mimitou, J. Jain,

A. Srivastava, T. Stuart, L. M. Fleming, B. Yeung, A. J. Rogers, R. Satija, Integrated analysis of multimodal single-cell data. *Cell*. **184**, 3573-3587.e29 (2021).

96. S. L. Wolock, R. Lopez, A. M. Klein, Scrublet: Computational Identification of Cell Doublets in Single-Cell Transcriptomic Data. *Cell Syst.* **8**, 281-291.e9 (2019).

97. I. Korsunsky, N. Millard, J. Fan, K. Slowikowski, F. Zhang, K. Wei, Y. Baglaenko, M. Brenner, P.-R. Loh, S. Raychaudhuri, Fast, sensitive and accurate integration of single-cell data with Harmony. *Nat. Methods*. **16**, 1289–1296 (2019).

98. H. Zhao, Z. Sun, J. Wang, H. Huang, J.-P. Kocher, L. Wang, CrossMap: a versatile tool for coordinate conversion between genome assemblies. *Bioinformatics*. **30**, 1006–1007 (2014).

99. C. Trapnell, D. Cacchiarelli, J. Grimsby, P. Pokharel, S. Li, M. Morse, N. J. Lennon, K. J. Livak, T. S. Mikkelsen, J. L. Rinn, The dynamics and regulators of cell fate decisions are revealed by pseudotemporal ordering of single cells. *Nat. Biotechnol.* **32**, 381–386 (2014).

100. Z. Gu, R. Eils, M. Schlesner, Complex heatmaps reveal patterns and correlations in multidimensional genomic data. *Bioinformatics*. **32**, 2847–2849 (2016).

101. J. Alquicira-Hernandez, J. E. Powell, Nebulosa recovers single cell gene expression signals by kernel density estimation. *Bioinformatics* (2021), doi:10.1093/bioinformatics/btab003.

102. Bioconductor - DiffBind, (available at https://bioconductor.org/packages/release/bioc/html/DiffBind.html).

103. G. Yu, L.-G. Wang, Q.-Y. He, ChIPseeker: an R/Bioconductor package for ChIP peak annotation, comparison and visualization. *Bioinformatics*. **31**, 2382–2383 (2015).

104. D. Ucar, E. O. Karakaslar, cinaR: A comprehensive R package for the dif-ferential analyses and functional interpretation of ATAC-seq data. *BioRxiv* (2021), doi:10.1101/2021.03.05.434143.

105. M. D. Robinson, D. J. McCarthy, G. K. Smyth, edgeR: a Bioconductor package for differential expression analysis of digital gene expression data. *Bioinformatics*. **26**, 139–140 (2010).

106. Ready-to-Use Curated Gene Sets for “cinaR” [R package cinaRgenesets version 0.1.1] (2021), (available at https://cran.r-project.org/web/packages/cinaRgenesets/index.html).

107. S. Heinz, C. Benner, N. Spann, E. Bertolino, Y. C. Lin, P. Laslo, J. X. Cheng, C. Murre, H. Singh, C. K. Glass, Simple combinations of lineage-determining transcription factors prime cis-regulatory elements required for macrophage and B cell identities. *Mol. Cell*. **38**, 576–589 (2010).

108. U. Raudvere, L. Kolberg, I. Kuzmin, T. Arak, P. Adler, H. Peterson, J. Vilo, g:Profiler: a web server for functional enrichment analysis and conversions of gene lists (2019 update). *Nucleic Acids Res.* **47**, W191–W198 (2019).

109. Z. Li, M. H. Schulz, T. Look, M. Begemann, M. Zenke, I. G. Costa, Identification of transcription factor binding sites using ATAC-seq. *Genome Biol.* **20**, 45 (2019).

110. S. Kurtenbach, J. W. Harbour, SparK: A Publication-quality NGS Visualization Tool. *BioRxiv* (2019), doi:10.1101/845529.

111. F. Ramírez, F. Dündar, S. Diehl, B. A. Grüning, T. Manke, deepTools: a flexible platform for exploring deep-sequencing data. *Nucleic Acids Res.* **42**, W187-91 (2014).

112. A. N. Schep, B. Wu, J. D. Buenrostro, W. J. Greenleaf, chromVAR: inferring transcription-factor-associated accessibility from single-cell epigenomic data. *Nat. Methods*. **14**, 975–978 (2017).

113. F. J. Ibarrondo, J. A. Fulcher, D. Goodman-Meza, J. Elliott, C. Hofmann, M. A. Hausner, K. G. Ferbas, N. H. Tobin, G. M. Aldrovandi, O. O. Yang, Rapid Decay of Anti-SARS-CoV-2 Antibodies in Persons with Mild Covid-19. *N. Engl. J. Med.* **383**, 1085–1087 (2020).

114. E. H. Y. Lau, O. T. Y. Tsang, D. S. C. Hui, M. Y. W. Kwan, W.-H. Chan, S. S. Chiu, R. L. W. Ko, K. H. Chan, S. M. S. Cheng, R. A. P. M. Perera, B. J. Cowling, L. L. M. Poon, M. Peiris, Neutralizing antibody titres in SARS-CoV-2 infections. *Nat. Commun.* **12**, 63 (2021).

115. W. H. Self, M. W. Tenforde, W. B. Stubblefield, L. R. Feldstein, J. S. Steingrub, N. I. Shapiro, A. A. Ginde, M. E. Prekker, S. M. Brown, I. D. Peltan, M. N. Gong, M. S. Aboodi, A. Khan, M. C. Exline, D. C. Files, K. W. Gibbs, C. J. Lindsell, T. W. Rice, I. D.

Jones, N. Halasa, IVY Network, Decline in SARS-CoV-2 Antibodies After Mild Infection Among Frontline Health Care Personnel in a Multistate Hospital Network - 12 States, April-August 2020. *MMWR Morb Mortal Wkly Rep*. **69**, 1762–1766 (2020).

116. E. Marklund, S. Leach, H. Axelsson, K. Nyström, H. Norder, M. Bemark, D. Angeletti, A. Lundgren, S. Nilsson, L.-M. Andersson, A. Yilmaz, M. Lindh, J.-Å. Liljeqvist, M. Gisslén, Serum-IgG responses to SARS-CoV-2 after mild and severe COVID-19 infection and analysis of IgG non-responders. *PLoS ONE*. **15**, e0241104 (2020).

117. K. K.-W. To, O. T.-Y. Tsang, W.-S. Leung, A. R. Tam, T.-C. Wu, D. C. Lung, C. C.-Y. Yip, J.-P. Cai, J. M.-C. Chan, T. S.-H. Chik, D. P.-L. Lau, C. Y.-C. Choi, L.-L. Chen, W.-M. Chan, K.-H. Chan, J. D. Ip, A. C.-K. Ng, R. W.-S. Poon, C.-T. Luo, V. C.-C. Cheng, K.-Y. Yuen, Temporal profiles of viral load in posterior oropharyngeal saliva samples and serum antibody responses during infection by SARS-CoV-2: an observational cohort study. *Lancet Infect. Dis.* **20**, 565–574 (2020).

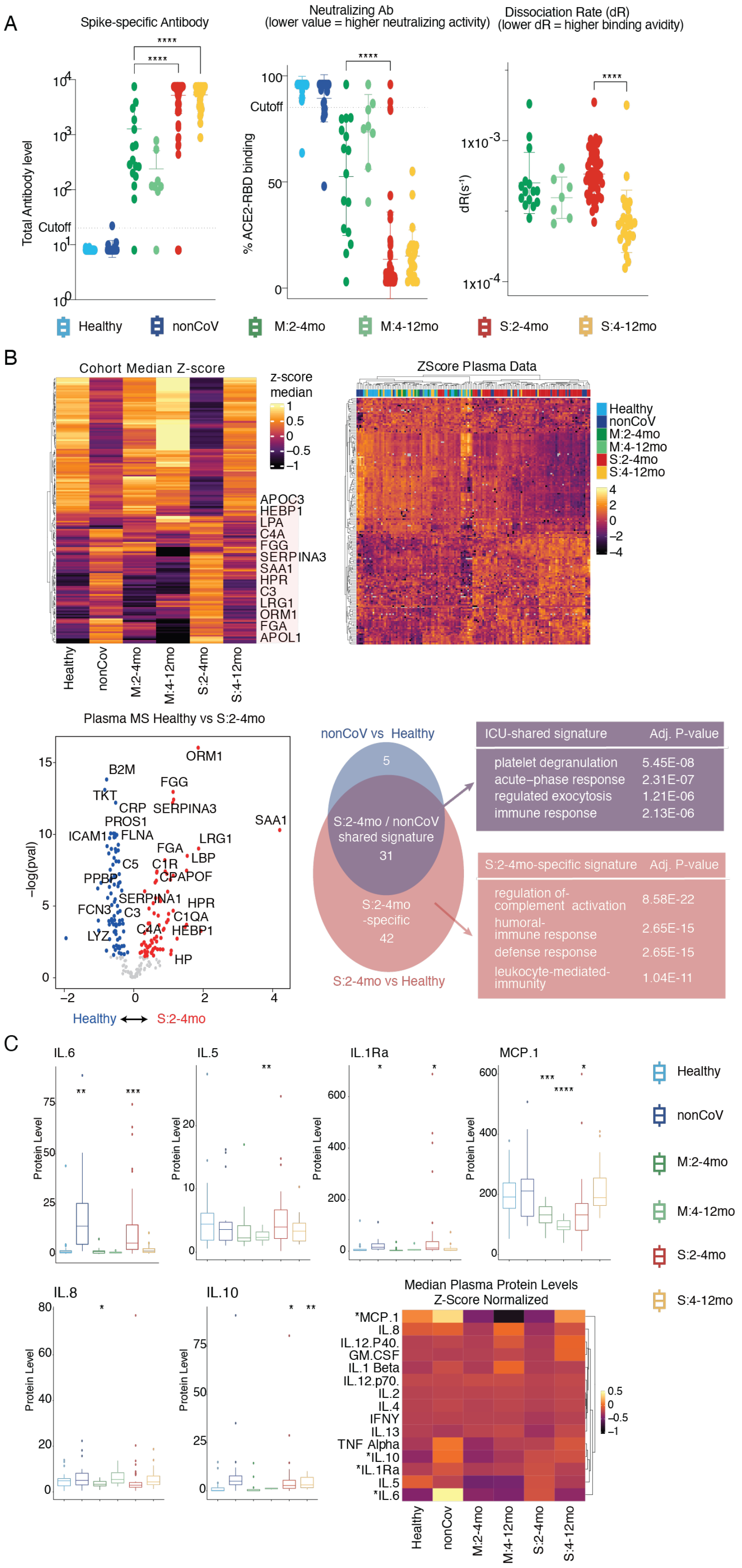
A
Spike-specific Antibody
Neutralizing Ab
(lower value = higher neutralizing activity)
Dissociation Rate (dR)
(lower dR = higher binding avidity)
Total Antibody level
Cutoff
% ACE2-RBD binding
dR(s-1)
Healthy
nonCoV
M:2-4mo
M:4-12mo
S:2-4mo
S:4-12mo
B
Cohort Median Z-score
z-score median
APOC3
HEBP1
LPA
C4A
FGG
SERPINA3
SAA1
HPR
C3
LRG1
ORM1
FGA
APOL1
ZScore Plasma Data
Plasma MS Healthy vs S:2-4mo
ORM1
B2M
FGG
TKT
CRP
SERPINA3
PROS1
SAA1
ICAM1
FLNA
LRG1
FGA
LBP
C5
C1R
PPBP
CPAPOF
SERPINA1
HPR
FCN3
C3
C1QA
C4A
HEBP1
LYZ
HP
−log(pval)
Healthy ⟷ S:2-4mo
nonCoV vs Healthy
S:2-4mo / nonCoV shared signature
S:2-4mo -specific
S:2-4mo vs Healthy
ICU-shared signature
Adj. P-value
platelet degranulation 5.45E-08
acute−phase response 2.31E-07
regulated exocytosis 1.21E-06
immune response 2.13E-06
S:2-4mo-specific signature
Adj. P-value
regulation of- complement activation 8.58E-22
humoral- immune response 2.65E-15
defense response 2.65E-15
leukocyte-mediated- immunity 1.04E-11
C
IL.6
IL.5
IL.1Ra
MCP.1
IL.8
IL.10
Protein Level
Median Plasma Protein Levels
Z-Score Normalized
*MCP.1
IL.8
IL.12.P40.
GM.CSF
IL.1 Beta
IL.12.p70.
IL.2
IL.4
IFNY
IL.13
TNF Alpha
*IL.10
*IL.1Ra
IL.5
*IL.6

**Fig. S1. Characterization of plasma. (A)** Viral S protein-specific antibody assays showing antibody level, neutralizing activity, and binding quality. (ANOVA, p*<0.05) **(B)** Plasma protein levels measured by LC-MS. Top, left: hierarchical clustering of cohort average protein levels in plasma with proteins of interest (increased expression in post-COVID-19 cohorts) labeled; top, right: individual study participant plasma protein levels; bottom, left: volcano plot comparing plasma proteins levels between the healthy cohort and early convalescent severe (S:2-4mo) cohort, with log2-fold change (x-axis) and p-val (y-axis); bottom, right: venn diagram of differential (vs healthy) plasma factors identified in comparison with nonCov and S:2-4mo. **(C)** Heatmap and boxplots showing plasma cytokine levels measured by Luminex platform. (*p<0.05, t-test, Healthy group as a reference) For heatmap, median cytokine level per cohort was normalized by Z-score. Cytokines with any significant difference between healthy and any clinical cohorts are indicated by asterisk.

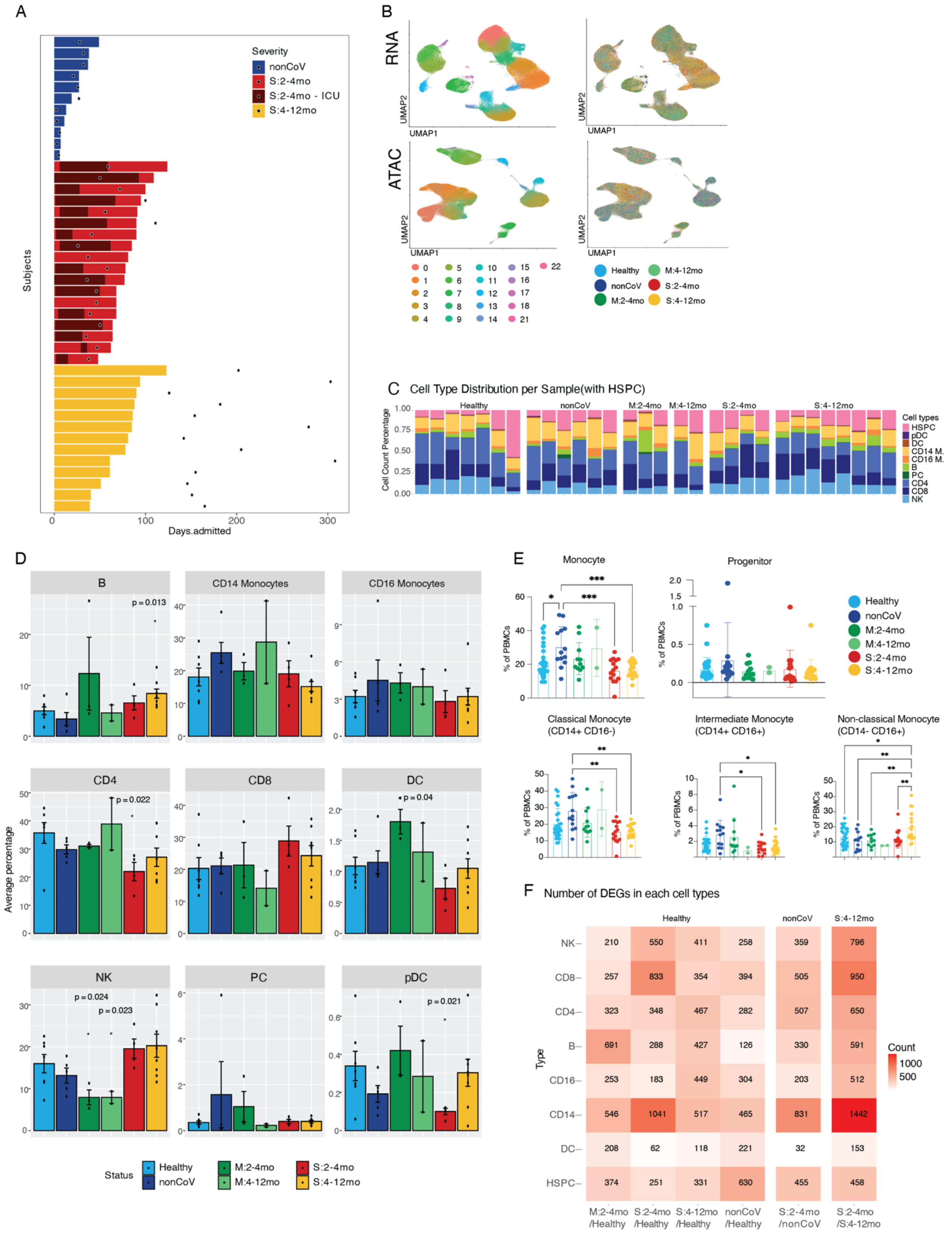


**Fig. S2. Characterization of cohort samples used for single-cell analysis.**

**(A)** Swimmer plot showing hospitalization periods, and sample collection time point for each patient in nonCOV, S(2-4mo), and S(4-12mo). **(B)** UMAP plots displaying unbiased clustering, sample and cohort distribution using snRNA-seq (RNA) and snATAC-seq (ATAC) data across all samples. **(C)** Stacked bar plot showing cell type composition in individual patient samples. **(D)** Barplot showing frequencies of each cell type across cohorts. Enriched HSPC population was removed from the frequency calculation. p values indicate significance relative to healthy controls, t-test. **(E)** Flow cytometry results showing frequencies of CD14+ monocytes, CD34+ HSPC, and monocyte subsets across cohorts. (ANOVA, * $p<0.05$) **(F)** Count matrix showing number of differentially expressed genes in various comparisons across cell types. B cell (B), CD4 T cell (CD4), CD8 T cell (CD8), dendritic cell (DC), natural killer cell (NK), plasma cell (PC), plasmacytoid DC (pDC).

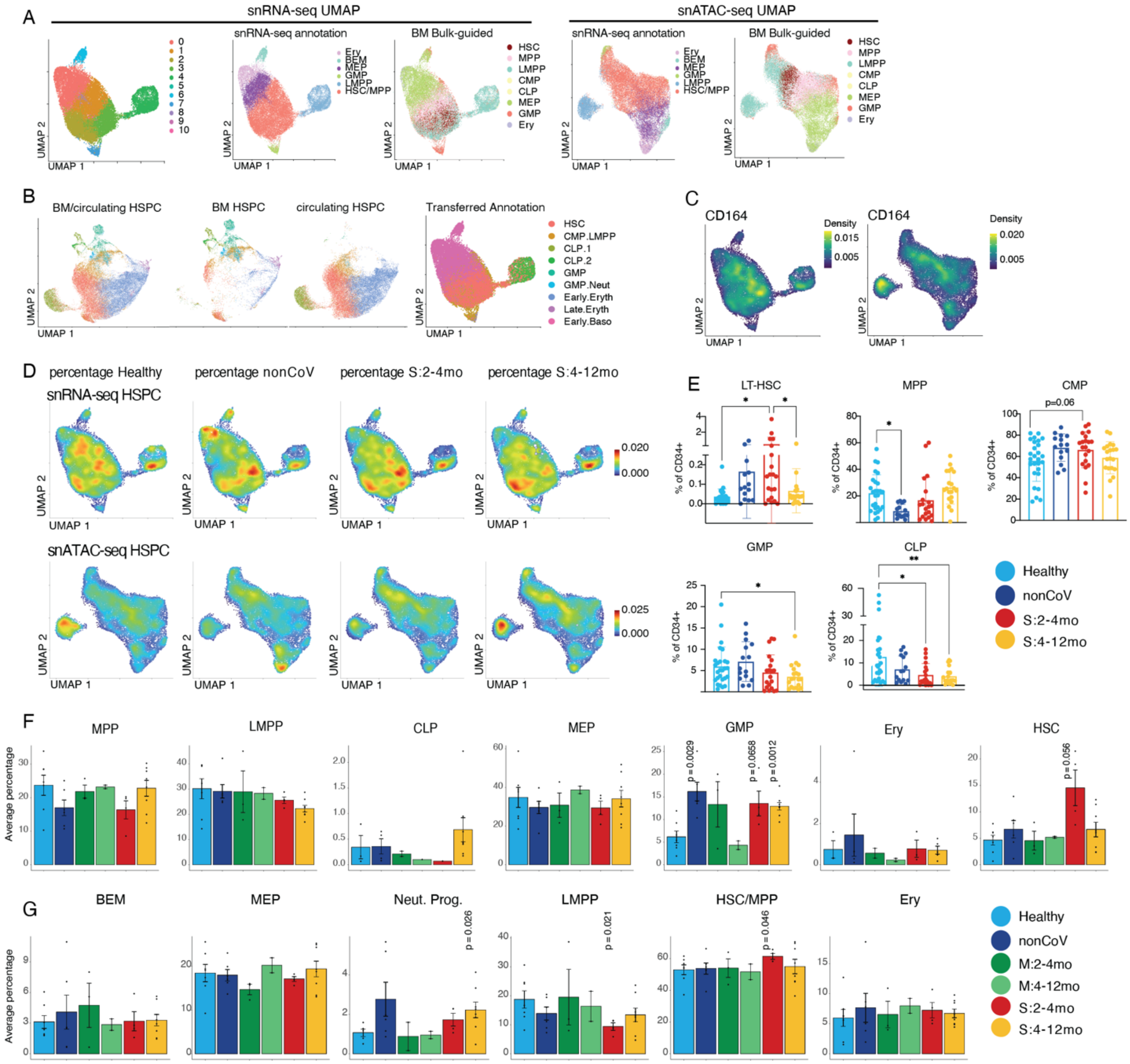


**Fig. S3. Characterization of HSPC used for single cell data.**

**(A)** Unbiased clustering of HSPC using snRNA-seq data (left), our HSPC annotation (middle), and bone marrow HSPC ATAC-seq-guided annotations (right) projected on snRNA-seq UMAP and snATAC-seq UMAP **(B)** UMAP representation of merged bone marrow (BM) HSPC scRNA-seq data (REF) and our circulating HSPC snRNA-seq data. Annotation is transferred from BM HSPC data. **(C)** Feature plots showing CD164 expressing cells on single-cell RNA/ATAC-seq HSPC UMAP plots. **(D)** Density projections for each cohort displayed on single-cell RNA/ATAC-seq HSPC UMAP plots. **(E)** Flow cytometry results showing frequencies of HSPC cell subsets across cohorts (ANOVA, *p <0.05) (see methods for HSPC flow panel). **(F)** Boxplots showing frequencies of each HSPC bulk-guided subclusters across cohorts (t-test, *p<0.05, Healthy group as a reference). **(G)** Boxplots showing frequencies of each HSPC subtypes annotated based on manually curated genes across cohorts.

B cells (B), CD14+ monocytes (CD14 M.), CD16+ monocytes (CD16 M.), CD4+ T cells (CD4), CD8+ T cells (CD8), dendritic cell (DC), hematopoietic stem and progenitor cells (HSPC), natural killer cells (NK), plasma B cells (PC), plasmacytoid dendritic cells (pDC), erythroid progenitor cells (Ery), neutrophil progenitor cells (Neut. Pro.), basophil-eosinophil-mast cell progenitor cells (BEM), lymphoid-primed multipotent progenitor cells (LMPP), megakaryocyte-erythroid progenitor cells (MEP), hematopoietic stem cells/multipotent progenitor cells (HSC/MPP), common lymphoid progenitor cells (CLP), granulocyte-macrophage progenitor cells (GMP), common myeloid progenitor cells (CMP)

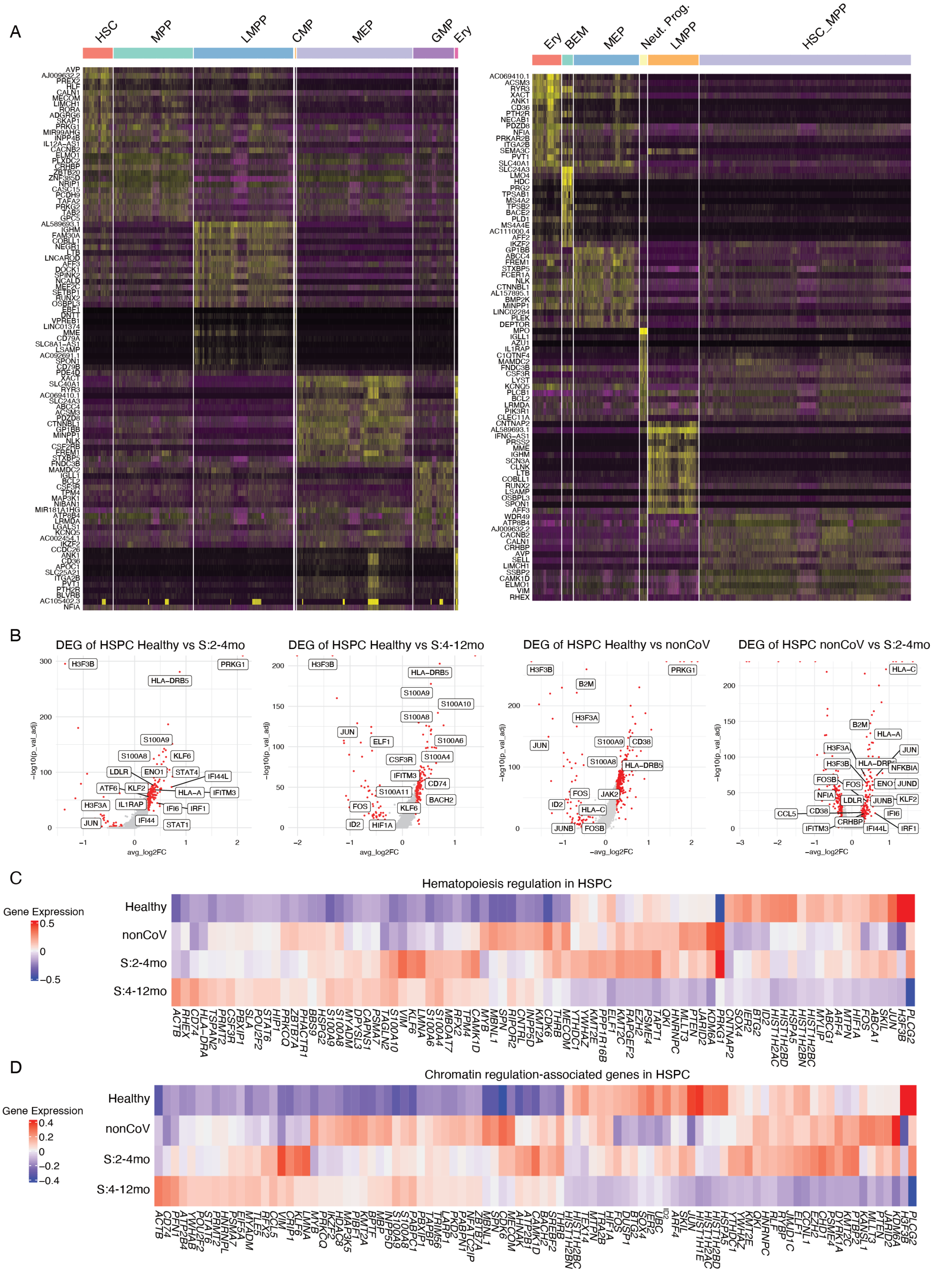
A
HSC
MPP
LMPP
CMP
MEP
GMP
Ery
BEM
Neut. Prog.
HSC_MPP
B
DEG of HSPC Healthy vs S:2-4mo
DEG of HSPC Healthy vs S:4-12mo
DEG of HSPC Healthy vs nonCoV
DEG of HSPC nonCoV vs S:2-4mo
-log10(p_val_adj)
avg_log2FC
-avg_log2FC
C
Hematopoiesis regulation in HSPC
Gene Expression
Healthy
nonCoV
S:2-4mo
S:4-12mo
D
Chromatin regulation-associated genes in HSPC
Gene Expression
Healthy
nonCoV
S:2-4mo
S:4-12mo

**Fig. S4. Transcriptomic signature of HSPC by sub-populations and clinical groups.**

**(A)** Heatmaps showing expression of top marker genes in each HSPC subpopulation are annotated based on bulk-guided annotation(left) and our manually curated genes for annotation(right). **(B)** Volcano plots showing differentially expressed genes in post-COVID19 subjects compared to healthy subjects. **(C-D)** Heatmap showing differentially expressed genes associated with hematopoietic regulation(top) and chromatin regulation(bottom) in healthy, convalescent severe and nonCoV subjects.

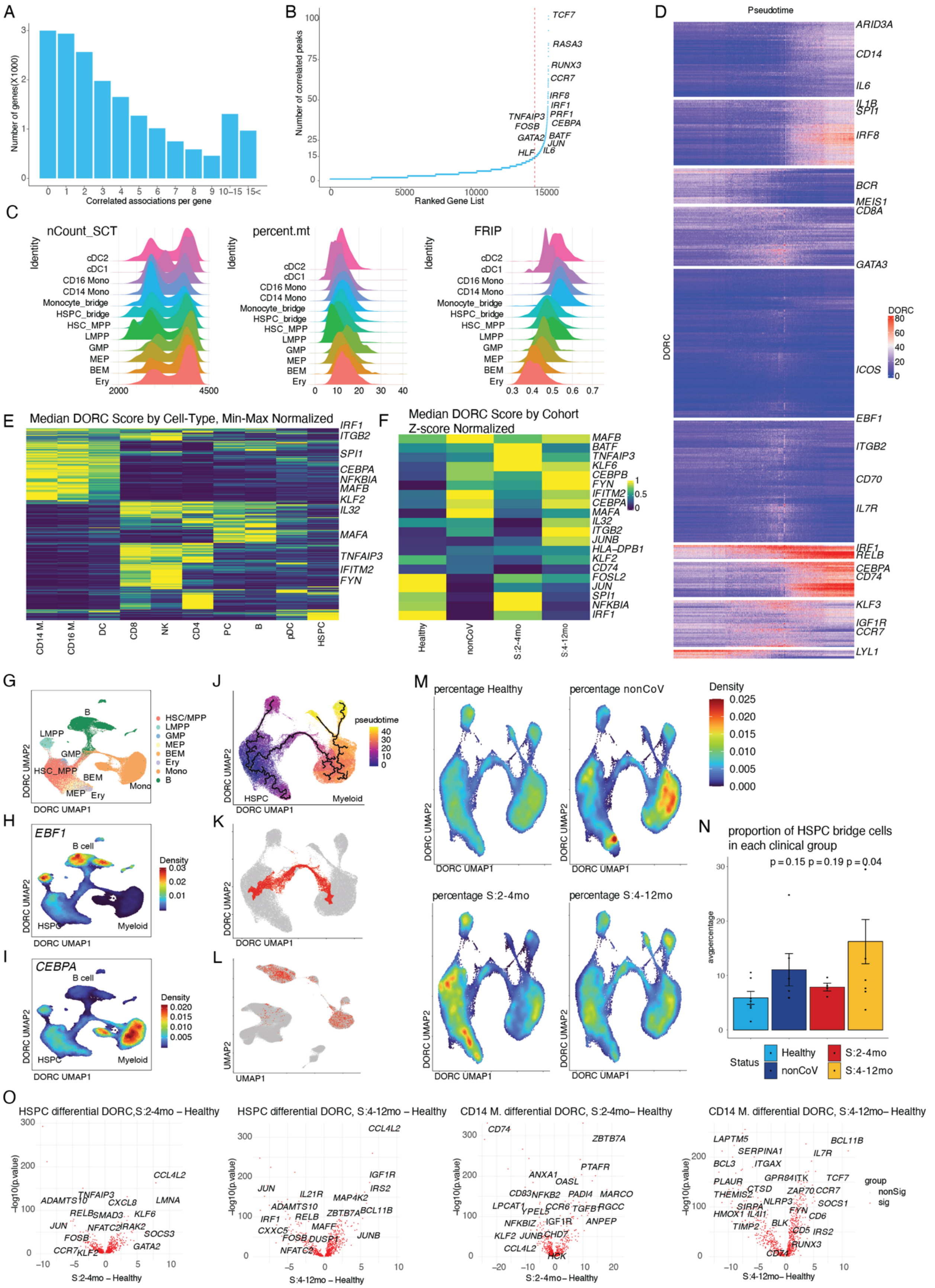

A
Number of genes(X1000)
Correlated associations per gene
0 1 2 3 4 5 6 7 8 9 10–15 15<
B
Number of correlated peaks
Ranked Gene List
TCF7
RASA3
RUNX3
CCR7
IRF8
IRF1
PRF1
CEBPA
TNFAIP3
FOSB
GATA2
BATF
JUN
HLF
IL6
C
nCount_SCT
percent.mt
FRIP
Identity
cDC2
cDC1
CD16 Mono
CD14 Mono
Monocyte_bridge
HSPC_bridge
HSC_MPP
LMPP
GMP
MEP
BEM
Ery
D
Pseudotime
DORC
ARID3A
CD14
IL6
IL1B
SPI1
IRF8
BCR
MEIS1
CD8A
GATA3
ICOS
EBF1
ITGB2
CD70
IL7R
IRF1
RELB
CEBPA
CD74
KLF3
IGF1R
CCR7
LYL1
E
Median DORC Score by Cell-Type, Min-Max Normalized
IRF1
ITGB2
SPI1
CEBPA
NFKBIA
MAFB
KLF2
IL32
MAFA
TNFAIP3
IFITM2
FYN
CD14 M.
CD16 M.
DC
CD8
NK
CD4
PC
B
pDC
HSPC
F
Median DORC Score by Cohort
Z-score Normalized
MAFB
BATF
TNFAIP3
KLF6
CEBPB
FYN
IFITM2
CEBPA
MAFA
IL32
ITGB2
JUNB
HLA-DPB1
KLF2
CD74
FOSL2
JUN
SPI1
NFKBIA
IRF1
Healthy
nonCoV
S:2-4mo
S:4-12mo
G
HSC/MPP
LMPP
GMP
MEP
BEM
Ery
Mono
B
DORC UMAP1
DORC UMAP2
J
pseudotime
HSPC
Myeloid
H
EBF1
B cell
Density
HSPC
Myeloid
K
I
CEBPA
B cell
Density
HSPC
Myeloid
L
UMAP1
UMAP2
M
percentage Healthy
percentage nonCoV
percentage S:2-4mo
percentage S:4-12mo
Density
N
proportion of HSPC bridge cells
in each clinical group
p = 0.15 p = 0.19 p = 0.04
avgpercentage
Status
Healthy
nonCoV
S:2-4mo
S:4-12mo
O
HSPC differential DORC,S:2-4mo – Healthy
HSPC differential DORC, S:4-12mo – Healthy
CD14 M. differential DORC, S:2-4mo– Healthy
CD14 M. differential DORC, S:4-12mo– Healthy
-log10(p.value)
S:2-4mo – Healthy
S:4-12mo – Healthy
S:2-4mo– Healthy
S:4-12mo– Healthy
group
nonSig
sig

**Fig. S5. Characterization of DORCs in HSPC and CD14+ monocytes related to Fig. 2**

**(A)** The number of significant peak-gene connections for all genes. **(B)** The number of significantly correlated peaks (p<0;05) for each gene. **(C)** Distribution of normalized count number, percentage of mitochondrial RNA, and FRIP score per cell across cell types annotated from HSPC-$CD14^+$ monocyte DORC UMAP plot. **(D)** Full heatmap showing activity of all DORC via pseudotime. **(E)** Heatmap showing DORC activity in individual cell types **(F)** Heatmap showing DORC activity of selected inflammatory gene-associated DORCs in HSPC across cohorts. **(G)** UMAP visualization of HSPC subsets, $CD14^+$ monocytes, $CD16^+$ monocytes, dendritic cells (DC), and B cells (B), using DORC score matrix. **(H)** DORC scores of Early B-Cell Factor 1 (EBF1) projected on the UMAP, showing expected distribution. **(I)** DORC scores of CEBPA projected on the UMAP, showing expected distribution in myeloid cells. **(J)** UMAP visualization of HSPC and myeloid cell types using DORC with trajectories obtained from Monocle3. Colors indicate pseudotime value given to cells. **(K-L)** HSPC and CD14+ monocytes DORC UMAP (K), and all-cell snRNA-seq UMAP (L) highlighting cells used for pseudotime analysis in Figure 2H. **(M)** Cohort density projected on the HSPC-$CD14^+$ monocytes DORC UMAP. **(N)** Boxplot showing cohort density on the HSPC bridge. (P values as indicated, t-test) **(O)** Volcano plots for differential activities of DORCs in HSPC and $CD14^+$ monocytes.

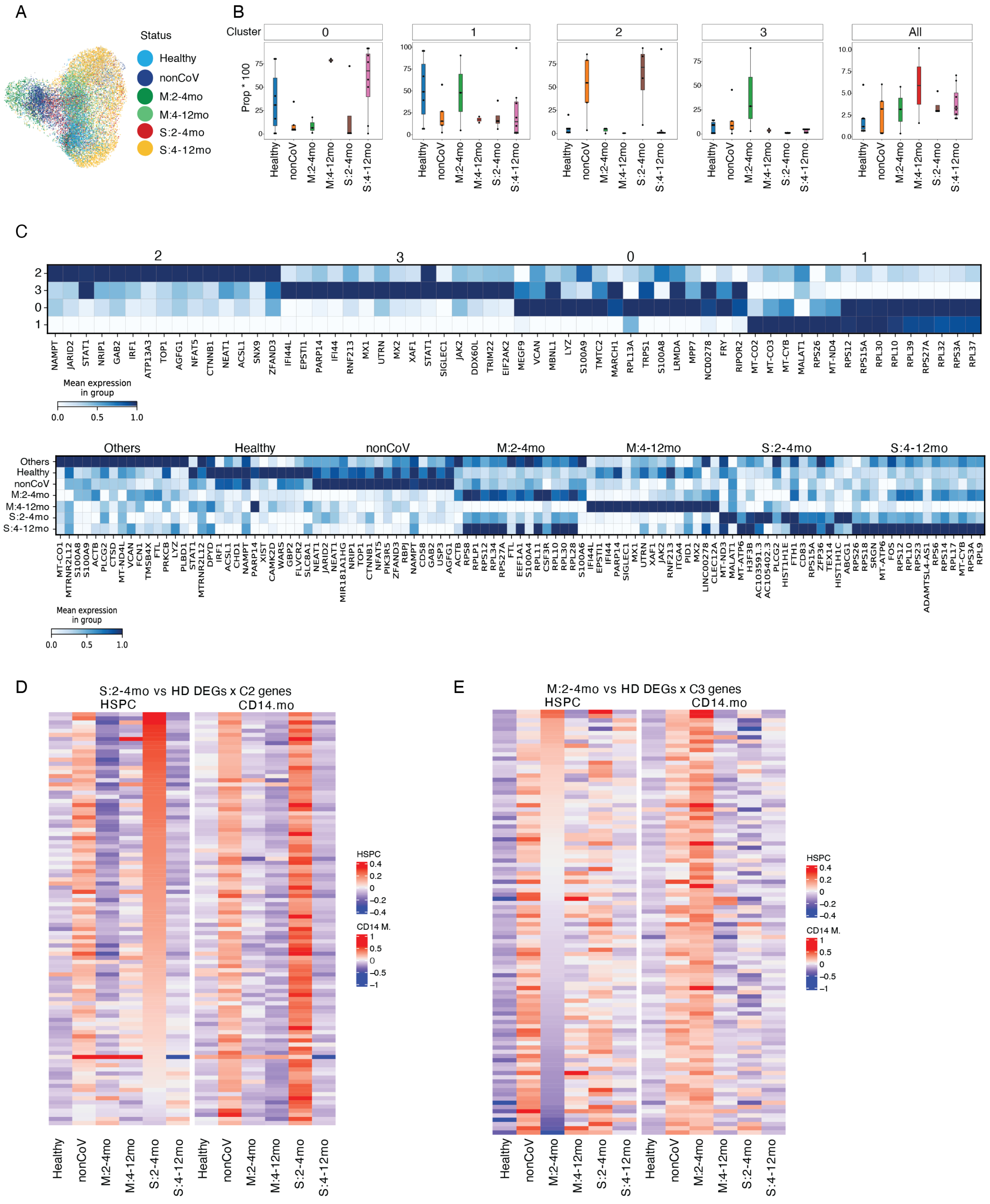


**Fig. S6. Characterization of subclusters of CD14+ monocytes related to Fig. 3.**

**(A)** CD14$^+$ monocytes UMAP plot colored by clinical cohorts. **(B)** Boxplot comparing the proportion of each of the four CD14$^+$ monocyte subclusters (Fig. 3A) across the individual and clinical cohorts. **(C)** Heat map representing scaled expression values of the top 15 genes defining: (i) the CD14$^+$ monocyte subclusters (upper panel) or (ii) clinical cohorts (lower panel). **(D-E)** Expression heatmaps for differentially expressed SC2 and SC3-specific genes in HSPC and CD14+ monocytes.

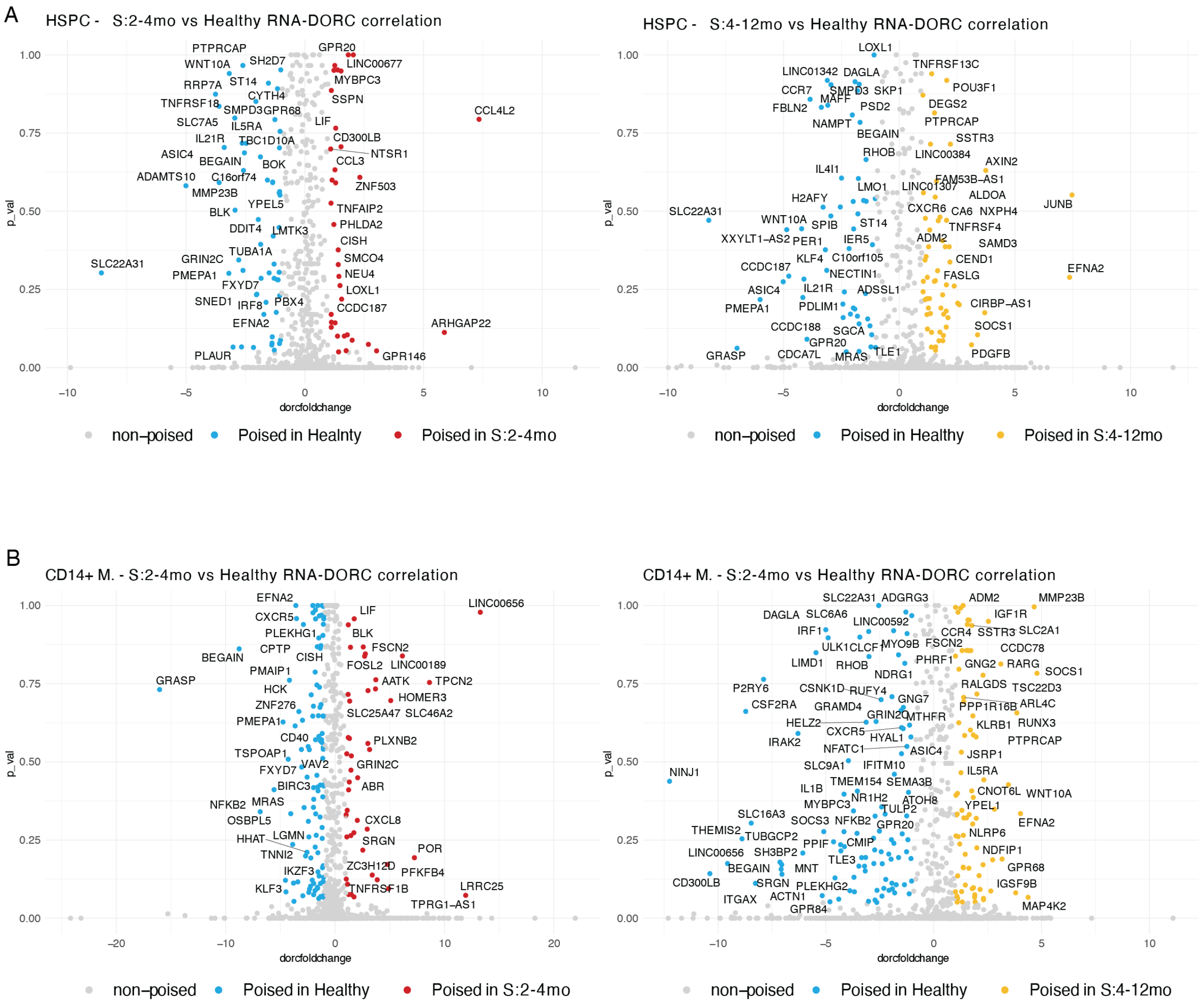


**Fig. S7. Poised genes in Severe post-COVID-19 groups.**

**(A-B)** Scatter plots showing poised genes in HSPC (A) and CD14$^{+}$ monocytes (B) in S:2-4mo (left) and S4-12mo (right) when compared to healthy. These genes have significant changes in DORC activity but not in gene expression levels. (DORC: $p < 0.05$, adjusted $p < 0.05$)

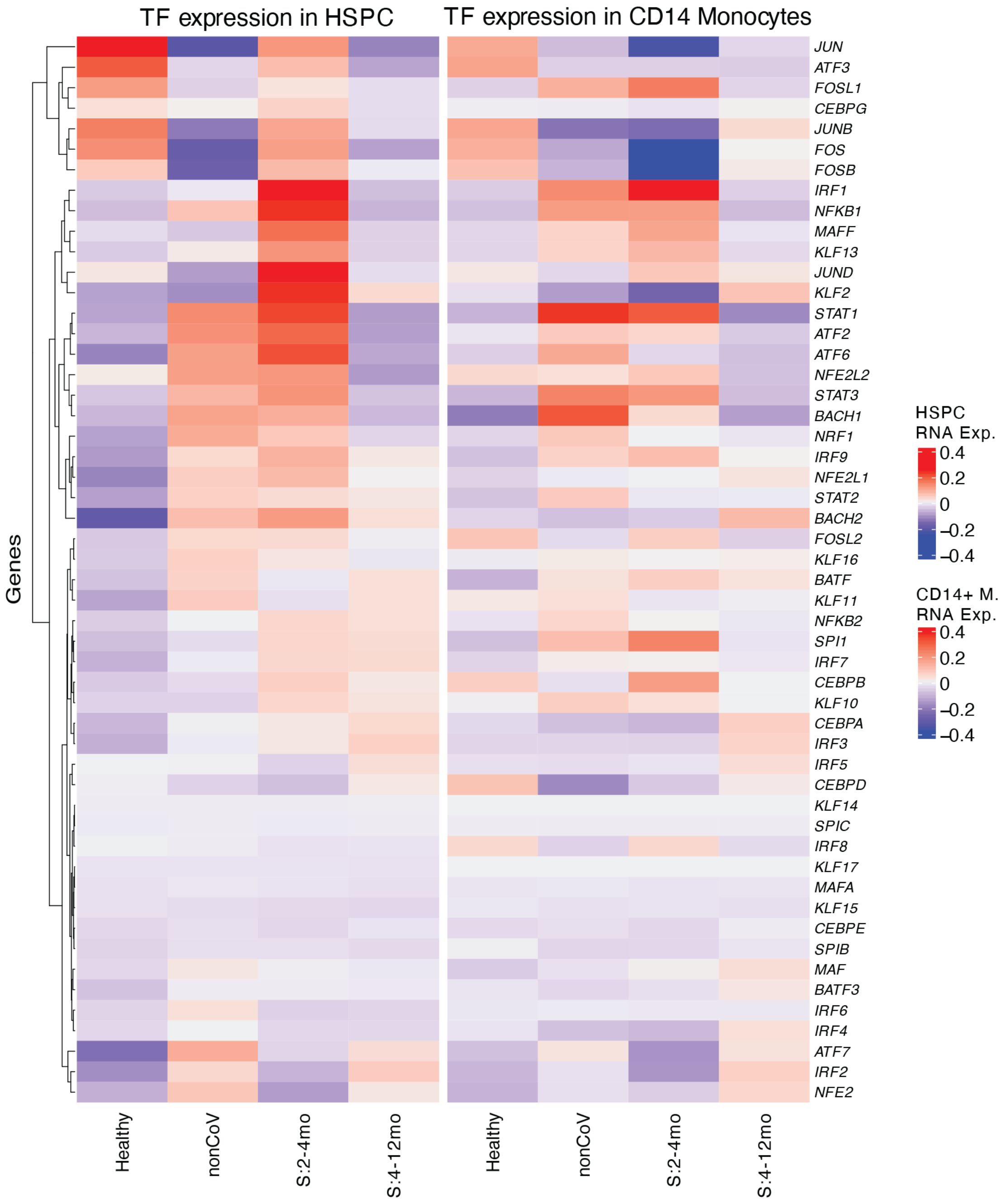


**Fig. S8. Transcription factor expressions in HSPC and CD14+ monocytes.**

Heatmap showing average expression of selected transcription factors in HSPC and CD14+ monocytes across clinical cohorts.

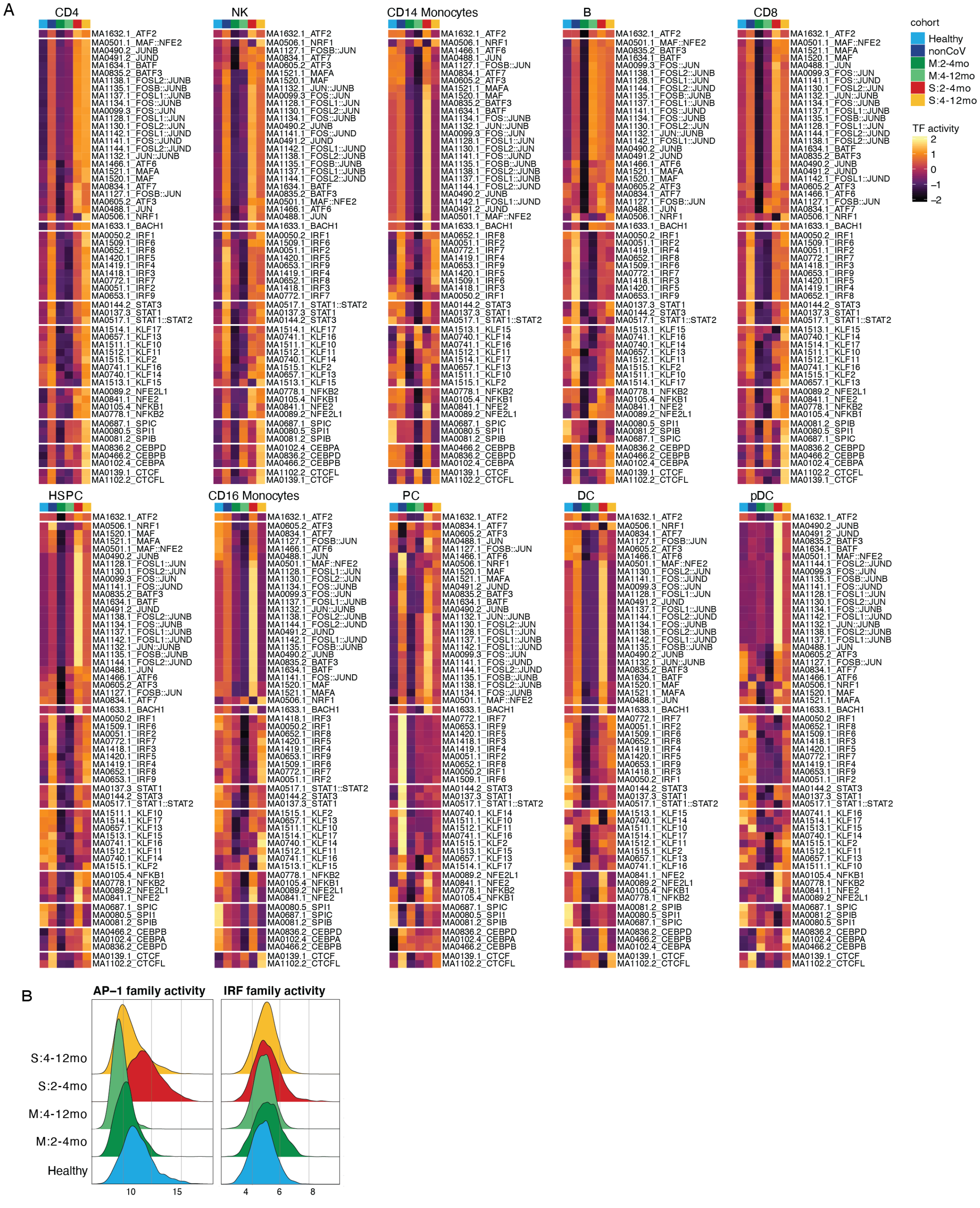


**Fig. S9. Transcription factor expression and TF activity related to Figure 5.**

**(A)** Heatmap showing motif activities of TF families across each clinical cohort in individual cell types. ChromVAR score was z-score-normalized by row after taking the median for each cohort. **(B)** Ridgeplot showing mean TF activity of IRF family and AP-1 family per cell grouped by cohorts.

### Cohort Demographic Information

| | | **Total (n=76)** | HD (n=25) | Mild: 2-4mo (n=8) | Mild: 4-12mo (n=2) | Severe: 2-4mo (n=18) | Severe: 4-12mo (n=12) | Non-COV (n=11) |
|---|---|---|---|---|---|---|---|---|
| Age (Mean ± SD) | | **55 ± 13.9** | 55 ± 15.4 | 45 ± 17.8 | 41 ± 1.4 | 55 ± 11 | 56 ± 12 | 63 ± 11.4 |
| Sex | Male | **53 (70%)** | 15 | 4 | 2 | 17 | 10 | 5 |
| | Female | **23 (30%)** | 10 | 4 | 0 | 1 | 2 | 6 |
| Race/Ethnicity | White | **35 (46%)** | 14 | 5 | 2 | 6 | 0 | 8 |
| | Black | **2 (3%)** | 1 | 0 | 0 | 1 | 0 | 0 |
| | Asian | **7 (9%)** | 4 | 0 | 0 | 2 | 1 | 0 |
| | Hispanic | **16 (21%)** | 2 | 1 | 0 | 6 | 6 | 1 |
| | Declined | **16 (21%)** | 4 | 2 | 0 | 3 | 5 | 2 |
| BMI (Mean ± SD)* | | **28 ± 5.7** | 25.6 ± 4.3 | 27.5 ± 6.1 | 28.2 ± 9.6 | 31.8 ± 6.2 | 29 ± 5 | 26.2 ± 6.1 |
| Comorbidities | Obesity* | **22 (31%)** | 3 | 2 | 1 | 10 | 3 | 3 |
| | Diabetes | **12 (15%)** | 1 | 1 | 0 | 6 | 3 | 1 |
| | Metabolic | **34 (44%)** | 6 | 2 | 0 | 14 | 5 | 7 |
| | Immune | **18 (23%)** | 8 | 2 | 0 | 3 | 3 | 2 |

HD - Healthy Donor, EC - Early Convalescence, LC - Late Convalescence
*Data not available for 7/78 Total, 3/25 HD, 3/8 Mild EC, and 1/18 Severe EC subjects

**Table S1. Cohort Demographic Information.**

**Data S1.**

Sample master chart

**Data S2.**

QC matrices of the single-cell dataset and bulk/pseudobulk ATAC-seq data used in this research.

**Data S3.**

Differentially expressed genes across cell types.
